# Heritable variance in fitness in wild vertebrates accounts for a small share of their missing neutral diversity and exceeds what recurrent mutation is estimated to maintain

**DOI:** 10.64898/2026.11.24.769021

**Authors:** Walid Mawass

**Affiliations:** Department of Human Genetics, University of Chicago, Chicago, IL, USA

## Abstract

Pedigree-based estimates show that wild populations carry substantial additive genetic variance in lifetime fitness. In classical theory, such variance reduces neutral diversity and is maintained by recurrent deleterious mutation. This variance is not expected to explain the missing genetic diversity of abundant species, but neither its share of that deficit nor the share maintained by mutation has been measured in individual natural populations. We measure both in 19 pedigreed populations of birds and mammals (18 for maintenance, 9 for diversity), allowing for extra-pair paternity and chromosome-scale linkage. If loci act multiplicatively on fitness, the measured variance roughly halves the median population’s effective size. On a logarithmic scale, it accounts for a median 7% (1–40%) of the observed 230- to 38,000-fold deficits; explaining them would require a long-term variance above one, larger than any estimate. At measured mutation rates and mean deleterious effects estimated in related species, mutation–selection balance maintains about a twentieth of the variance in the median population and less than all of it in 16 of 18 populations. This excess persists in 12 populations when a larger mutational target, undetected large-effect mutations, and an estimate inflated twofold are combined, and in 6 under stronger departures. Because the variance that mutation–selection balance can maintain removes at most 5% of neutral diversity at unlinked sites in populations with diversity data, no rescaled estimate fits both equilibria. With current estimates, most of the measured variance is maintained by other processes, and most of the missing diversity has other causes.

## 1 Introduction

The additive genetic variance in relative fitness, *V*_*A*_(*w*), is the heritable component of variation in reproductive success [1, 2], and pedigree-based animal models fitted to lifetime breeding success now estimate it for total fitness in wild populations [3, 4]. By Fisher’s fundamental theorem and the secondary theorem of selection [5, 6], a non-zero *V*_*A*_ implies an ongoing increase in mean genetic fitness and directional selection on traits genetically correlated with fitness, without specifying which alleles respond. Yet wild populations often change less than these theorems predict [7, 8]. Bonnet et al. [9] meta-analyzed pedigree estimates from 19 long-term studies of birds and mammals and found that *V*_*A*_ is often substantial, which they interpreted as evidence of contemporary adaptive evolution. Such a variance, however, is measured over a few generations, whereas its two population-genetic consequences are long-term. Heritable variance in fitness reduces neutral diversity, and selection depletes that variance unless some process maintains it. How much of these populations’ missing neutral diversity does the measured variance account for, and how much of that variance can recurrent mutation maintain?

The effective population size (*N*_*e*_) implied by a population’s neutral genetic diversity and mutation rate is smaller than its census size (*N*_*c*_), often by several orders of magnitude [10, 11]. This deficit grows with abundance, so that across animal species genetic diversity varies over a much narrower range than census size (Lewontin’s paradox [12, 13]). Heritable variance in fitness is one of the reasons *N*_*e*_ is below the census size, and theory predicts the reduction in *N*_*e*_ from that variance alone, whatever the genetic architecture [2, 14]. Random variation in family size causes drift on its own [15]. Heritable variance in fitness adds to that drift because an ancestor’s advantage passes to its descendants, which increases the variance among ancestors in long-term genetic contribution [2], the quantity that sets drift at a neutral site [16]. Theories of drift under selection differ in how they compute the long-term contribution, but all assume that selection remains constant over the few generations in which an ancestor’s advantage decays. Santiago and Caballero [14, 17] modeled the advantage as decaying by a constant fraction each generation; assuming Poisson non-heritable variance in family size, the equilibrium ratio of effective size to the number of breeding adults, *N*, at an unlinked locus is

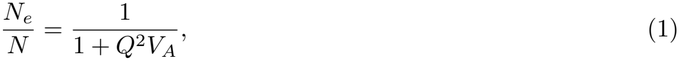

where *V*_*A*_ is the long-term equilibrium additive genetic variance in relative fitness and *Q*, Robertson’s cumulative transmission factor, sums over generations the fraction of an ancestor’s selective advantage that persists in its descendants. Woolliams et al. [18, 19] generalized this framework to multiple fitness components, including correlated reproductive success of mates.

Whereas these ancestor-centered models follow the descendants of an individual, Barton and Etheridge [20] followed a gene copy, which shares its carrier’s advantage only while it stays associated with the selected alleles the carrier transmits. That association is the linkage disequilibrium (LD) between a neutral site and the selected sites, which arises whether or not the sites are linked [21]. From this gene-centered perspective, Robertson’s *Q* is this LD summed over the generations before it decays. At an unlinked site, independent assortment halves the LD each generation, so *Q* = 2 [17, 22]. The two perspectives coincide there because LD and the share of an ancestor’s breeding value carried by its descendants are both halved each generation. Consequently, *N*_*e*_ is reduced by the same amount at every unlinked neutral site, and Equation (1) gives that reduction provided *V*_*A*_ is small. When loci act multiplicatively on fitness, the long-term contribution becomes exponential in the breeding value, and heritable variance in fitness multiplies *N*_*e*_ at unlinked sites by 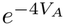 under random mating and by 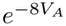 under strict monogamy [23]. At a linked site, by contrast, the association lasts until a rare crossover breaks it or selection removes the allele, so the reduction varies along the genome and, where that removal is slow, can be much stronger than the unlinked reduction [24, 25]. Studies of this variation, mostly under models of background selection, have inferred heritable variance in fitness or the strength of purifying selection from diversity along human chromosomes or from temporal allele-frequency covariances [26–29].

Because a linked association can last as long as the selected allele segregates, the reduction in *N*_*e*_ at linked sites depends on the mechanism that maintains *V*_*A*_. Mutation–selection–drift balance (MSDB) is the classical null model for this maintenance [30, 31]. When purifying selection is strong relative to drift, MSDB maintains an additive genetic variance in fitness of approximately

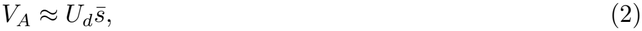

where *U*_*d*_ is the diploid genome-wide deleterious mutation rate and 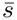 is the arithmetic mean of the distribution of deleterious fitness effects (DFE) [32, 33]. Measured variance in fitness, however, is often larger. Charlesworth [33] showed that standing genetic variance in *Drosophila* fitness components exceeds the MSDB expectation by more than an order of magnitude, which implies a contribution from balancing selection, fluctuating selection, or departures from equilibrium [34, 35]. Similarly, Connallon and Czuppon [36] argued qualitatively that vertebrate deleterious mutation rates are too low to sustain the estimates of Bonnet et al. [9], without testing those estimates against measured 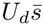. Because *U*_*d*_ and 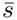 can now be estimated from comparative and population-genomic data [37–40], the share of each population’s *V*_*A*_ that recurrent mutation maintains can be measured.

Whatever maintains it, heritable variance in fitness reduces *N*_*e*_ at unlinked sites, yet studies of the diversity deficit have focused on linked effects. At MSDB, this unlinked reduction is the free-recombination limit of background selection, long considered negligible because unlinked associations decay within a few generations [22]. However, Matheson and Masel [41] showed that with vertebrate-like recombination maps, the unlinked reduction is larger than the linked one whenever *U*_*d*_ exceeds 1. The unlinked reduction can also be predicted from a pedigree estimate of *V*_*A*_ and the mating system, without a fine-scale recombination map and without assuming that *V*_*A*_ is maintained at MSDB. Charlesworth and Jensen [35] applied this prediction to the cross-species mean estimate of Bonnet et al. [9] and obtained a 16% to 55% reduction in neutral diversity. A reduction of that size is far smaller than the deficits of Lewontin’s paradox, so heritable variance in fitness is not expected to explain them. What has not been estimated is the size of the reduction in each population, given its own estimate of *V*_*A*_, mating system, and linkage map.

Both the diversity prediction and MSDB concern the long-term additive genetic variance in relative fitness, a parameter that a contemporary pedigree estimate [9] does not directly measure. Pedigree studies span a few decades of particular cohorts and environments, over which additive genetic variance and heritability can fluctuate [42–44]. Neutral diversity, by contrast, reflects an effective size averaged over the roughly 2*N*_*e*_ generations that pairs of lineages take to coalesce, and so depends on a *V*_*A*_ sustained over that span. Because MSDB describes a stationary variance, testing a population against MSDB also tests the stationarity on which equilibrium diversity predictions rely. By applying both tests to the same populations, we therefore ask whether a single stationary variance can satisfy both models, or whether departures from equilibrium are required to explain the heritable variance in fitness, the diversity deficit, or both.

Here we test the pedigree estimates of Bonnet et al. [9] against both equilibrium models, 18 of the 19 for maintenance at MSDB and the 9 with diversity and census data for neutral diversity, allowing for the mating system, including extra-pair paternity, and for linkage along chromosomes. Under multiplicative fitness effects, the predicted unlinked reduction roughly halves the effective size in the median population. Among the 9 populations with diversity data, it accounts for a median of 7% of the deficit on a logarithmic scale. Explaining the observed deficits, which range from 230- to 38,000-fold, would require a long-term *V*_*A*_ *>* 1, larger than any empirical estimate, and including linkage does not resolve the shortfall in any population. Recurrent deleterious mutation maintains only about 5% of the measured *V*_*A*_ in the median population. Because the long-term variances required by the two equilibria differ by more than a hundredfold in every population with diversity data, no single stationary variance satisfies both. That gap depends on each population’s mutational input, mating system, and observed deficit, and not on its pedigree estimate.

## 2 Theory and Model

### 2.1 The long-term additive genetic variance in fitness

The central quantity is the additive genetic variance in relative fitness, *V*_*A*_(*w*), written *V*_*A*_ hereafter, the variance of breeding values for lifetime reproductive success standardized to mean fitness (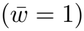. Its square root is the standard deviation of the relative fitness that individuals’ breeding values predict, so at *V*_*A*_ = 0.1 an individual one standard deviation above the mean expects about a third more offspring than average. Bonnet et al. [9] estimated *V*_*A*_ on this observed scale, by back-transforming the latent parameters of their animal model (subsection 3.1), so the estimates are already on the scale the theory uses.

For a single variance to satisfy both tests, its average over the roughly 2*N*_*e*_ generations that the genealogy spans must equal the stationary variance of MSDB (subsection 2.2). A pedigree estimate instead describes the cohorts and environment of a single study. Its back-transformation also depends on the state of the population during the study, such as the proportion of individuals that never breed, so the same latent genetic variance would give a different *V*_*A*_ in another environment. We therefore write the required long-term variance as

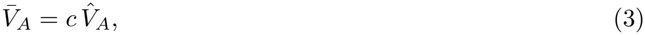

where 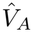 is the pedigree estimate and *c* is an unknown scaling factor, which measures how representative the study is of the population and period that neutral diversity records. Because the lineages sampled for diversity move among localities, whereas a pedigree samples one, *c* includes spatial as well as temporal departures from the long-term variance. Table S1 lists all notation.

### 2.2 The variance maintained at mutation–selection–drift balance

At MSDB, deleterious mutations contribute additive genetic variance in fitness at a rate *V*_*m*_ = *U*_*d*_ E[*s*^2^] per generation, where *U*_*d*_, the diploid genome-wide deleterious mutation rate, is the mean number of new deleterious mutations in a zygote, and *s* is the heterozygous effect on relative fitness. Under linkage equilibrium, the standing variance is 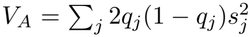 over sites segregating for a semidominant deleterious allele at frequency *q*_*j*_, and each generation selection removes a fraction *s* of this variance and drift a fraction 1*/*(2*N*_*e*_) [33, 45]. For a single effect *s*, the diffusion solution for a semidominant allele [46] (Appendix A.1) gives the equilibrium variance

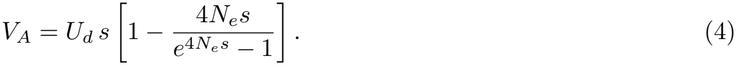

When selection is strong relative to drift, each deleterious allele is held near its deterministic mutation– selection balance frequency, yielding [22, 32]

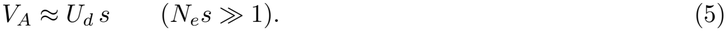

Intuitively, a mutation adds variance in proportion to *s*^2^ for the roughly 1*/s* generations before selection removes it, so it contributes in proportion to *s*, and only the number of new mutations and their mean effect matter. Because the bracketed term in Equation (4) never exceeds one, *U*_*d*_ *s* is the most variance MSDB can maintain at any effective size. Inverting Equation (5) yields the deleterious effect required to sustain a given long-term variance,

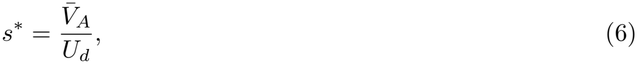

which we evaluate at the pedigree estimate (*c* = 1).

Under a distribution of fitness effects (DFE), a deleterious class *k*, with heterozygous effect *s*_*k*_, receives a fraction *φ*_*k*_ of new mutations and, provided *N*_*e*_*s*_*k*_ *≫* 1, contributes *U*_*d*_*φ*_*k*_*s*_*k*_ to the standing variance. The classes together maintain 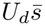, where 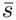 is the arithmetic mean of the DFE, so *s*^*\**^ estimates that mean whatever the shape of the DFE. Nearly neutral mutations (*N*_*e*_*s*_*k*_ ≲ 1) contribute less (Appendix A.3). We assume this strong-selection limit throughout, which reduces MSDB to mutation–selection balance (Appendix A.2 treats weak selection).

The variance *V*_*A*_ describes selection at two levels. At the level of the phenotype, it measures directional selection, which raises the mean breeding value for fitness by *V*_*A*_ each generation (Fisher’s fundamental theorem). At the level of alleles, selection at MSDB is purifying, whether it is unconditional or arises from stabilizing selection on rare variants. The additive variance in fitness, the per-generation gain in mean fitness it produces, and the mutational loss that offsets that gain therefore all equal 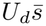 (with 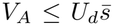 when stabilizing selection permits common alleles). We use *directional* only for phenotypes, and *purifying* and *positive* for selection against and for an allele.

### 2.3 Effective size at unlinked sites

Consider a diploid population of a constant number *N* of breeding adults, with an equal sex ratio and pairing at random with respect to breeding value (Appendices B and C give general forms). A neutral site in this population assorts independently of all selected loci, and we suppose for now that only segregation erodes its association with its selected background. We follow a neutral gene copy from an ancestor (*t* = 0) whose breeding value for log-fitness deviates from the population mean by 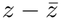. Because each subsequent carrier receives half of its breeding value from the parent transmitting the copy and half from a randomly drawn mate, the carrier’s expected deviation halves each generation, to 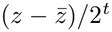 in generation *t*. For small *V*_*A*_, the carrier’s expected relative fitness is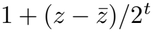. Because the expected number of copies is multiplied by the carrier’s relative fitness each generation, the ancestor’s expected long-term contribution is

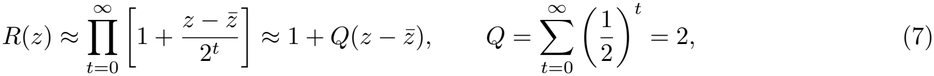

which gives Robertson’s cumulative transmission factor (Equation (1)). *R* is the share of the population’s genes, many generations later, that descends from this ancestor, relative to the share of an average ancestor.

The pedigree estimate 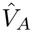 is the variance of the *t* = 0 factor alone, the ancestor’s own relative fitness, and the later terms follow from Mendelian inheritance. Intuitively, *Q* = 2 means that an ancestor’s descendants double its initial heritable advantage, provided selection remains roughly constant over the few generations in which that advantage decays.

Drift at a neutral site is set by the variance in long-term contributions among ancestors [16]. We express the reduction that heritable variance in fitness causes as *N*_*e*_*/N*, the effective size relative to that of a neutral population of *N* breeding adults with Poisson-distributed family sizes, in which *N*_*e*_ = *N* . In terms of the expected long-term contributions of all the ancestors in a generation, this ratio is

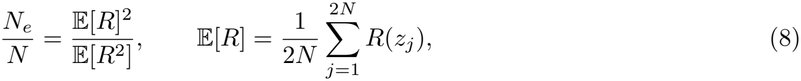

where ancestral gene copy *j* is carried by an ancestor with breeding value *z*_*j*_ and E[*R*^2^] averages *R*(*z*_*j*_)^2^ in the same way. The ratio follows from the pairwise coalescence probability (Appendix B.1). Two lineages are more likely to meet in an ancestor that left a large share of descendants, so the more unequal the shares, the faster lineages coalesce. Every reduction at unlinked sites that we report is computed from this ratio. The theories of drift under selection that we use all compute *R* and differ in how it depends on the breeding values of an ancestor and its mates (Table S2). For small *V*_*A*_, the linear approximation 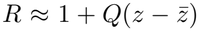 gives *Q*^2^*V*_*A*_, the variance in long-term contributions that inherited fitness differences add to the Poisson baseline [2, 18, 47] (Equation 16 of [14]; Appendix B.2). For an unlinked site under random mating (*Q* = 2), Equation (1) therefore simplifies to *N*_*e*_*/N* = 1*/*(1 + 4*V*_*A*_), which we call the *small-variance approximation*.

Because Equation (8) comes from the asymptotic rate at which two lineages coalesce, *N*_*e*_ is the asymptotic inbreeding effective size. In the Kingman coalescent limit, this size is the coalescent effective size *N*_*eC*_ that neutral diversity measures, *π* = 4*N*_*eC*_ *μ*, where *μ*is the per-site mutation rate (Appendix B.1 relates it to the variance effective size).

In the gene-centered perspective of Barton and Etheridge [20], 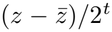 is the linkage disequilibrium (LD) between the neutral copy and its carrier’s selected background, and the ancestor-centered and gene-centered perspectives coincide at an unlinked site. Only the gene-centered perspective extends to the other processes that break the association, selection (next paragraph) and recombination (subsubsection 2.6.2).

Selection further erodes an ancestor’s advantage by removing carriers of deleterious alleles. The retention factor *Z* is the fraction of a neutral site’s association with its selected background that survives one generation of selection, which Santiago and Caballero [14] treat as constant across generations. With mates’ breeding values correlated by *r*_*mf*_, Santiago and Caballero’s cumulative transmission factor is

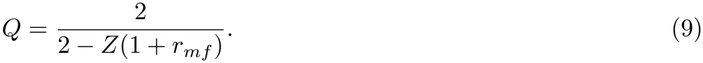

For a Gaussian breeding value composed of many loci of small effect under multiplicative fitness, *Z* = 1 exactly. Because each unit of breeding value multiplies fitness by the same factor everywhere in the distribution, selection shifts the mean without compressing the variance among parents, so only segregation erodes a lineage’s relative advantage (Appendix B.4). The association with a deleterious allele, by contrast, is lost with the carriers that selection removes [22, 25], so under strong selection the association with an allele of effect class *k* is retained at

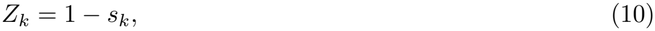

regardless of *V*_*A*_, and an unlinked site under random mating has *Q* = 2*/*(1 + *s*_*k*_) *<* 2 (Appendix A.4). Because purging erodes the association by a fraction *s*_*k*_ each generation whereas independent assortment halves it, purging changes *Q* little for small *s*_*k*_. Under MSDB, *Q* = 2 is therefore the largest value that *Q* takes, and the process maintaining *V*_*A*_ affects the unlinked reduction only through *Z*. We compute the unlinked reduction at *Z* = 1, which is biased toward passing the diversity test, and reserve *Z*_*k*_ for linked sites (subsubsection 2.6.2).

Because this reduction depends on the individual that carries the site and not on the haplotype on which the site lies, it is the same at every independently assorting neutral site, and linkage adds to it at linked sites (subsubsection 2.6.2). Under MSDB, it converges to the limit of background selection at unlinked loci (Appendix A.5). Like *N*, the factors *Q* and *Z* refer to breeding adults (subsection 3.1).

This model of the reduction in *N*_*e*_ requires fixed additive effects, so that an allele’s effect on fitness does not depend on its own frequency, which excludes underdominance, stabilizing selection, and frequency-dependent selection. It also requires that an ancestor’s advantage keep its sign over the few generations in which its contribution is settled. This is satisfied under MSDB, sweeps of positively selected alleles, and positively autocorrelated fluctuating selection (Appendix B.1). Additionally, the retention factor must remain constant across generations [14], and allele frequencies at the selected loci must change little while LD persists [21]. The infinitesimal model and MSDB at unlinked sites meet this last condition, whereas alleles that sweep between extreme frequencies violate it and generate more drift than the concurrent variance in fitness predicts.

These conditions concern only the few generations over which an ancestor’s contribution is settled, not the roughly 2*N*_*e*_ generations of the genealogy, over which *V*_*A*_ may change. Because each generation contributes an inverse effective size *N/N*_*e*_(*t*) = 1 + *Q*^2^*V*_*A*_(*t*), long-term diversity reflects the harmonic mean of *N*_*e*_ [15], which over *T* generations is

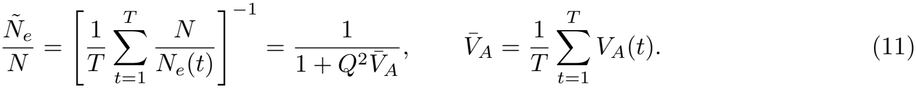

Under the small-variance approximation and constant *Q*, neutral diversity therefore depends only on the arithmetic mean 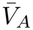 of Equation (3), not on how *V*_*A*_ fluctuates over time.

### 2.4 Large variances and the multiplicative bound

The small-variance approximation and its extension to other mating systems (subsubsection 2.6.1) require *V*_*A*_ *≪* 1, because they keep only the terms of *R*(*z*) that are linear in the breeding value. The per-generation advantages then add rather than multiply, and the products of advantages from different generations, which grow with *V*_*A*_, are dropped. Most pedigree estimates meet this condition. However, inverting the small-variance approximation for the observed deficits in neutral diversity (*N*_*eC*_ */N ≪* 1) gives 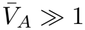, a range in which the approximation overstates the variance required.

Large variances therefore require a non-linear model of fitness, and the generalized linear mixed models of Bonnet et al. [9] already assume one, with expected breeding success exponential in a Gaussian latent breeding value *z* (*w ∝ e*^*z*^). Under a log link, the additive variance in relative fitness on the observed scale equals the latent variance, 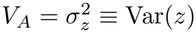, because the average derivative of expected fitness with respect to *z* is mean fitness [48, Equations 15–16], whereas the total genetic variance in relative fitness, 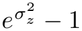, is larger by a part that is non-additive on the observed scale. The pedigree estimate 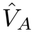 is the variance of *z* that the multiplicative model requires, exactly for the log-link component of the animal model and approximately once its zero-inflation component is included.

Barton and Etheridge [20] solved this multiplicative model exactly for parents drawn in proportion to *e*^*z*^, with *z* additive and Gaussian. Because a lineage’s per-generation selective advantages sum exactly on the log scale without truncation, the long-term contribution *R* is exponential rather than linear in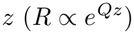, with *Q* = 2 in the exponent at unlinked sites, and Equation (8) evaluates to exp[*−*Var(ln *R*)]. Under random mating, heritable variance in fitness multiplies the coalescence rate by 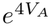[20, 23] *(Appendix B*.*1) and*

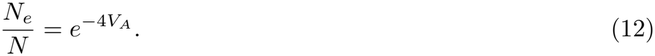

In the limit of small *V*_*A*_, both Equation (12) and the small-variance approximation converge to 1 *−* 4*V*_*A*_, but as *V*_*A*_ increases, exponential compounding produces a much stronger reduction in *N*_*e*_. The forms we use are nested rather than competing. The small-variance approximation is the limit for small *V*_*A*_ of every continuous mapping from breeding value to fitness, whereas the *multiplicative model* is exact for the mapping above, which the animal model assumes. Where the multiplicative model and its limit disagree, the difference measures how much a prediction depends on how loci combine to determine fitness. Under strict monogamy, where pairs form at random and family productivity scales with 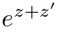 for partners with breeding values *z* and *z*′, the exponent doubles and the reduction becomes 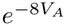 [23]. We write both reductions as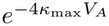, where *κ*_max_, the variance of long-term log-contributions in units of its random-mating value, is 2 under social monogamy and 1 under random mating; modeling extra-pair paternity gives a smaller reduction (subsubsection 2.6.1).

The exponential reduction depends on the unbounded upper tail of the Gaussian distribution for log-fitness. A 100-fold reduction in *N*_*e*_, for instance, comes mostly from ancestors more than four standard deviations above the mean (Appendix B.3). In natural populations, ecological and physiological constraints, such as territorial saturation, clutch-size trade-offs, or the bounded probability of breeding in Bonnet et al.’s zero-inflation component, truncate this tail and weaken the resulting drift. We therefore treat 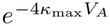 as the largest reduction in neutral diversity that a stationary variance *V*_*A*_ can cause, and call it the *multiplicative bound*. The coalescence rate is linear in *V*_*A*_(*t*) under Equation (11) but convex under the bound 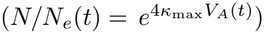. By Jensen’s inequality, a variance that fluctuates around a mean 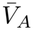 removes more diversity than a constant variance with that mean. Consequently, the minimum variance that accounts for a neutral diversity deficit (subsection 2.5) is a lower limit on the constant variance that would produce the same average coalescence rate, (4*κ*_max_)^*−*1^ ln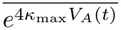, where the bar denotes the average over generations, and not on 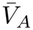 itself. This constant variance exceeds 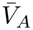 appreciably only if *V*_*A*_ reaches, in some generations, values well above its average. A population of *N* breeding adults, moreover, realizes the bound only down to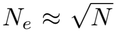, because beyond that point the ancestors that produce the reduction are rarer than one per generation (Appendix B.3).

### 2.5 The two equilibrium tests as limits on the long-term variance

Let

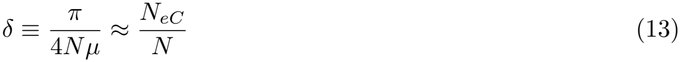

denote the diversity ratio, the fraction of its neutral expectation that a population’s diversity retains. Our *N*_*e*_*/N* predicts *δ* only if heritable variance in fitness is the sole departure from the Poisson baseline, the population is at long-term equilibrium, and *N* is the harmonic mean, over the genealogy (Equation (11)), of the number of breeding adults in the population that *π* samples. We distinguish this parameter from its estimate, 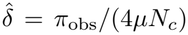, in which *π*_obs_ is the observed diversity and *N*_*c*_, a contemporary census of all adults, stands in for *N* (subsection 3.8). We call 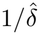 the *deficit*, so a 1,000-fold deficit leaves a thousandth of the expected diversity. The diversity ratio is usually orders of magnitude smaller [10, 11] than the demographic ratio *N*_*eV*_ */N* of a single generation, which compounds all sources of reproductive variance and is typically 0.1–0.5 [49]. Inverting the multiplicative bound at *δ* gives the minimum long-term variance that accounts for the deficit:

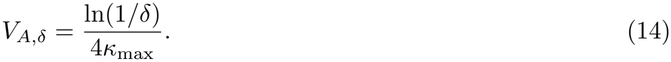

Because the multiplicative bound maximizes the reduction in *N*_*e*_ per unit of variance, a population with 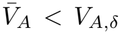 cannot owe its diversity deficit solely to the effect of heritable variance in fitness at unlinked sites. The variance *V*_*A,δ*_ grows only logarithmically with the deficit, by ln(10)*/*(4*κ*_max_) *≈* 0.58*/κ*_max_ for each further tenfold loss of diversity, so under random mating a 100-fold deficit requires 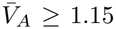 and a 10,000-fold deficit 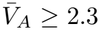 (Figure S1). Inverting the small-variance approximation instead gives a ceiling that holds only in populations retaining at least about a fifth of their neutral expectation under random mating and a ninth under strict monogamy (Appendix B.2).

In terms of 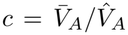 (Equation (3)), each equilibrium sets a one-sided threshold, the limit on how far the estimate can be rescaled before its test fails. We call the calculation of both thresholds for each population the threshold analysis. Because MSDB maintains at most 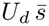 (subsection 2.2), a population passes the maintenance test only if

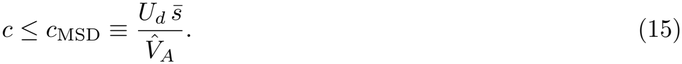

When *c*_MSD_ *<* 1, this ratio is the largest fraction of the pedigree estimate that recurrent deleterious mutation can sustain. On the diversity side, heritable variance in fitness accounts for the deficit through its effect at unlinked sites only if 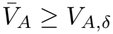, so a population passes the diversity test only if

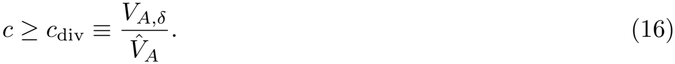

When *c*_div_ *>* 1, *c*_div_ is the smallest factor by which the long-term variance must exceed the pedigree estimate to explain the missing diversity. Its reciprocal, 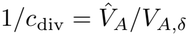, is the share of the deficit that 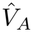 accounts for under the multiplicative bound. We measure this share on a logarithmic scale, as the logarithm of the predicted reduction, 1*/*(*N*_*e*_*/N* )_pred_, divided by that of the deficit, 1*/δ*, because successive reductions in effective size multiply. Because both thresholds are conditions for equilibrium, a population that requires *c* ≠ 1 departs from stationarity. Table 1 summarizes how the two verdicts combine when evaluated at the pedigree estimate (*c* = 1).

**Table 1:**
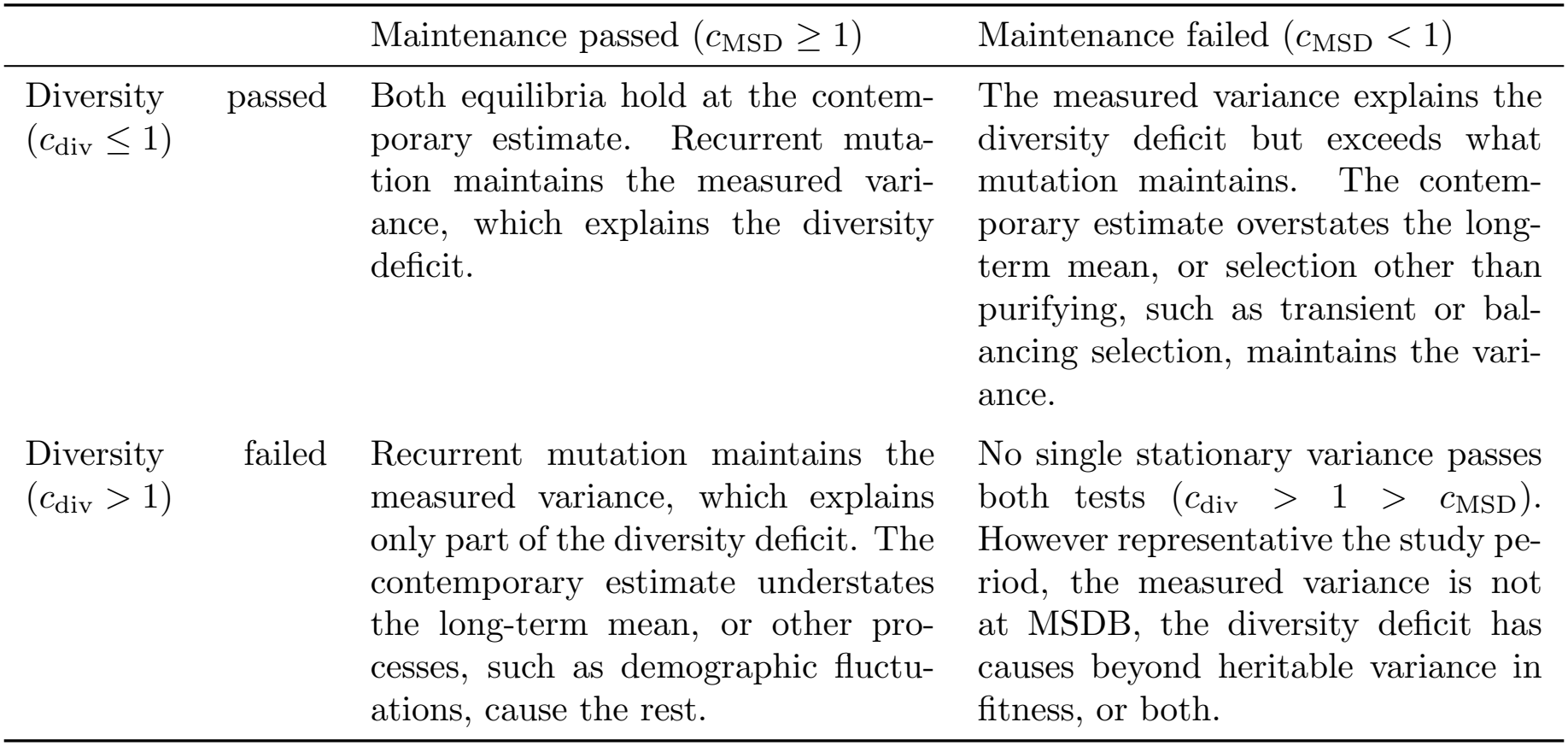
Whether the pedigree estimate must be scaled up or down shows which equilibrium assumption fails. The scaling factor is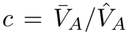. The diversity test is satisfied for *c ≥ c*_div_ and the maintenance test for *c ≤ c*_MSD_; rows and columns classify the combined verdict at the contemporary pedigree estimate (*c* = 1). Both thresholds are derived under approximations biased toward passing (subsection 2.7).

|  | Maintenance passed (*c*<sub>MSD</sub> ≥ 1) | Maintenance failed (*c*<sub>MSD</sub> < 1) |
|---|---|---|
| Diversity passed (*c*<sub>div</sub> ≤ 1) | Both equilibria hold at the contem-porary estimate. Recurrent muta-tion maintains the measured vari-ance, which explains the diversity deficit. | The measured variance explains the diversity deficit but exceeds what mutation maintains. The contem-porary estimate overstates the long-term mean, or selection other than purifying, such as transient or bal-ancing selection, maintains the vari-ance. |
| Diversity failed (*c*<sub>div</sub> > 1) | Recurrent mutation maintains the measured variance, which explains only part of the diversity deficit. The contemporary estimate understates the long-term mean, or other pro-cesses, such as demographic fluctu-ations, cause the rest. | No single stationary variance passes both tests (*c*<sub>div</sub> > 1 > *c*<sub>MSD</sub>). However representative the study pe-riod, the measured variance is not at MSDB, the diversity deficit has causes beyond heritable variance in fitness, or both. |

A single scaling factor *c* satisfies both tests if and only if *c*_div_ *≤ c*_MSD_. We call this condition, that both equilibria hold at a single long-term variance, the compatibility test, which a population passes only if

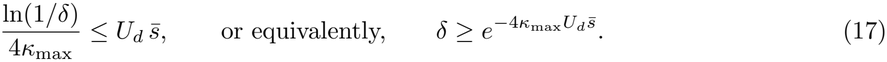

The compatibility test (Condition (17)) does not involve the pedigree estimate. Because 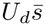 is the mean fitness lost to new deleterious mutations each generation, the bound 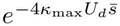 shows that the mutational loss of fitness in a single generation sets the largest loss of diversity that MSDB can cause at unlinked sites. At a typical vertebrate value of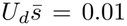, MSDB reduces neutral diversity at unlinked sites to no less than 0.96 of its neutral expectation under random mating (a 4% loss). Explaining even a 100-fold deficit (*δ* = 0.01) would require 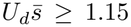 under random mating, a mutational loss larger than mean fitness itself in every generation. The condition therefore restates, for a population with a measured mutation rate, DFE, and diversity, the known result that variance maintained by recurrent mutation removes little neutral diversity at unlinked sites [22]. The pedigree estimate is used only in the threshold analysis, where it gives each population’s position relative to the two thresholds and, from that position, the two shares. Even a long-term variance smaller than 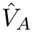 by the factor *c*_MSD_, and so small enough to be maintained at MSDB, would leave almost all of the diversity deficit unexplained.

### 2.6 Extensions

#### 2.6.1 Mating system

Under social monogamy the nest is the unit of productivity, so a parent’s family size depends on its social partner’s breeding value as well as its own, and extra-pair paternity reassigns a fraction *α* of a nest’s offspring to outside sires without changing how many the nest raises. For small *V*_*A*_ the reduction becomes (Appendix C.1)

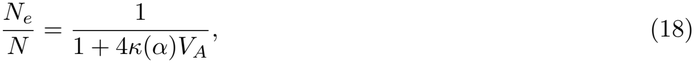

where *V*_*A*_ is the variance common to both sexes that Bonnet et al. estimate, and the mating-system scalar *κ*(*α*), the variance in long-term contributions averaged over the sexes in units of its random-mating value 4*V*_*A*_, is the factor by which the mating system amplifies the drift that a given *V*_*A*_ causes (*κ* = 1 under random mating). Without extra-pair paternity (*α* = 0), social partners share every offspring, and with it each other’s advantage, which doubles the variance of long-term contributions (*κ* = 2, Santiago and Caballero’s result for pair-mated families). As extra-pair paternity increases, a male’s reproductive success depends less on his social partner’s breeding value and more on his own, so when his extra-pair success is proportional to his breeding value (Model B in Appendix C), *κ* decreases monotonically to 9*/*8 at *α* = 1, still above the random-mating value because a female’s family size still depends on her social partner (Appendix C.2). Under the multiplicative model the reduction lies between 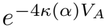 and 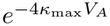 (Appendix C.4). The multiplicative bound takes the stronger of the two, 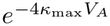, with *κ*_max_ = 2 whatever the rate of extra-pair paternity, so the mating system changes the variance that a given deficit requires at most twofold.

#### 2.6.2 Linkage

Recombination separates a neutral site from linked selected loci less often than independent assortment separates unlinked sites. Unless purging removes the deleterious allele first, the association therefore lasts longer than at an unlinked site and reduces *N*_*e*_ further, by an amount that varies along the genome. This reduction at linked sites requires two inputs that a pedigree does not supply, each chromosome’s share of the variance and how long its associations persist, which standard models of background selection derive from MSDB [28, 29, 50]. Here, by contrast, the pedigree estimate supplies the total variance and the genome annotation sets each chromosome’s share, 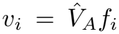, where *f*_*i*_ is the proportion of functional sequence on chromosome *i*. Only persistence requires MSDB, through the retention factor *Z*_*k*_ of each effect class (Equation (10)) weighted by its equilibrium share of *V*_*A*_, 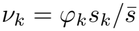 (Appendix B.4). The linked term is a hybrid in our analysis, in which the measured variance sets the size of the reduction at linked sites and the DFE sets how fast the associations decay, so it predicts a larger reduction than MSDB whenever 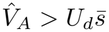 and recovers the MSDB prediction at 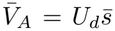. The expressions below are for a single effect class with retention factor *Z*.

We treat the chromosome as the unit of the linked reduction, because pedigree linkage maps give its genetic length reliably but resolve its fine-scale recombination landscape poorly. We place the neutral site and selected loci uniformly along chromosome *i*, of genetic length *M*_*i*_ Morgans, averaging the reduction over all pairwise positions. LD between a neutral site and a selected locus *ℓ* Morgans apart is retained across a generation of recombination and selection in the proportion *ρ* = (1 *−r*(*ℓ*))*Z*, where *r*(*ℓ*) is the recombination fraction under Haldane’s mapping function. Robertson’s *Q* is the sum of these retained proportions over the generations that follow, Σ _*t≥*0_ *ρ*^*t*^ = 1*/*(1 *− ρ*). At an unlinked site 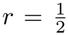, so *ρ* = *Z/*2 and the sum is Equation (9) with *r*_*mf*_ = 0, which equals 2 at *Z* = 1. At a linked site *ρ* is larger, so the same sum 1*/*(1 *− ρ*) exceeds the unlinked *Q* by an amount that depends on the distance *ℓ*. This factor is the expected number of generations that a neutral copy stays associated with a selected allele *ℓ* Morgans away, until recombination separates them or selection removes the allele. Because it varies with position, a chromosome has no single *Q*, and the reduction depends instead on its square averaged over the positions of the neutral site and the selected alleles on chromosome *i*, 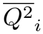 (the average of the square, not the square of an average), which has a closed form under Haldane’s map (Equation (D4), Appendix D).

At linked sites the ancestor-centered and gene-centered perspectives part, because a neutral copy stays associated only with the selected alleles on its own haplotype, which carries half of the chromosome’s variance (*v*_*h*_ = *v*_*i*_*/*2), whereas the homologous haplotype, the social partner, and the other chromosomes assort independently of it and contribute through the unlinked term. When fitness is multiplicative across loci (subsection 2.4), the two contributions combine additively in the exponent (Appendix C.3):

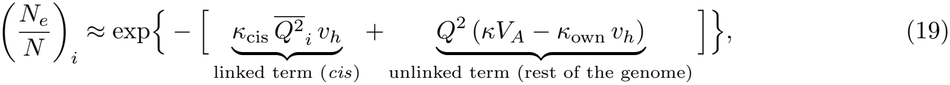

where *κ*_cis_ and *κ*_own_ weight the haplotype’s variance by the carrier’s own breeding value (both 1 under random mating and Model B; see Appendix C.3), and *Q* = 2*/*(2 *− Z*) evaluates Equation (9) at the same retention factor as the linked term. Dividing Equation (19) by the unlinked reduction at the same retention factor gives the linked factor, the additional reduction in *N*_*e*_ that linkage on chromosome *i* causes:

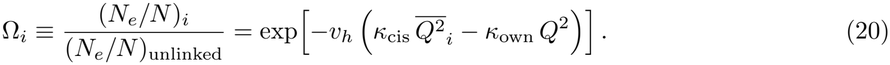

Intuitively, Ω_*i*_ is the share of the diversity predicted at unlinked sites that a neutral site on chromosome *i* keeps once linkage is added. It cannot exceed one, because linkage can only lengthen an association 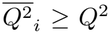 for every effect class) and the carrier’s breeding value weights the haplotype’s variance equally at linked and unlinked sites (*κ*_cis_ = *κ*_own_; Appendix D.5). With neutral sites uniform across the physical genome, the genome-wide reduction is then the average over the *n* autosomes weighted by their physical lengths *L*_*i*_, which sum to *L*_total_:

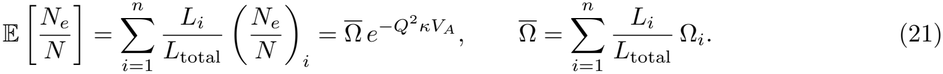

### 2.7 Approximations biased toward passing both tests

We chose approximations biased toward passing each test, so neither a failed test nor the result that no single variance passes both (*c*_div_ *>* 1 *> c*_MSD_) is a consequence of them. On the diversity side, six choices each maximize the reduction in effective size that a given *V*_*A*_ causes, and so minimize *V*_*A,δ*_: unbounded multiplicative fitness, the largest mating-system scalar *κ*_max_, complete retention (*Z* = 1) at unlinked sites, the transmission factor of fecundity applied to pre-breeding survival, a map without crossover interference, and the full 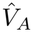 in the linked term. On the maintenance side, the strong-selection limit and a mutational target in which every mutation, synonymous ones included, is deleterious maximize the variance that recurrent mutation maintains, and so maximize *c*_MSD_ (Supplementary Methods justify each approximation).

Four factors could act in the opposite direction. First, if breeding values are strongly right-skewed rather than Gaussian, so that the variance in fitness is concentrated in a few extreme lineages, selection increases their variance among parents and *Z* can exceed one, which produces more drift than the Gaussian model predicts (Appendix B.4). Second, the linked model places selected and neutral sites uniformly along each chromosome, whereas clustering of functional loci in low-recombination regions strengthens linked selection (Appendix D.6). Third, under the multiplicative bound a fluctuating variance removes more diversity than its mean (subsection 2.4). Finally, an underestimate of *U*_*d*_ would raise 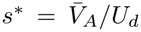, so we evaluate all maintenance tests across two mutation-rate estimates and a range of functional target sizes derived from the genome annotations (Methods).

## 3 Materials and Methods

### 3.1 Pedigree estimates of additive genetic variance in fitness

Posterior distributions of the additive genetic variance in relative fitness, *V*_*A*_, were obtained from Bonnet et al. [9] for 19 longitudinally monitored wild vertebrate populations: 11 socially monogamous populations (10 bird populations and the meerkat) and 8 polygynous or polygynandrous mammal populations. Bonnet et al. fitted a zero-inflated overdispersed Poisson animal model [3, 51, 52] to lifetime breeding success, including both sexes in each dataset with sex modeled as a fixed effect. We applied no standardization to these estimates because they were already back-transformed to the observed scale of relative fitness via the multivariate delta method, *V*_*A*_ = **d**^*⊤*^**Gd**, where **G** is the additive genetic covariance matrix of the latent predictors and **d** is the vector of average derivatives of mean relative fitness with respect to them. For each population, we used the 4,000 posterior draws of 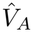 from these model fits to compute the posterior distribution of each derived quantity, summarizing results by the posterior mode and the 95% highest posterior density (HPD) interval. Across the 19 populations, the posterior modes of 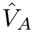 range from 0.003 to 0.50 (median 0.10).

Individuals with zero lifetime breeding success make up 53% (yellow baboons, Amboseli) to 96% (blue tits, Muro) of the records of Bonnet et al. (their Data S1, excluding missing lifetime breeding success), so 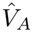 includes heritable variance in pre-breeding survival as well as fecundity (subsection 2.3). Offspring were counted when first marked, from shortly after birth in mammals to ringing or fledging in birds [9], so each individual’s survival to breeding age counts toward its own lifetime breeding success rather than its parent’s. Robertson’s *Q* sums an inherited advantage over the generations in which it acts. An advantage in fecundity acts first in the ancestor’s own reproduction (*t* = 0) and then, halved each generation, in that of its descendants, so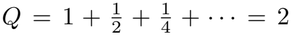 . An advantage in survival acts one stage earlier, before reproduction, and the ancestors whose contributions set drift are breeding adults that have already survived. A survival advantage therefore changes nothing at *t* = 0 and first acts on whether the ancestor’s offspring, which inherit half of it, survive to breed. The sum starts one generation late, 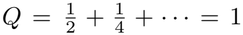, so, per unit of *V*_*A*_, survival to breeding age adds a quarter as much as fecundity to the variance of long-term contributions (*Q*^2^ = 1 rather than 4) [2, 14]. Because the data do not separate these two components of fitness, we applied *Q* = 2 to all of 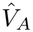, which overstates the predicted reduction in *N*_*e*_ unless pre-breeding survival and adult fecundity are negatively genetically correlated (subsection 2.7).

### 3.2 Mating system and the reduction at unlinked sites

We classified populations as socially monogamous when offspring are reared by a social pair (or by a dominant breeding pair in the cooperatively breeding superb fairy-wrens and meerkats) and as randomly mating otherwise. For each socially monogamous population, we obtained the extra-pair paternity rate *α*, the proportion of offspring not sired by the social male, from published molecular parentage studies of the same population (Table S3). For superb fairy-wrens, offspring sired by within-group subordinate helpers were counted alongside extra-group paternity as not sired by the dominant male; for meerkats, *α* was the proportion of the dominant female’s pups not sired by the dominant male. For the Karori hihi and the Hoge Veluwe great tits, which lack parentage estimates, we used the estimates from the Tiritiri Matangi source population and from the neighboring Westerheide population, respectively (Westerheide *α* = 0.177; earlier estimates 0.065–0.125 [53]). Across the 11 socially monogamous populations, *α* ranges from 0.097 (meerkats) to 0.61 (superb fairy-wrens).

We set *κ* = 1 under random mating and evaluated *κ*(*α*) under social monogamy with Equation (C6) (Model B of Appendix C.1), giving values from 1.32 to 1.86. We computed the reduction in *N*_*e*_ at un-linked sites (subsection 2.3) at *Z* = 1 and with no correlation between mates’ breeding values for fitness, because no estimate of that correlation exists. For each population, we evaluated the multiplicative model from the moment equation for long-term contributions (Equation (C11), Appendix C.4); the multiplicative bound using *κ*_max_ (Equation (C12)); and the small-variance approximation, the limit of both for small *V*_*A*_ (Equation (18)).

### 3.3 Mutational target and deleterious mutation rate

We downloaded genome assemblies and annotations from NCBI Datasets (accessions in Table S3). Sixteen populations have an annotated chromosome-level assembly, representing their own species in all cases except the yellow baboons and Soay sheep, for which we used the olive baboon (*Papio anubis*) and domestic sheep (*Ovis aries*, Rambouillet) reference assemblies, respectively. The spotted hyena and hihi assemblies are annotated only at the scaffold level, with the hihi annotation taken from the B10K consortium [54]. Because the sole available superb fairy-wren assembly [55] is chromosome-level but unannotated, we applied the maintenance test to the remaining 18 populations. For each assembled autosome *i*, we recorded the physical length *L*_*i*_, the number of annotated protein-coding genes, and the coding sequence length *L*_coding,*i*_, the union of all annotated CDS features, so that isoforms and overlapping reading frames were counted once. Sex chromosomes, mitochondrial sequences, and unplaced scaffolds were excluded. For the scaffold-level spotted hyena and hihi assemblies, we scaled the total genome-wide CDS length by the autosomal CDS fraction of their nearest chromosome-level relatives: 0.965 for the hyena (from the meerkat) and 0.948 for the hihi (the mean across the collared flycatcher, great tit, song sparrow, and blue tit assemblies).

We used two estimates of the per-site mutation rate *μ*. The first, *μ*_direct_, is a pedigree or parent–offspring trio estimate for the focal species or its closest available relative, as compiled by Lewin and Eyre-Walker [11]. Species-specific direct estimates were available for the blue tit, collared flycatcher, and rhesus macaque [39, 56–58]; the remaining populations were assigned the direct estimate of their closest relative (Table S3, including feliform carnivore estimates from Armstrong et al. [59]). The second estimate, *μ*_phylo_, is the phylogenetic mixed-model prediction of Wang and Obbard [60], matched to the same species as *μ*_direct_ (Table S3) except for the meerkat, for which we used the domestic cat (*Felis catus*) prediction. Because the phylogenetic model is trained on the direct estimates, the two rates are not independent, so we did not combine them; we report results under *μ*_direct_ and repeated every analysis under *μ*_phylo_, which is 7% to 60% higher.

The primary diploid autosomal deleterious mutation rate was *U*_*d*_ = 4*μL*_coding_, with *L*_coding_ = *Σ* _*i*_ *L*_coding,*I*_ (subsection 2.2), which ranges from 0.34 to 1.26 under *μ*_direct_. The factor of 4 counts 2*μ*mutations per base pair of the diploid genome and a regulatory target equal in length to the exome. Treating every mutation in this target as deleterious, synonymous sites included, biases the maintenance test toward passing (subsection 2.7). Each chromosome’s share of the target in the linked model (subsubsection 2.6.2) is *f*_*i*_ = *L*_coding,*i*_*/L*_coding_. For a larger target, we took for each autosome the union of coding sequences, untranslated regions (UTRs) of protein-coding transcripts, and exons of non-coding RNA genes (*L*_annot,*i*_), plus a fraction *ζ* of the remaining autosomal sequence (*L*_other,*i*_ = *L*_*i*_ *− L*_annot,*i*_), and evaluated *U*_*d*_(*ζ*) = 2*μΣ* _*i*_(*L*_annot,*i*_ + *ζL*_other,*i*_) for *ζ* from 0% to 15%, updating *f*_*i*_ accordingly. Because the B10K hihi annotation includes only CDS features, we imputed its UTR and non-coding RNA fractions from the mean of the four chromosome-level passerine assemblies.

### 3.4 Assigning a distribution of fitness effects

We assigned each population an empirical distribution of fitness effects (DFE) from the gamma DFEs of new nonsynonymous mutations that Lin et al. [40] estimated across species with a uniform pipeline. Lin et al. converted population-scaled selection coefficients to *s* with each species’ ancestral effective size and assumed additive effects, so 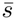 is the mean heterozygous selection coefficient. Because they detected strong phylogenetic signal in the mean deleterious effect across vertebrates (Pagel’s *λ* = 0.84 for 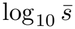) and greater DFE similarity within than between vertebrate classes, we assigned DFEs by phylogenetic proximity (Table S3). Where a species-specific estimate was available (collared flycatcher), we assigned its own DFE; where two reference species were equally related to the focal taxon, we used an equal mixture of their DFEs. The remaining passerines received an equal mixture of the pied and collared flycatcher (*Ficedula* genus) DFEs; the ruminants (bighorn sheep, Soay sheep, and red deer) received an equal mixture of the vaquita and fin whale DFEs (the closest available relatives within Cetartiodactyla); the primates (rhesus macaques and yellow baboons) received the human DFE; and the rodents received the house mouse DFE. Because the carnivore (wolf) DFE fits in Lin et al. [40] had unconstrained heavy tails, we assigned the spotted hyena and meerkat the cetacean mixture as the closest well-constrained reference within Laurasiatheria. The resulting mean heterozygous effects range from 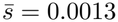 to 0.0195. Because the mean of a gamma DFE fitted to site-frequency spectra depends on its poorly constrained tail of large effects, and the assigned means came from species as distant as the flycatchers for every passerine and the cetaceans for the ruminants and carnivores, we also varied the mean together with the target and the estimate in our joint analysis (subsection 3.6).

Phylogenetic signal within mammals alone was weaker (Pagel’s *λ* = 0.45, non-significant), and the human DFE of Lin et al. [40] has a higher mean than earlier human estimates [61]. However, a smaller 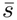 makes the maintenance test harder to pass, by lowering 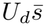, and the diversity test easier, by slowing purging (*Z*_*k*_ = 1 *− s*_*k*_) and strengthening linked selection, so no choice of DFE favors both tests. We applied the same assigned DFE to the maintenance test, the linked reduction, and the threshold analysis. For a gamma DFE, the variance weights 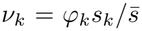 (subsubsection 2.6.2) follow a gamma density with the same scale and a shape larger by one, which we discretized into 240 logarithmically spaced classes over *s ∈* [10^*−*8^, 1] (relative error in 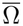 below 2 *×* 10^*−*3^ against 960 classes).

### 3.5 Maintenance test

For every posterior draw of *V*_*A*_, we computed *s*^*\**^ = *V*_*A*_*/U*_*d*_ (Equation (6)), pairing each draw with a draw of *U*_*d*_ from the log-normal distribution implied by the 95% confidence interval of the mutation rate (direct or phylogenetic), and tested each draw of *s*^*\**^ against the mean 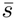 of the assigned DFE (subsection 3.4). We report the posterior mode and 95% HPD interval of *s*^*\**^ and the posterior probabilities 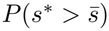 and *P* (*s*^*\**^ *>* 1), the second of which is the probability that the variance exceeds the deleterious mutation rate itself. To check the strong-selection assumption of MSDB (*N*_*e*_*s ≫* 1), we evaluated 2*N*_*e*_*s*^*\**^ in populations with diversity data, at *N*_*e*_ = *π/*(4*μ*) and *s*^*\**^ at its posterior mode, and checked whether *N*_*e*_ exceeds the effective size *U*_*d*_*/*(2*V*_*A*_) below which selection is weak relative to drift (Appendix A.2).

### 3.6 Joint sensitivity of the maintenance test

We varied the deleterious target, the mean deleterious effect, and the pedigree estimate together for the 18 populations with an annotated assembly. The target was either the primary target or the annotated functional sequence plus a fraction *ζ* = 0, 3, 8, or 15% of the unannotated sequence. To represent large-effect mutations that site-frequency spectra do not detect, we gave 0, 0.5, 1, 2, or 5% of new mutations a heterozygous effect of 0.5 and the remainder the assigned DFE, which raised the mean effect by 0.0025 to 0.025; applying this tail to the whole target, coding or not, biases the test toward passing. To represent inflation of the pedigree estimate, for instance by shared environments, we divided it by 1, 2, 3, 5, or 10. In each of the 125 combinations, we counted the populations in which the estimate, at its posterior mode and the point value of the direct mutation rate, still exceeded the variance that mutation maintains, and computed the posterior probability of that excess as in the maintenance test.

### 3.7 Linkage maps and effective size at linked sites

Genetic map lengths were compiled from published pedigree linkage maps (Table S3). For yellow baboons, we used the map of a captive *Papio hamadryas* colony [62], substituting chromosomes 5, 12, and 18 from its second-generation revision [63]. The rhesus macaque map [64] is male-only and from a captive colony of Indian origin rather than the Cayo Santiago population; because female recombination rates exceed male rates in most mammals, it overstates the linked reduction, which biases the diversity test toward passing. All other maps are sex-averaged (unconfirmed for the baboon map). We matched linkage groups to assembly chromosomes by name, and microchromosomes and other ambiguous groups by concordance between physical and genetic map length. For the song sparrow, whose assembly numbers chromosomes by physical size, each chromosome was assigned to the great tit chromosome [65] sharing the plurality of its annotated protein-coding genes (shared fraction 0.59 to 1.0), and through it to the corresponding song sparrow linkage group. Map lengths *M*_*i*_ are in Morgans. Because the published blue tit map [66] is fragmentary, the hihi assembly [67] is not chromosome-level, and the superb fairy-wren assembly [55] is unannotated, the mapped set comprises 9 populations, and the remaining 10 were analyzed only at unlinked sites. Every autosome with a map length of at least 1 cM, including microchromosomes, contributed to the linked term.

We partitioned 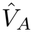 among autosomes in proportion to coding length, 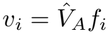, and report the partition by annotated gene counts as a sensitivity analysis. We computed 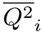 from its closed form under Haldane’s map function (Equation (D7)), which agreed with numerical integration of Equation (D4) to within one part in 10^11^ for every map length and selection coefficient examined. The association with each effect class was retained at *Z*_*k*_ = 1 *− s*_*k*_ (Equation (10)), and the linked and unlinked terms were integrated over the classes of the assigned DFE with weights *ν*_*k*_ (subsection 3.4), using the same retention factor in both terms of Equation (19). To separate the effect of the linkage maps from that of the DFE, we also evaluated a uniform *s* = 0.01 and the population-specific *s* = *s*^*\**^ at which MSDB maintains 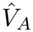. When testing against an observed deficit, we evaluated the unlinked factor at *Q* = 2 under the multiplicative bound, a combination biased toward passing, and also under the small-variance approximation.

### 3.8 Nucleotide diversity, census size, and the diversity test

For 7 populations, we used the species-mean nucleotide diversity *π* compiled by Lewin and Eyre-Walker [11], which averages published estimates across fourfold-degenerate, synonymous, or genome-wide sites [13, 39, 68–71]. None of these estimates comes from a focal study population; 5 are for the study species itself, whereas the estimates for red deer and yellow baboons are based on *Cervus elaphus yarkandensis* and *Papio anubis*, respectively. For 4 additional populations, we used published whole-genome estimates from other populations of the same species: for the two great tit populations, diversity across all sites in 10 European males (potentially including the Z chromosome) [72]; for bighorn sheep, 6 Rocky Mountain genomes mapped to the domestic sheep reference [73]; and for song sparrows, the coastal subspecies *M. m. rufina* [74]. The bighorn and song sparrow values come from SNP panels filtered on minor allele frequency and are therefore probably biased downward. Because none of these values is restricted to neutral autosomal sites, and fourfold and synonymous sites are themselves affected by linked selection, *π* only approximates neutral diversity. We excluded estimates of heterozygosity from a single genome. We treated *π/*(4*μ*) as the coalescent effective size [11], which estimates the harmonic mean of *N*_*e*_ over the genealogy that the diversity test concerns (Equation (11)). Every genomic estimate of *N*_*e*_, including trajectories inferred by sequentially Markovian coalescent and site-frequency-spectrum methods, already includes the drift caused by heritable variance in fitness, so none supplies the long-term number of breeding adults that the test requires.

Total census sizes *N*_*c*_ came from Buffalo [10] for 10 of the 19 populations (9 after excluding the domestic sheep value for the St. Kilda Soay sheep). These are density *×* geographic range estimates, a contemporary quantity that counts all adults and, in some density surveys, juveniles [11]. Because Buffalo calculated bird densities from body masses predicted from body length, the resulting bird census sizes are probably too low; recomputing densities from measured body masses [75] with Damuth’s avian allometry [76] raises blue tit and collared flycatcher densities by about 1.2 orders of magnitude. Because no population has a count of breeding adults, 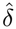 (subsection 2.5) uses *N*_*c*_ in place of *N*, which lowers it below *δ* by the breeder fraction (a factor of 2 to 10) and widens the apparent gap between predicted and observed diversity. The species-level values describe the two Corsican blue tit populations (Pirio and Muro) less closely than the mainland population at Rouvière. Corsican blue tits have been isolated from the mainland for about 4,500 generations, with a long-term effective size about 7.5 times smaller and 13% lower heterozygosity at polymorphic sites [77], but because two Corsican lineages have only about a 4% chance of coalescing on the island within that time, most of their genealogy predates the isolation and is described by the species-level values. Table S3 records the source of each parameter, and Table 2 summarizes the direction in which each source of error shifts the apparent diversity gap.

**Table 2:**
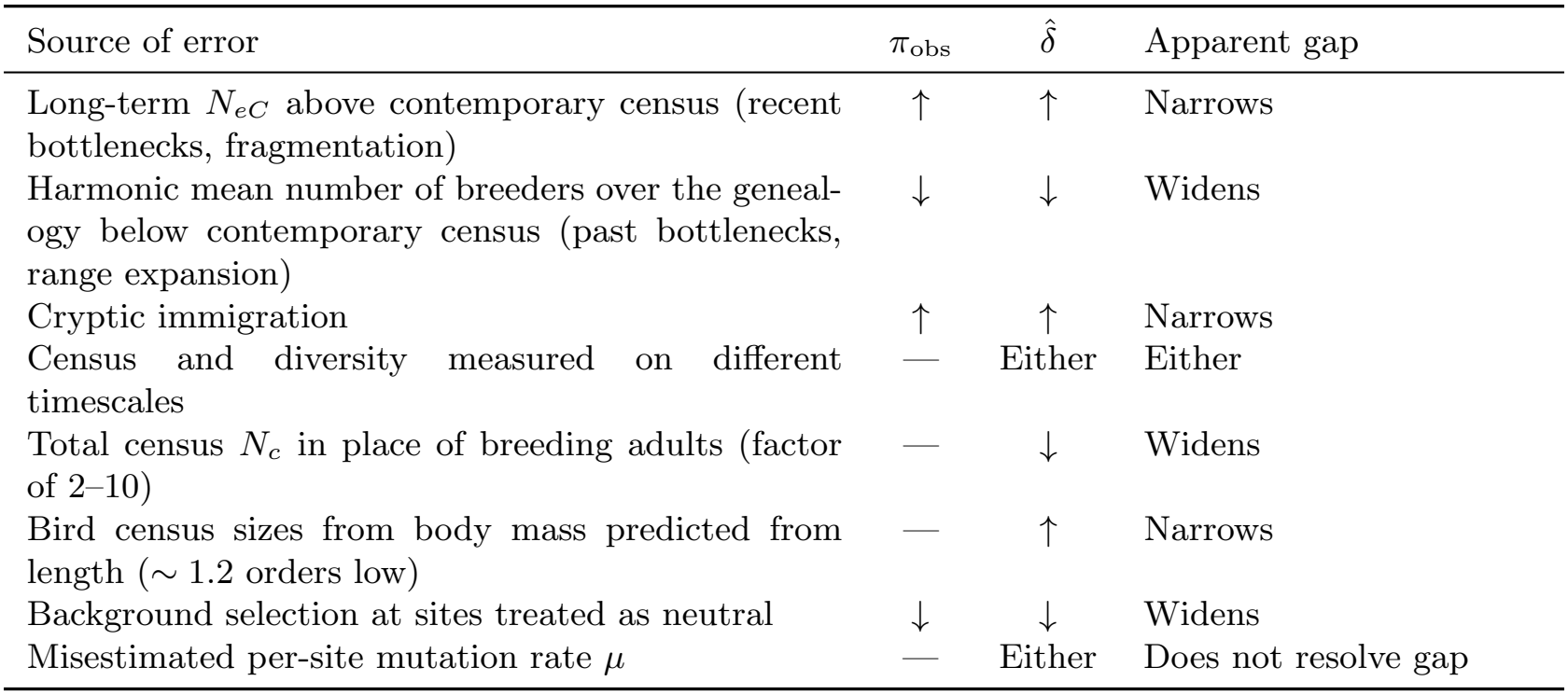
Sources of error in the diversity ratio alter the apparent diversity gap in both directions. Directions indicate how each source of error shifts observed diversity *π*_obs_, the estimated diversity ratio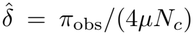, and the apparent gap between predicted and observed diversity. Dashes indicate no directional effect.

| Source of error | π<sub>obs</sub> | δ̂ | Apparent gap |
|---|---|---|---|
| Long-term *N<sub>eC</sub>* above contemporary census (recent bottlenecks, fragmentation) | ↑ | ↑ | Narrows |
| Harmonic mean number of breeders over the geneal-ogy below contemporary census (past bottlenecks, range expansion) | ↓ | ↓ | Widens |
| Cryptic immigration | ↑ | ↑ | Narrows |
| Census and diversity measured on different timescales | — | Either | Either |
| Total census *N<sub>c</sub>* in place of breeding adults (factor of 2–10) | — | ↓ | Widens |
| Bird census sizes from body mass predicted from length (∼ 1.2 orders low) | — | ↑ | Narrows |
| Background selection at sites treated as neutral | ↓ | ↓ | Widens |
| Misestimated per-site mutation rate *μ* | — | Either | Does not resolve gap |

Nine populations from 6 species have estimates of both *π* and *N*_*c*_. We call them the *deficit set* ; 6 of them also belong to the mapped set of subsection 3.7. For these 9 populations, we tested *π*_pred_ = 4*μN*_*c*_ (*N*_*e*_*/N* )_pred_ against *π*_obs_, with (*N*_*e*_*/N* )_pred_ under both the small-variance approximation and the multiplicative bound. A population with 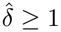 would be a failed model check; none reached it. We evaluated the minimum long-term variance required to account for each observed deficit, *V*_*A,δ*_ (Equation (14)), and its linked counterpart as the root of Equation (21) set equal to 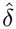, found with uniroot in R on log_10_ *V*_*A*_.

### 3.9 Threshold analysis and compatibility test

We expressed the long-term variance as 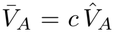 (subsection 2.1) and followed two ratios as functions of *c* in each of the 9 populations in the deficit set: the maintenance ratio 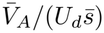 at the assigned DFE, and the ratio of predicted to observed diversity, 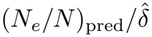. Each test passes where its ratio is *≤* 1. The ratio of predicted to observed diversity was evaluated under the multiplicative bound, which is biased toward passing at every 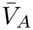, with the small-variance approximation shown alongside it for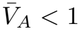. The values of *c* at which the two ratios reach 1 are the thresholds *c*_MSD_ and *c*_div_ (subsection 2.5), and a population passes the compatibility test where *c*_div_ *≤ c*_MSD_. Because some species contribute several populations that share *π* and *N*_*c*_, we averaged the log ratios within each species and reported the cross-species mean with a *t*-based 95% confidence interval. With linkage, the predicted *N*_*e*_*/N* was multiplied by 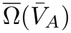 at the assigned DFE, and we report the linked prediction at 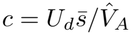 as the prediction of MSDB with background selection.

### 3.10 Uncertainty carried by each test

Posterior uncertainty in *V*_*A*_ was propagated through every derived quantity (*N*_*e*_*/N, s*^*\**^, and all linked factors), and uncertainty in the mutation rate through *U*_*d*_ in the maintenance test (subsection 3.5) and through its 95% confidence interval in the diversity ratio. All other inputs were fixed at point values: total census sizes *N*_*c*_, nucleotide diversity *π* (including the 4 proxy values from other populations), DFE parameters (beyond the spread of the mixture between reference species), extra-pair paternity rates *α*, and linkage maps *M*_*i*_. A quantity that lies above or below a reference value across its HPD interval therefore does so given the posterior uncertainty of *V*_*A*_ alone, and we evaluated census-size errors separately where they could change a verdict.

### 3.11 Independence of the inputs from diversity data

None of the parameters of either test was estimated from the diversity *π*_obs_ against which the diversity predictions are evaluated. Diversity is otherwise used only to check the strong-selection assumption through 2*N*_*e*_*s*^*\**^, so the diversity test is a prediction from independently measured inputs, not a fit. The one source shared across parameters, Lewin and Eyre-Walker [11], compiles both *μ*_direct_ and *π*, which are measured by independent methods, de novo mutations in parent–offspring trios and pairwise differences among unrelated individuals.

## 4 Results

### 4.1 Heritable variance in fitness implies a modest reduction in effective size at unlinked sites

Table 3 and Figure 1 give the reduction in effective size at unlinked sites implied by the pedigree estimates of the additive genetic variance in relative fitness, *V*_*A*_. Under the multiplicative model, *N*_*e*_*/N* ranges from 0.021 to 0.98, with a median of 0.53 (Figure 1a, gray circles). The median reduction of 47% lies within the 16 to 55% that Charlesworth and Jensen [35] obtained from the mean of the same estimates with the small-variance approximation. The predicted reductions are of the same order as those commonly attributed to variance in family size and unequal sex ratios [49]. The largest predicted reductions occur where 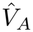 is largest, in the blue tits at Pirio and Muro and in the spotted hyenas; at Muro, *N*_*e*_*/N* = 0.021 is close to the 0.019 that the multiplicative bound gives. For the same *V*_*A*_, socially monogamous populations have a lower *N*_*e*_*/N* than randomly mating populations because social partners share each other’s advantage (subsubsection 2.6.1; Figure 1b).

**Table 3:**
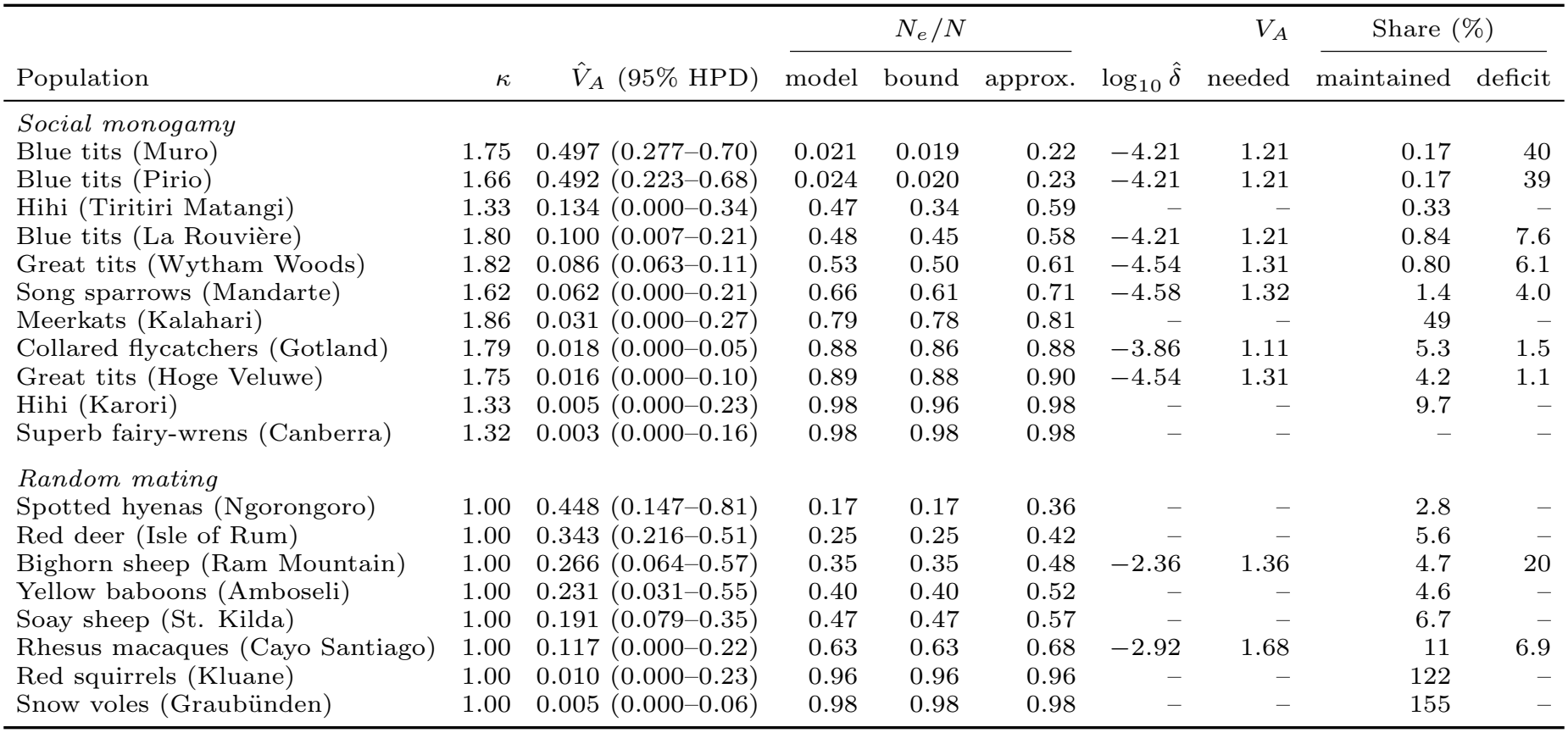
Under multiplicative fitness, the measured *V*_*A*_ accounts for a small share of each observed deficit, and in most populations recurrent mutation maintains a small share of the measured *V*_*A*_.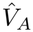: posterior mode and 95% HPD interval [9]. Populations are grouped by mating system and listed by species and study site. *N*_*e*_*/N* at the posterior mode: the multiplicative model; the multiplicative bound; and their limit for small *V*_*A*_, the small-variance approximation.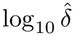: observed diversity relative to its neutral expectation at the census size, 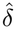, on a log scale, where a diversity estimate and a census size are available (Methods); its reciprocal, 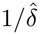, is the deficit. *V*_*A*_ needed: the minimum variance that could account for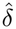. Shares, in percent, at the posterior mode and the point value of the direct mutation rate. Maintained: the share of 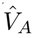that recurrent mutation maintains at the assigned DFE,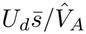; values above 100 mean that mutation maintains all of it. Deficit: the share of the observed deficit that 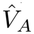 accounts for under the multiplicative model, on a logarithmic scale, ln[1*/*(*N*_*e*_*/N* )]*/* ln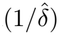.

|  |  |  | N<sub>e</sub>/N |  |  |  | V<sub>A</sub> | Share (%) |  |
|---|---|---|---|---|---|---|---|---|---|
| Population | κ | V̂<sub>A</sub> (95% HPD) | model | bound | approx. | log<sub>10</sub> δ̂ | needed | maintained | deficit |
| *Social monogamy* |  |  |  |  |  |  |  |  |  |
| Blue tits (Muro) | 1.75 | 0.497 (0.277–0.70) | 0.021 | 0.019 | 0.22 | −4.21 | 1.21 | 0.17 | 40 |
| Blue tits (Pirio) | 1.66 | 0.492 (0.223–0.68) | 0.024 | 0.020 | 0.23 | −4.21 | 1.21 | 0.17 | 39 |
| Hihi (Tiritiri Matangi) | 1.33 | 0.134 (0.000–0.34) | 0.47 | 0.34 | 0.59 | – | – | 0.33 | – |
| Blue tits (La Rouvière) | 1.80 | 0.100 (0.007–0.21) | 0.48 | 0.45 | 0.58 | −4.21 | 1.21 | 0.84 | 7.6 |
| Great tits (Wytham Woods) | 1.82 | 0.086 (0.063–0.11) | 0.53 | 0.50 | 0.61 | −4.54 | 1.31 | 0.80 | 6.1 |
| Song sparrows (Mandarte) | 1.62 | 0.062 (0.000–0.21) | 0.66 | 0.61 | 0.71 | −4.58 | 1.32 | 1.4 | 4.0 |
| Meerkats (Kalahari) | 1.86 | 0.031 (0.000–0.27) | 0.79 | 0.78 | 0.81 | – | – | 49 | – |
| Collared flycatchers (Gotland) | 1.79 | 0.018 (0.000–0.05) | 0.88 | 0.86 | 0.88 | −3.86 | 1.11 | 5.3 | 1.5 |
| Great tits (Hoge Veluwe) | 1.75 | 0.016 (0.000–0.10) | 0.89 | 0.88 | 0.90 | −4.54 | 1.31 | 4.2 | 1.1 |
| Hihi (Karori) | 1.33 | 0.005 (0.000–0.23) | 0.98 | 0.96 | 0.98 | – | – | 9.7 | – |
| Superb fairy-wrens (Canberra) | 1.32 | 0.003 (0.000–0.16) | 0.98 | 0.98 | 0.98 | – | – | – | – |
| *Random mating* |  |  |  |  |  |  |  |  |  |
| Spotted hyenas (Ngorongoro) | 1.00 | 0.448 (0.147–0.81) | 0.17 | 0.17 | 0.36 | – | – | 2.8 | – |
| Red deer (Isle of Rum) | 1.00 | 0.343 (0.216–0.51) | 0.25 | 0.25 | 0.42 | – | – | 5.6 | – |
| Bighorn sheep (Ram Mountain) | 1.00 | 0.266 (0.064–0.57) | 0.35 | 0.35 | 0.48 | −2.36 | 1.36 | 4.7 | 20 |
| Yellow baboons (Amboseli) | 1.00 | 0.231 (0.031–0.55) | 0.40 | 0.40 | 0.52 | – | – | 4.6 | – |
| Soay sheep (St. Kilda) | 1.00 | 0.191 (0.079–0.35) | 0.47 | 0.47 | 0.57 | – | – | 6.7 | – |
| Rhesus macaques (Cayo Santiago) | 1.00 | 0.117 (0.000–0.22) | 0.63 | 0.63 | 0.68 | −2.92 | 1.68 | 11 | 6.9 |
| Red squirrels (Kluane) | 1.00 | 0.010 (0.000–0.23) | 0.96 | 0.96 | 0.96 | – | – | 122 | – |
| Snow voles (Graubünden) | 1.00 | 0.005 (0.000–0.06) | 0.98 | 0.98 | 0.98 | – | – | 155 | – |

**Figure 1:**
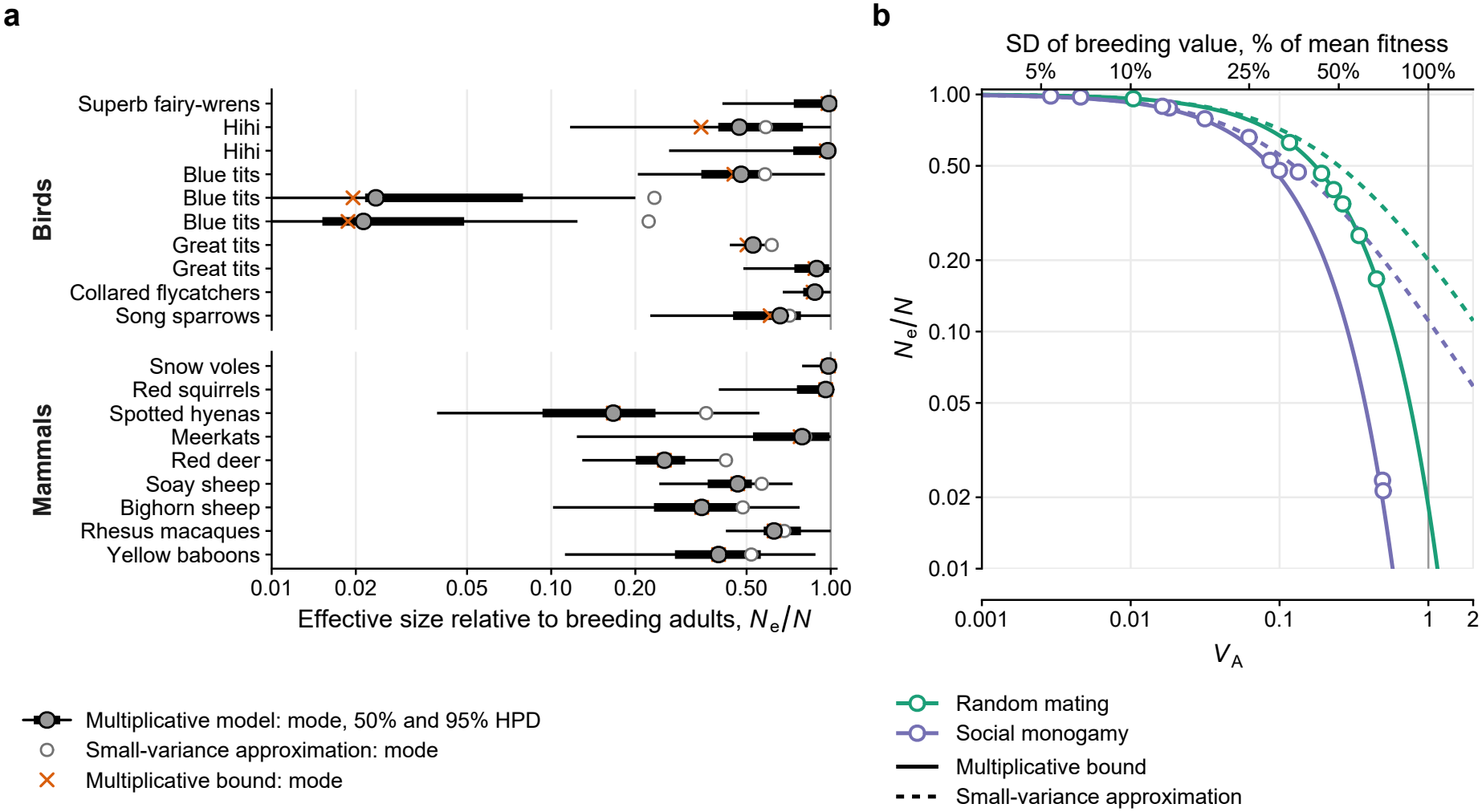
The measured additive variance in relative fitness predicts a modest reduction in effective size at unlinked sites. (a) *N*_*e*_*/N*, effective size relative to the number of breeding adults, for the 19 populations of Bonnet et al. [9]; populations of the same species appear in the order of Table S3, which gives their locations. Gray circles and bars: the multiplicative model at the posterior mode of 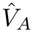, with 50% (thick) and 95% (thin) HPD intervals. Open circles: the small-variance approximation, the limit of the multiplicative model for small *V*_*A*_. Crosses: the multiplicative bound, which under random mating coincides with the multiplicative model. (b) The same predictions against *V*_*A*_ at *κ* = 1 (random mating, green) and *κ* = 2 (strict monogamy, purple), the multiplicative bound (solid) and the small-variance approximation (dashed). Points: each population at its posterior mode under the multiplicative model, colored by mating system; socially monogamous populations lie between the curves because extra-pair paternity lowers *κ* below 2. Top axis: the standard deviation of breeding value as a percentage of mean fitness,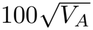; the vertical line marks *V*_*A*_ = 1, where it equals mean fitness.

The small-variance approximation understates the largest predicted reductions and gives *N*_*e*_*/N* from 0.22 to 0.98, with a higher median of 0.61 (Figure 1a, open circles). The multiplicative model predicts a larger reduction because, with expected fitness exponential in the breeding value, the variance of long-term contributions grows exponentially, as 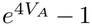, rather than linearly, as 4*V*_*A*_ (subsection 2.4). Lineages in the upper tail of breeding values therefore leave more descendants than the small-variance approximation allows.

In the 14 populations with 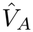 below 0.25, the multiplicative model predicts an *N*_*e*_*/N* within a quarter of the small-variance approximation, so the prediction depends little on how loci combine to determine fitness. The difference exceeds a quarter only in the five populations with larger estimates, the blue tits at Pirio and Muro, spotted hyenas, red deer, and bighorn sheep, and is about tenfold in those two blue tit populations.

### 47.2 Heritable variance in fitness accounts for a small share of each observed deficit

Because neutral diversity in the deficit set lies orders of magnitude below what census size predicts, reductions this modest are expected to account for a small part of each deficit. The diversity ratio, 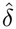, ranges from 10^*−*2.36^ in bighorn sheep to 10^*−*4.58^ in song sparrows, deficits of 230- to 38,000-fold (Table 3). On the logarithmic scale on which successive reductions combine (subsection 2.5), 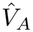 accounts under the multiplicative model for a median of 6.9% of the deficit (quartiles 4.0–19.5%), from 1.1% in the great tits at Hoge Veluwe to 40% in the blue tits at Muro (Table 3; Figure 2). The share is 1.2 to 41% (median 6.9%) under the multiplicative bound and 1.0 to 15.5% (median 5.6%) under the small-variance approximation. At the posterior modes of 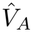, predicted diversity exceeds observed diversity even under the multiplicative bound, by a factor of 80 (bighorn sheep) to 31,000 (great tits at Hoge Veluwe). These factors are also the smallest by which *N*_*c*_ would have to overstate the long-term number of breeding adults for heritable variance in fitness to account for each deficit (subsection 2.5). The minimum variance that could account for each deficit, *V*_*A,δ*_ (Equation (14); Figure 2, orange diamonds), is 1.1 to 1.7, above the upper limit of the 95% HPD interval of all nine estimates (at most 0.70). *V*_*A,δ*_ stays above that limit across the 95% CI of the mutation rate (smallest ratio 1.66, Muro blue tits) and when only breeding adults are counted, which would raise 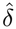 up to tenfold (Table 2; smallest ratio 1.17, bighorn sheep at the lower end of the mutation-rate CI). Even combined with the lower end of the demographic range, *N*_*eV*_ */N* = 0.1 [49], the predicted reduction accounts for a deficit of about 20-fold at most in the median population.

**Figure 2:**
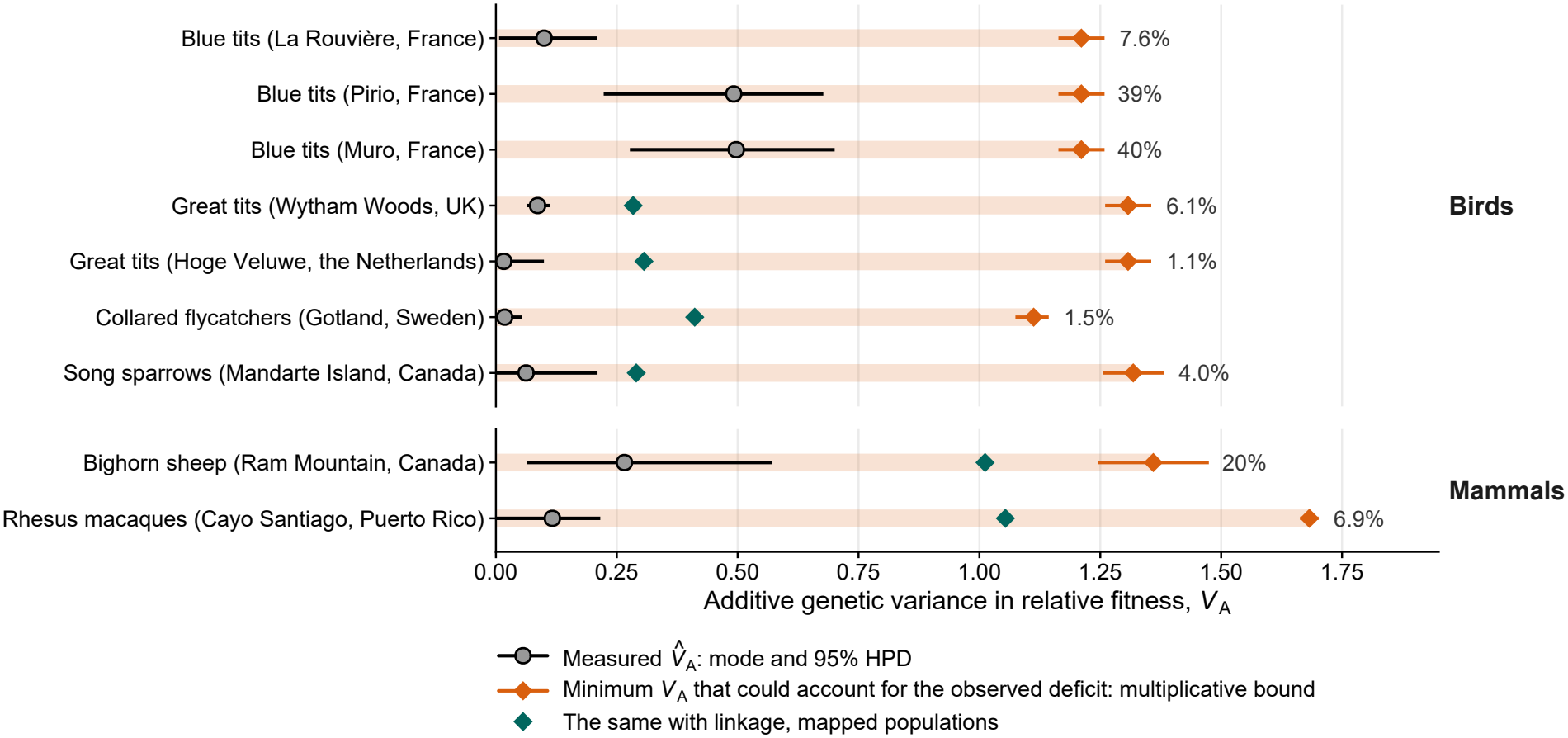
Each pedigree estimate is a small share of the additive variance in relative fitness that its population’s observed deficit would require. One row for each of the nine populations with *π* and a census size. Gray circles and bars: the posterior mode of 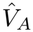 and its 95% HPD interval. Orange diamonds: the minimum *V*_*A*_ that could account for the observed deficit, *V*_*A,δ*_, under the multiplicative bound at the population’s *κ*_max_, with bars over the 95% CI of the mutation rate; the shaded track extends from zero to this value. Teal diamonds: the same with linkage, at the assigned empirical DFE, for the six mapped populations. Percentages: the share of each deficit that 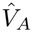 accounts for under the multiplicative model, on a logarithmic scale (Table 3); the position of the estimate along its track is the same share under the multiplicative bound. A population would pass the diversity test where its estimate reaches its diamond.

A long-term 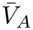 of 1.1 to 1.7 is large. Under the multiplicative model, an individual whose breeding value lies one standard deviation above the mean would expect three to four times as many offspring as an individual at the mean, a large but not inconceivable difference in a single generation, and a much harder difference to sustain over the roughly 2*N*_*e*_ generations back to the common ancestor of two gene copies. Under the multiplicative bound, each required value is 70 to 95% (median 88%) of the variance at which the upper-tail ancestors that produce the reduction become rarer than one per generation among *N*_*c*_ breeding adults, and 48 to 83% if one adult in ten breeds (Appendix B.3). Because diversity reflects the harmonic mean of *N*_*e*_ over the genealogy (Equation (11)), the variance measured over a pedigree counts toward each required value only in proportion to the share of the genealogy that the pedigree spans. That share is very small, because a pedigree covers a few decades, whereas the diversity of the deficit set reflects *π/μ≈* 4*N*_*e*_, or about 90,000 to 1,000,000, generations.

### 4.3 Linkage amplifies the reduction only where purging is slow

For the mapped set (subsection 3.7), Figure 3 combines the reduction at unlinked sites under the multi-plicative bound with the reduction at linked sites along each chromosome, at 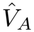 and the assigned empirical DFE. Linkage lowers *N*_*e*_*/N* from 0.25–0.63 to 0.15–0.44 in mammals. In birds, *N*_*e*_*/N* with linkage is 0.66 in collared flycatchers and 0.43 in the great tits at Hoge Veluwe but 0.066 in song sparrows and 0.016 in the great tits at Wytham Woods, differences that follow mainly from 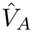. With linkage, 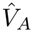 accounts on the logarithmic scale for 4.7 to 40% of each observed deficit (median 19%) in the six mapped populations of the deficit set, against 1.2 to 20% under the multiplicative bound without linkage. The shortfall at the posterior mode, which is also the factor by which *N*_*c*_ would have to overstate the long-term number of breeding adults, ranges from 53-fold (bighorn sheep) to 15,000-fold (great tits, Hoge Veluwe). With the small-variance approximation as the unlinked factor it ranges from 75- to 15,000-fold, so the shortfall depends little on the fitness mapping assumed. The variance each deficit requires with linkage also exceeds the upper limit of each population’s 95% HPD interval at any mutation rate within its 95% CI, although linkage lowers it from 1.1–1.7 to 0.28–0.41 in the four bird populations of both sets and to about 1.0 in bighorn sheep and rhesus macaques (Figure 2, teal diamonds). The margin is narrowest for bighorn sheep and song sparrows. The upper HPD limit of bighorn sheep gives *N*_*e*_*/N* = 0.045 against an observed diversity ratio of 0.0043. If only one adult in ten breeds, the variance that the deficits of these two populations require is within a few percent of their upper HPD limits at the point value of the mutation rate and below those limits at the lower end of its CI. The song sparrows would also reach their limit if the bird census sizes were too high and not too low.

**Figure 3:**
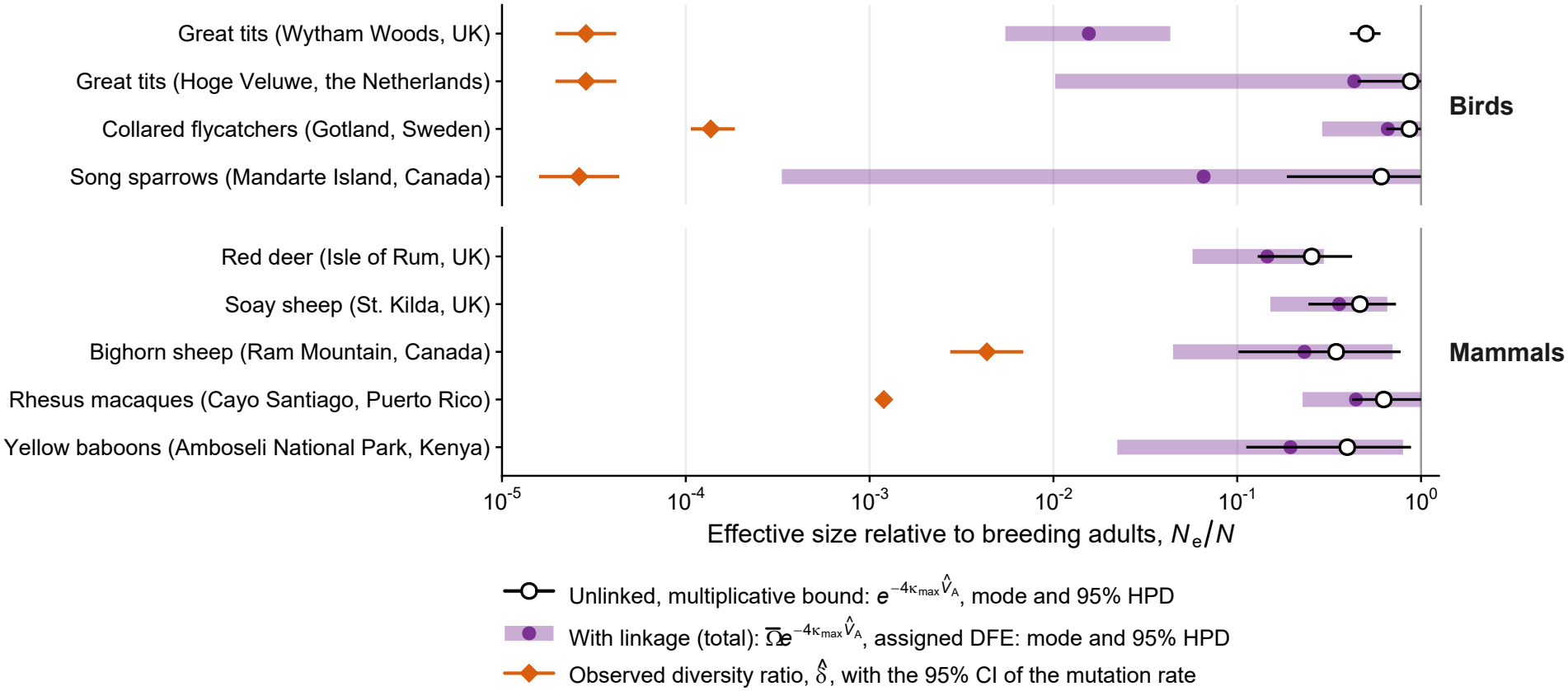
Linkage amplifies the reduction in effective size, yet the measured variance still accounts for a small part of each observed deficit. Genome-wide *N*_*e*_*/N* for the nine populations with linkage maps. Open circles: unlinked, multiplicative bound, mode and 95% HPD. Purple: with linkage, at the empirical DFE assigned to each population (Methods); dot at the mode, bar over the 95% HPD of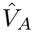.Orange diamonds: the observed diversity ratio, 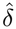, with bars over the 95% CI of the mutation rate. Red deer and yellow baboons lack an own-species census size, and Soay sheep a diversity estimate.

The linked term is an upper limit on what linkage adds, because it attributes the measured variance to alleles purged as slowly as the assigned DFE implies, which stay associated with linked neutral sites longest but could not maintain that variance (subsubsection 2.6.2). When we instead give every allele the effect *s*^*\**^ at which MSDB maintains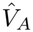, the linked factor, which is the genome-wide *N*_*e*_*/N* with linkage relative to its value without, is 0.97–0.99 in every population, because alleles of that effect are purged within a few generations, before recombination would separate them from most linked sites. Chromosomes that carry more selected target per unit of map length are predicted to lose more of their neutral diversity, and the chromosomes of both taxa, both macrochromosomes and microchromosomes, lie on the same curve (Figure S4c). The contrast between birds and mammals arises from the assigned DFEs and not from the maps. Partitioning the variance by gene count or counting part of the unannotated sequence as target does not change these results (Supplementary Results). The measured variance is small relative to the deficits. As we show next, it is nonetheless larger in most populations than recurrent mutation maintains at the mean deleterious effects assigned from related species.

### 4.4 Maintaining the measured variance would require unrealistic mean deleterious effects

For the 18 populations with an annotated assembly, the mean heterozygous effect that MSDB would require to maintain 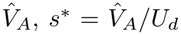 (Equation (6)), has posterior modes from 0.0064 in snow voles to 0.64 in the blue tits at Muro and the spotted hyenas, with a median of about 0.16 (Figure 4). The assigned empirical DFEs have means of 0.0013–0.0021 in birds and 0.010–0.020 in mammals, and the posterior probability that *s*^*\**^ exceeds the assigned mean is at least 0.89 in 17 of the 18 populations (0.72 in snow voles, whose estimate is close to zero). At the posterior modes of 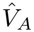 and the point value of the direct mutation rate, *s*^*\**^ is 21 times the assigned mean in the median population, 10 to 590 times in birds and up to 35 times in mammals. Equivalently, 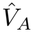 exceeds the variance that recurrent mutation maintains at the assigned DFE, 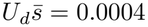 to 0.02, in 16 of 18 populations, all but the snow voles and red squirrels, and by less than threefold only in the meerkats (Figure S8). The share of 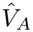 that recurrent mutation maintains, 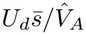 (subsection 2.5), has a median of 4.7% (quartiles 1.0–9.0%), from 0.17% in the blue tits at Muro to 155% in the snow voles; it is 49% in the meerkats and 0.17 to 11% (median 1.4%) in the deficit set (Table 3). In four populations, the spotted hyenas, the blue tits at Muro and Pirio, and the hihi on Tiritiri Matangi, more than 5% of the posterior lies where 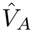 exceeds *U*_*d*_ itself, a variance that no selection coefficient can maintain, because a mutation cannot remove more than all of its carrier’s fitness.

**Figure 4:**
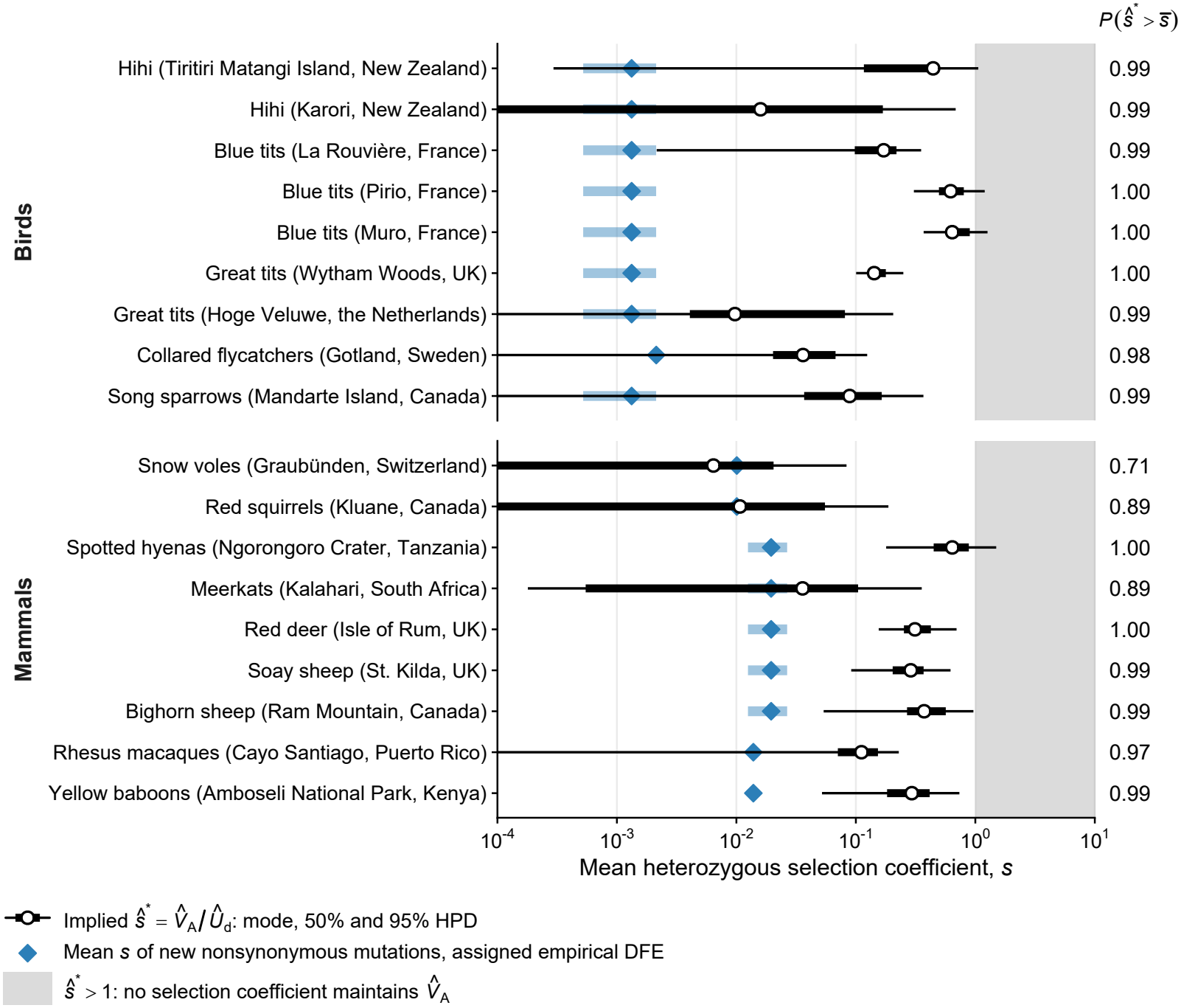
Maintaining each measured variance at MSDB would require mean selection coefficients above the mean effects assigned from related species. The mean heterozygous selection coefficient *s*^*\**^ that MSDB would require, for the 18 populations with an annotated assembly. Circles and bars: posterior mode, 50% (thick) and 95% (thin) HPD over the 4,000 posterior draws of *V*_*A*_, with *U*_*d*_ drawn from the 95% CI of the direct mutation rate. Blue diamonds: mean *s* of the empirical DFE assigned to the population (Methods); where the assignment is an equal mixture of two species, the blue bar spans their means. Gray: *s*^*\**^ *>* 1, where *V*_*A*_ *> U*_*d*_ and no selection coefficient maintains the variance. Right: posterior probability that *s*^*\**^ exceeds the assigned mean.

In the 11 populations for which diversity gives an estimate of *N*_*e*_, 2*N*_*e*_*s*^*\**^ is 2,900 to 320,000, so selection is strong relative to drift and MSDB maintains an expected variance 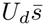. A variance carried by weakly selected mutations would instead require an effective size well below *U*_*d*_*/*(2*V*_*A*_), 3.3 individuals for the median population at the posterior modes of *s*^*\**^ (0.78–78). With the phylogenetic mutation rate, the median *s*^*\**^ is 0.12. The annotated functional target (coding sequence, untranslated regions, and non-coding RNA exons) is 0.8 to 2.0 times the doubled exome, so the primary *U*_*d*_ is close to what the annotation alone gives. In seven of the nine bird populations, the target required to maintain 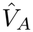 at the assigned DFE would exceed the genome itself. At the effects 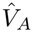 requires, purging would also cut short an ancestor’s advantage at unlinked sites and multiply *Q*^2^ by 1*/*(1 + *s*^*\**^)^2^ (Appendix A.4), 0.37–0.99 (median 0.75), so the *Q* = 2 used above overstates the reduction at unlinked sites.

A tail of large effects hidden from site-frequency spectra raises the maintained variance in proportion to its contribution to the mean effect (subsection 2.2). It would raise the mean to the median *s*^*\**^ only if about 30% of new mutations had a heterozygous effect of 0.5. Counting part of the unannotated sequence as target (*ζ*) lowers 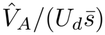 in every population but changes few verdicts. At *ζ* = 15% the median ratio decreases from 21 to 4, and at the 7–9% of non-coding sequence estimated to be deleterious in humans [78] it is 6 or 7. Of the 16 populations that fail at the primary target, 15 still fail at *ζ* = 15%; the exception is the meerkats, which pass once *ζ* exceeds about 3% (Figure S6). Fourteen populations still fail when 5% of new mutations have an effect of 0.5, and fourteen when the estimate is ten times the long-term variance.

Combined, the three inputs bring the median population close to passing, although most populations still fail (subsection 3.6, Figure S9). With 8% of the unannotated sequence, a 1% large-effect tail, and an estimate inflated twofold, 12 populations fail (7 birds and 5 mammals), 7 of them with posterior probability at least 0.95, but the median ratio decreases from 21 to 1.9, so recurrent mutation would maintain about half of the median population’s variance rather than 4.7%. With 15%, a 2% large-effect tail, and a threefold inflation, 6 populations fail, the blue tits at Rouvière, Pirio, and Muro, the hihi on Tiritiri Matangi, the great tits at Wytham Woods, and the spotted hyenas, and the median ratio is 0.63. All 18 pass only when 15%, a 5% tail, and a tenfold inflation are combined, and the two Corsican blue tit populations are the last to pass. Because a large-effect tail adds the same amount to every mean effect, a 1% tail raises the bird means (0.0013 to 0.0021) three- to fivefold but the mammal means, about ten times larger, only by a quarter to a half, so the bird verdict depends mainly on the tail and the mammal verdict on the target and the estimate (Supplementary Results).

### 4.5 No single long-term variance passes both tests

A larger long-term variance brings the diversity test closer to passing and the maintenance test further from it. For the threshold analysis, we rescaled the pedigree estimate, 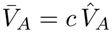, for the deficit set (subsection 3.8) and followed each test as *c* changes (Figure 5). The maintenance test is passed only for *c ≤* 0.024 on average across species (0.0017–0.11 across populations), a long-term variance about forty times smaller than the pedigree estimate, and the diversity test only for *c ≥* 9.4 (2.4–80 across populations, median 14), about ten times larger. Every population in the deficit set fails both tests at the pedigree estimate. The two thresholds are separated by a factor of about 390 on average, and by 108 to 1,890 in individual populations, so every population also fails the compatibility test, the case of Table 1 in which the equilibrium assumed by the two models does not hold. Where the two average curves cross, at *c ≈* 3.9, the long-term variance would be 160 times what MSDB maintains, and predicted diversity would still be 160 times the observed.

**Figure 5:**
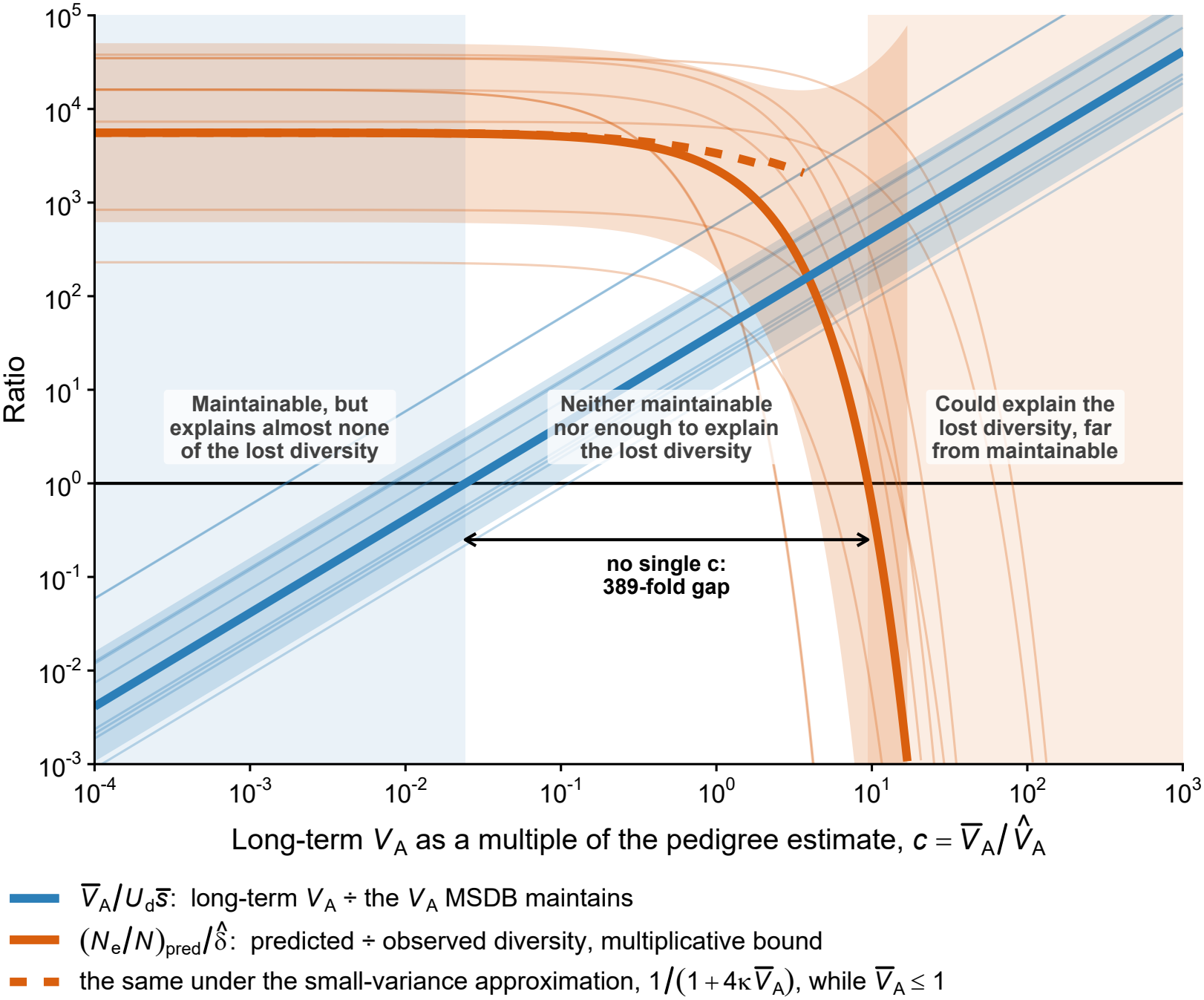
No single rescaling of the pedigree estimates passes both tests. The two tests as functions of the rescaling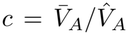, for the nine populations with an observed deficit. Blue: long-term variance relative to the variance MSDB maintains at the assigned empirical DFE,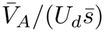. Orange: predicted relative to observed diversity, 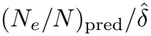, under the multiplicative bound (solid), and under the small-variance approximation while 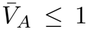 (dashed). Each test is passed where its ratio is at most 1 (black line). Bold lines: mean over species of the log_10_ ratio, populations of a species averaged first, with *t*-based 95% intervals across species (shading). Faint lines: populations. The arrow spans the gap between the two thresholds, the values of *c* at which the two average curves reach 1. Unlinked term only; Figure S10 adds linkage.

The variance that recurrent mutation maintains at each population’s assigned DFE, 0.0007 to 0.013 in the deficit set, would lower neutral diversity at unlinked sites by only 0.6 to 5% even under the multiplicative bound (*N*_*e*_*/N* = 0.95–0.99). The outcome of the compatibility test therefore does not depend on the pedigree estimates, which determine only on which side of each threshold a population lies and by how much (Equation (17)).

Adding linkage narrows the gap but does not resolve it (Figure S10). For the six populations of both sets, the long-term variance that would explain the deficit with linkage is 3.3 to 22.6 times the pedigree estimate in birds and 3.8 to 9.0 times in mammals, still 81 to 442 times what MSDB at the same DFE maintains. At the variance MSDB does maintain, linkage barely changes the prediction, so MSDB with background selection accounts for almost none of the missing diversity. Averaged over species, predicted diversity exceeds observed diversity 4,240-fold with linkage and 4,430-fold without. With 15% of the unannotated sequence counted as target it does so 3,240-fold, and the variance MSDB then maintains removes no more than about half of neutral diversity in any population (rhesus macaques, *N*_*e*_*/N* = 0.51).

## 5 Discussion

We tested pedigree estimates of additive variance in relative fitness from 19 wild vertebrate populations [9] against the two equilibrium predictions that classical population genetics makes for such a variance, its effect on neutral diversity and its maintenance by mutation. Propagated to neutral diversity, the measured variance roughly halves the effective size of the median population if, as the animal model behind the estimates assumes, the effects of loci on fitness multiply and breeding values for log-fitness are normally distributed. A variance this small has that much effect because inheritance adds as much again to an ancestor’s own advantage through its descendants, and partners who share their offspring share each other’s advantage. No account of the missing diversity relies on heritable variance in fitness alone, and as expected this halving is a small part of what each population has lost. It amounts to a median of about a fifteenth of each deficit on a logarithmic scale, even if nothing limits the fitness of the most favored lineages and socially monogamous partners share all their offspring. In the median population, mutation–selection–drift balance maintains about a twentieth of the same variance, at the mean deleterious effects estimated in related species and with the annotated functional sequence as the deleterious target. Both equilibria therefore leave most of what they are meant to explain to other processes. The question these estimates raise is how a variance measured over a few decades relates to the long-term variance that each equilibrium describes.

Our diversity test shows that the median reduction at unlinked sites lies within the 16 to 55% that Charlesworth and Jensen [35] obtained from the mean of the same estimates, using an approximation that holds while the variance is small [14, 17]. Because each tenfold loss of diversity requires the same increment of variance, the observed 230- to 38,000-fold deficits would require long-term variances above one, larger than any estimate. Most of each deficit is likely to come from processes that act over the many generations that neutral diversity reflects, which neither a pedigree nor a contemporary census records. Diversity reflects the harmonic mean of population size over the genealogy, and bottlenecks and range contractions hold that mean close to the smallest sizes, as Charlesworth and Jensen emphasize. A contemporary census therefore overstates the population size from which we computed the diversity expected without selection. Recurrent extinction and recolonization lower species-wide diversity [79] by the same mechanism through which heritable fitness differences increase drift, acting on demes rather than individuals, because a deme’s success, like an individual’s heritable advantage, persists across generations [80]. Because relatives share their deme, a pedigree within a single deme cannot measure this source of drift, which, like a bottleneck, raises the variance that the diversity test requires without implying that the estimate is too low. Demography and deme turnover therefore remain the expected causes of the missing diversity, and heritable variance in fitness adds a share to theirs rather than competing with them as an explanation.

Heritable variance in fitness reduces diversity at a neutral site in two ways, through the individual (or equivalently the diploid genome) that carries the site, an effect shared by every site, and through the chromosome segment that carries it, an effect that varies along the genome. Maps of background selection explain much of the variation in diversity along human chromosomes [28, 29] because they estimate the second effect relative to a genome-wide baseline that already includes the first [28, 81]. Such maps cannot measure the genome-wide effect, which accounts of the diversity deficit have presumed small [41], whereas a pedigree measures the variance in fitness among individuals on which that genome-wide effect depends. The same maps attribute all selected variance to recurrent deleterious mutation and, to match the depth of the reductions in diversity, fit deleterious mutation rates and selected targets higher than independent estimates support [28, 29]. Recurrent mutation maintains only a small share of the variance measured in wild pedigrees. The pattern of diversity along a chromosome, which depends on where selected sites lie and how fast selection removes them, can therefore follow mutation–selection–drift balance while the depth of the reduction requires variance from other sources. Along chromosomes, the reduction depends on what maintains the variance [24, 25, 82]. Our derivation shows that linkage adds to the loss of diversity only while the variance on the chromosome copy that carries a neutral site stays with it beyond the generation or two that an unlinked association lasts. The alleles that stay longest are those purged most slowly, which contribute the smallest share of the maintained variance, so the conditions for a large effect of linkage and for a large maintained variance exclude each other. At the measured variance, linkage reduces the effective size of every population with a map, yet in each population with diversity data predicted diversity still exceeds observed diversity at least 53-fold. Genomic data from these pedigreed populations can test how linkage acts, since diversity on each chromosome should decrease with the amount of selected sequence per unit of map length, along a single curve shared by macrochromosomes and microchromosomes.

Our maintenance test shows that, at the mean deleterious effects estimated in related species, lifetime fitness in wild vertebrates carries more additive variance than recurrent deleterious mutation can maintain.

Charlesworth [33] found the same for the fitness components of *Drosophila*, so the long-standing question of how quantitative genetic variation is maintained [34] applies to fitness itself. Because a new mutation adds variance in proportion to the square of its effect for a time inversely proportional to that effect, it contributes in proportion to its effect whatever its size. Only the deleterious mutation rate and the mean effect therefore set the variance that mutation maintains, and drift can only lower it. Site-frequency spectra constrain the mean effect more weakly than any other feature of a distribution of fitness effects, because rare mutations of large effect influence it. Yet large-effect mutations hidden from the spectra, such as transposable-element insertions, could raise it to the mean effect of about 16% that the median population requires only if they made up a large share of new mutations. Molecular estimates imply a high deleterious mutation rate in humans [83], recently put at 2 to 10 per generation [84] and above 4 once slightly deleterious mutations that fix are counted [85], with about 7–9% of the non-coding genome under purifying selection [78]. However, counting even 15% of the unannotated sequence of our assemblies as deleterious leaves the median population with four times the variance that mutation maintains, and in most bird populations the deleterious sequence required would be larger than the genome. When the target, the mean effect, and the estimate vary together, the excess persists in 12 of 18 populations if 8% of the unannotated sequence is deleterious, 1% of new mutations have large effects, and the estimate is twice the long-term variance, and in 6 under larger departures. The outcome is firm in a few populations, those with the largest estimates and birds with small mean effects, and provisional for the median population. A higher deleterious mutation rate also removes more diversity through background selection, yet even with the largest target we considered, the variance that mutation maintains removes at most about half of neutral diversity, so a mutational account narrows the maintenance gap without explaining the missing diversity.

To ask what maintains the remaining variance, we distinguish selection on phenotypes from selection on the alleles that underlie them. At the phenotypic level, additive variance in relative fitness measures directional selection, and it sets the rate of evolutionary change in every trait genetically correlated with fitness [5, 6, 86]. Under mutation–selection–drift balance, that directional selection consists only of the continual purging of deleterious alleles, so in the median population purifying selection accounts for about a twentieth of the directional selection on fitness at the mutational effects estimated in related species. Selection on breeding date shifts from year to year across 39 bird and mammal populations [87]. As a population lags behind a shifting optimum of this kind, standing variation in selected traits is continuously converted into additive variance in fitness without a larger mutational target [88, 89]. Under stabilizing selection toward a slowly moving optimum, new mutations that move the trait toward the displaced optimum are favored, so they rise to higher frequencies instead of being purged and amplify the genetic variance of the trait. Provided selection on individual alleles is strong relative to drift, this added variance depends much less on the mutation rate and the number of loci than the variance maintained under a fixed optimum [90]. Similarly, a single overdominant or negatively frequency-dependent locus displaced from its equilibrium by drift contributes as much additive variance in fitness as 100 to 10,000 loci at mutation–selection balance, particularly in small populations [36]. Migration–selection balance also adds variance whenever immigrant alleles adapted to neighboring habitats displace a local population from its optimum [91], as may occur in three of our populations, the Mediterranean blue tits, whose laying date and other heritable traits diverge between deciduous and evergreen oak patches a few kilometers apart despite persistent gene flow [77, 92]. In Fisher’s fundamental theorem, a shifting optimum, local maladaptation from gene flow, and environmental change all count as a deterioration of the environment that offsets the selective gain in mean fitness, which explains why a large heritable variance in fitness need not produce runaway population growth [9, 93, 94]. Yet this deterioration offsets only the change in mean fitness within a generation. It does not offset the drift that the same heritable differences in fitness cause over the far longer span that neutral diversity reflects. A pedigree cannot separate balancing selection from a moving optimum, but genomic and long-term phenotypic data can, because balancing selection maintains polymorphisms of large effect at intermediate frequency, whereas under a moving optimum breeding values shift while phenotypic stasis masks the change [7].

Shared environments, maternal effects, and indirect genetic effects that the animal model does not include can inflate pedigree estimates [95–98], whereas an estimate that misses episodes of intense selection would understate the variance. Because both tests rely on the same long-term variance, which removes more diversity the larger it is but is harder to maintain, correcting an inflated estimate would narrow the maintenance gap and widen the diversity gap by the same factor. Our threshold analysis shows that, within every population with diversity data, the long-term variance that the diversity test requires is at least 108 times the variance that the maintenance test allows, and at least 81 times with linkage. Every population therefore fails our compatibility test, the condition that a single long-term variance pass both tests, and no error that scales the estimate up or down changes that verdict. A variance that changes over time is not such an error, because the two tests respond to it differently. Mutation maintains only the long-term mean, and while the variance is small, diversity also depends only on that mean. If the effects of loci on fitness are multiplicative, however, generations of exceptionally intense selection remove more diversity than their share of the mean, whereas selection that reverses direction within a generation or two removes less, because an ancestor’s advantage then no longer accumulates through its descendants. Whether such generations remove a substantial share of neutral diversity is unknown, since no pedigree spans more than a few decades. If the estimates were overstated twentyfold, the factor that the median population requires to pass the maintenance test, our two results would reduce to one already known, that the variance mutation maintains removes a few percent of neutral diversity at sites unlinked to selected loci. Taken alone, the diversity test is passed by two populations, bighorn sheep and song sparrows, and then only with linkage, with one adult in ten breeding, and with the mutation rate at the lower end of its confidence interval. To reconcile the two tests, an assumption of unknown direction, such as discrete generations in iteroparous populations [99], a single deme per population, or distributions of fitness effects [40] and diversity taken from related species, would have to shift one result relative to the other by about two orders of magnitude. Because the usual criticisms of pedigree estimates describe errors that scale the estimate, the gap reflects the processes that the equilibrium models leave out more than it reflects error in the estimate.

Predicting the loss of diversity from the variance in fitness alone also requires that allele frequencies at selected loci change little while a neutral lineage remains associated with them [21], a condition met at mutation–selection–drift balance and while polygenic traits track a moving optimum. Sweeps of positively selected alleles violate the condition and cause more drift per unit of variance, but the slow rate of molecular substitution limits their cumulative effect. If the effects of loci on fitness are multiplicative, the predicted drift comes from rare ancestors of exceptionally high fitness, and the variances that the observed deficits would require reach 70 to 95% of the value at which a finite population contains fewer than one such ancestor per generation. As the variance approaches that value, lineages no longer coalesce steadily in pairs. Several lineages instead trace back at once to the same exceptional ancestor, so coalescence comes in bursts and the genealogy becomes a multiple-merger coalescent, the form that highly skewed offspring numbers, rapid adaptation, and occasional very large families produce [100–102]. Heritable variance in fitness large enough to explain the observed deficits would change the shape of the genealogy as well as its depth, and the diversity it leaves would no longer follow from an effective size alone. Balancing selection departs further from the condition, because a polymorphism at its equilibrium contributes no additive variance in fitness, so whatever drift it causes comes from displacements away from the equilibrium. Slight oscillations of balanced polymorphisms can reduce linked diversity over wide chromosomal regions, and seasonally fluctuating balancing selection at 100 loci reduces genome-wide effective size by 30 to 76%, largely independently of census size [103, 104]. Our conclusion that heritable variance in fitness is too small to explain the missing diversity therefore holds while purifying selection or a moving optimum maintains the variance, and whether fluctuating balanced polymorphisms remove more diversity than their contribution to that variance predicts remains the open question.

The compatibility test involves only the deleterious mutation rate, the mean effect of new mutations, the mating system, and the loss of diversity. The conflict therefore lies between what mutation can maintain and the diversity that is missing, and the condition applies unchanged to any future estimate of heritable variance in fitness. Temporal change in allele frequencies provides one such estimate, because its covariance across generations measures the heritable variance in fitness carried by linked selected loci [26, 27]. Replicate populations founded from a common base, as in long-term evolve-and-resequence experiments, come closer to the long-term variance that both tests concern. The change in allele frequency that replicates share over several generations gives the additive genetic covariance in fitness across replicates and generations, and not a variance in the environment of a single generation [105]. Pedigreed populations with genome data can also measure the reduction in effective size directly from the expected genetic contributions of ancestors to later generations [16, 106], although such a measure includes all sources of variance in reproductive success, of which heritable variance in fitness is one.

## Additional Information

### Data Availability

All inputs are published or derived from published data. Posterior draws of *V*_*A*_ from the model fits of Bonnet et al. [9] were provided by its lead author, and the phenotypic records and relatedness matrices are archived as their Data S1; genome assemblies and annotations are from NCBI (accessions in Table S3); mutation rates and nucleotide diversity are from Wang and Obbard [60], Lewin and Eyre-Walker [11] and the sources listed in Methods and Table S3; census sizes are from Buffalo [10]; and DFE parameters are from Lin et al. [40]. One data layer (scripts/R/00 prepare data.R) joins these sources through a single population crosswalk and writes the tables that every analysis takes as input. Analyses were run in R 4.6.0, with posterior modes and HPD intervals from MCMCglmm [107] and coda, and data handling and figures with dplyr, tidyr, readr, readxl and ggplot2. All code for the simulations, analysis and figure production, and the processed input tables, are available at https://github.com/mawassw/va-fitness-diversity; the posterior draws of *V*_*A*_ are not redistributed there and are available on request from the authors of Bonnet et al. [9].

### Author Contributions

WM led the development of the analytical framework. WM conducted the analysis of models and the generation of visuals. WM drafted the manuscript.

## Acknowledgments

We thank Timothée Bonnet for sharing the posterior distributions of the animal-model fits. We extend our thanks to members of the Berg Lab for their input on this project: Axel Zagal, Sharon Zhou, and Kellen Riall. We also thank members of the University of Chicago’s popgen supergroup for their comments and support.

## Notation

**Table S1:**
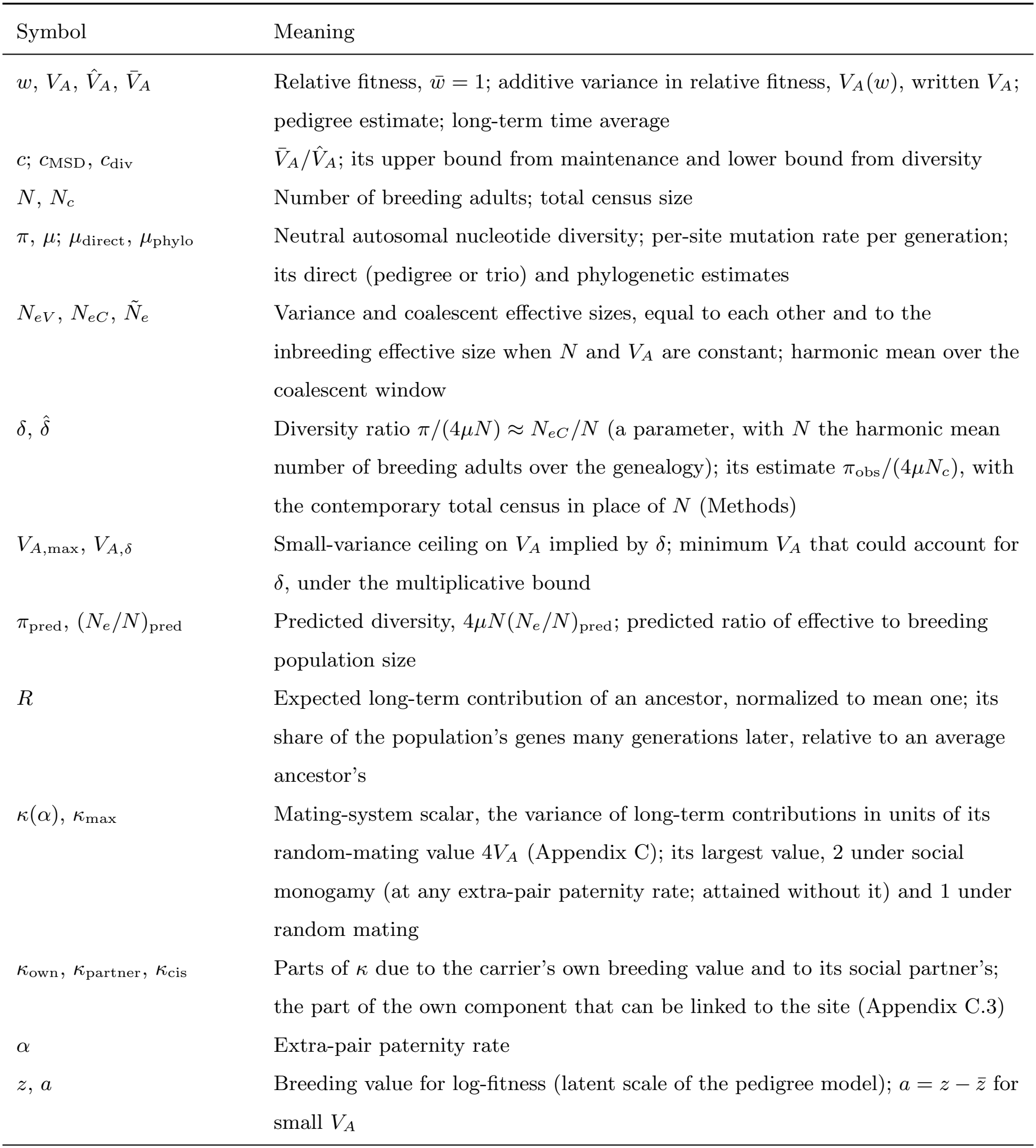
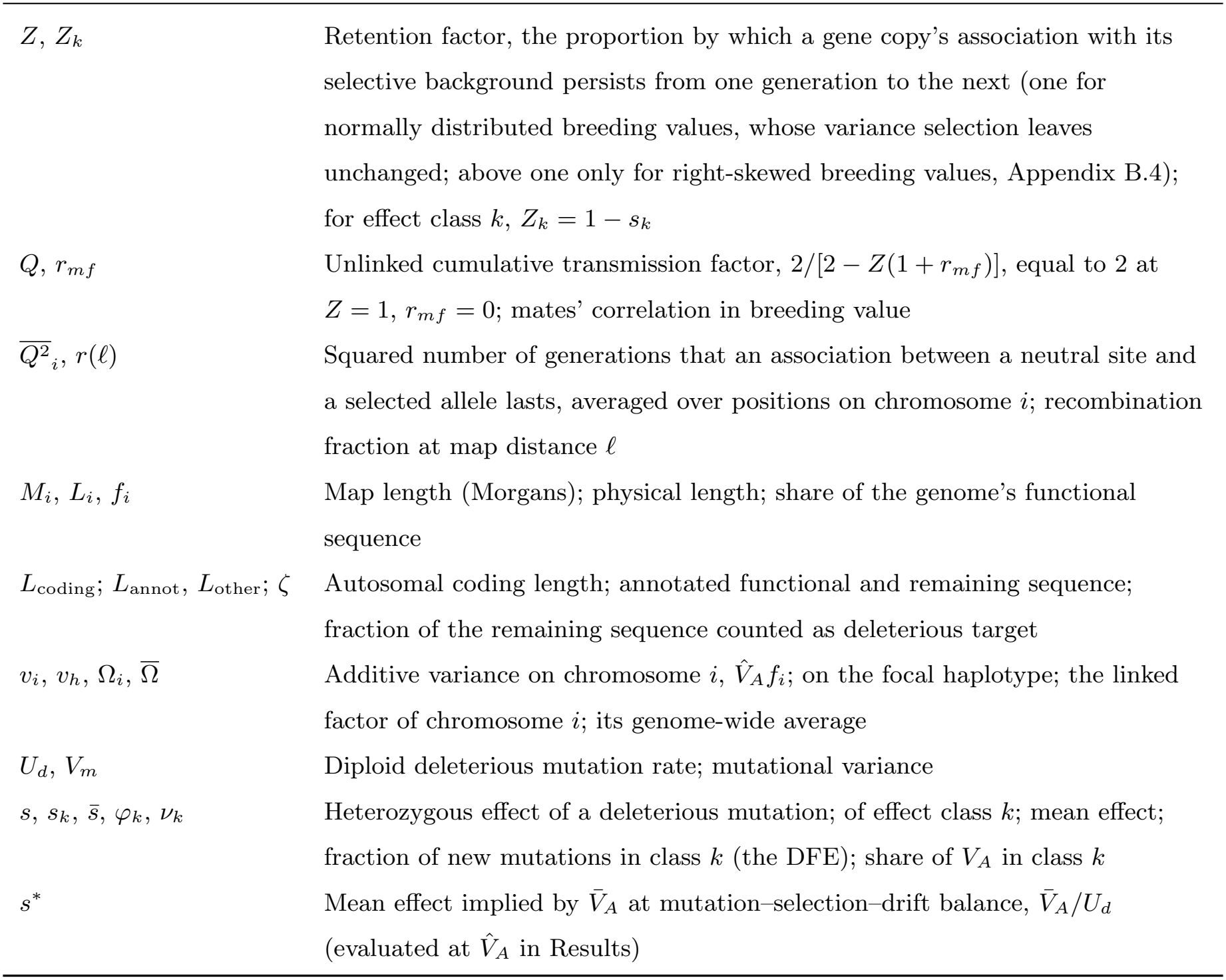
Notation used in the main text. Symbols local to a derivation in an appendix are defined where they appear there.

| Symbol | Meaning |
|---|---|
| w, V<sub>A</sub>, $\hat{V}_A$, $\bar{V}_A$ | Relative fitness, $\bar{w}$ = 1; additive variance in relative fitness, V<sub>A</sub>(w), written V<sub>A</sub>; pedigree estimate; long-term time average |
| c; c<sub>MSD</sub>, c<sub>div</sub> | $\bar{V}_A/\hat{V}_A$; its upper bound from maintenance and lower bound from diversity |
| N, N<sub>c</sub> | Number of breeding adults; total census size |
| π, μ; μ<sub>direct</sub>, μ<sub>phylo</sub> | Neutral autosomal nucleotide diversity; per-site mutation rate per generation; its direct (pedigree or trio) and phylogenetic estimates |
| N<sub>eV</sub>, N<sub>eC</sub>, $\tilde{N}_e$ | Variance and coalescent effective sizes, equal to each other and to the inbreeding effective size when N and V<sub>A</sub> are constant; harmonic mean over the coalescent window |
| δ, $\hat{\delta}$ | Diversity ratio π/(4μN) ≈ N<sub>eC</sub>/N (a parameter, with N the harmonic mean number of breeding adults over the genealogy); its estimate π<sub>obs</sub>/(4μN<sub>c</sub>), with the contemporary total census in place of N (Methods) |
| V<sub>A,max</sub>, V<sub>A,δ</sub> | Small-variance ceiling on V<sub>A</sub> implied by δ; minimum V<sub>A</sub> that could account for δ, under the multiplicative bound |
| π<sub>pred</sub>, (N<sub>e</sub>/N)<sub>pred</sub> | Predicted diversity, 4μN(N<sub>e</sub>/N)<sub>pred</sub>; predicted ratio of effective to breeding population size |
| R | Expected long-term contribution of an ancestor, normalized to mean one; its share of the population’s genes many generations later, relative to an average ancestor’s |
| κ(α), κ<sub>max</sub> | Mating-system scalar, the variance of long-term contributions in units of its random-mating value 4V<sub>A</sub> (Appendix C); its largest value, 2 under social monogamy (at any extra-pair paternity rate; attained without it) and 1 under random mating |
| κ<sub>own</sub>, κ<sub>partner</sub>, κ<sub>cis</sub> | Parts of κ due to the carrier’s own breeding value and to its social partner’s; the part of the own component that can be linked to the site (Appendix C.3) |
| α | Extra-pair paternity rate |
| z, a | Breeding value for log-fitness (latent scale of the pedigree model); a = z − $\bar{z}$ for small V<sub>A</sub> |
| Z, Z<sub>k</sub> | Retention factor, the proportion by which a gene copy’s association with its selective background persists from one generation to the next (one for normally distributed breeding values, whose variance selection leaves unchanged; above one only for right-skewed breeding values, Appendix B.4); for effect class k, Z<sub>k</sub> = 1 − s<sub>k</sub> |
| Q, r<sub>mf</sub> | Unlinked cumulative transmission factor, 2/[2 − Z(1 + r<sub>mf</sub>)], equal to 2 at Z = 1, r<sub>mf</sub> = 0; mates’ correlation in breeding value |
| $\overline{Q^2}_i$, r(ℓ) | Squared number of generations that an association between a neutral site and a selected allele lasts, averaged over positions on chromosome i; recombination fraction at map distance ℓ |
| M<sub>i</sub>, L<sub>i</sub>, f<sub>i</sub> | Map length (Morgans); physical length; share of the genome’s functional sequence |
| L<sub>coding</sub>; L<sub>annot</sub>, L<sub>other</sub>; ζ | Autosomal coding length; annotated functional and remaining sequence; fraction of the remaining sequence counted as deleterious target |
| v<sub>i</sub>, v<sub>h</sub>, Ω<sub>i</sub>, $\overline{\Omega}$ | Additive variance on chromosome i, $\hat{V}_A$f<sub>i</sub>; on the focal haplotype; the linked factor of chromosome i; its genome-wide average |
| U<sub>d</sub>, V<sub>m</sub> | Diploid deleterious mutation rate; mutational variance |
| s, s<sub>k</sub>, $\bar{s}$, φ<sub>k</sub>, ν<sub>k</sub> | Heterozygous effect of a deleterious mutation; of effect class k; mean effect; fraction of new mutations in class k (the DFE); share of V<sub>A</sub> in class k |
| s<sup>*</sup> | Mean effect implied by $\bar{V}_A$ at mutation–selection–drift balance, $\bar{V}_A$/U<sub>d</sub> (evaluated at $\hat{V}_A$ in Results) |

## Appendix A: Mutation–selection–drift balance and the implied selection coefficient

### A.1 Stationary variance

Suppose deleterious mutations arise at rate *U*_*d*_ per diploid genome per generation across *L*_target_ independent sites in a population of constant effective size *N*_*e*_. We define *U*_*d*_ = 2*μL*_target_, with per-site mutation rate *μ*satisfying 4*N*_*e*_*μ≪* 1. Mutations are semidominant with heterozygous effect *s ≪* 1 on relative fitness (genotype fitnesses 1, 1 *− s*, 1 *−* 2*s*), so a site at mutant frequency *q* contributes 2*q*(1 *− q*)*s*^2^ to the additive variance in relative fitness *V*_*A*_. Under a distribution of fitness effects (DFE) with density *φ*(*s*) and arithmetic mean 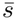, class *k* has heterozygous effect *s*_*k*_ and receives fraction *φ*_*k*_ of new mutations.

For a semidominant deleterious allele, the stationary expected heterozygosity per site (2*q*(1 *− q*)) under the diffusion approximation [46] is

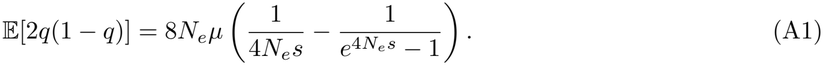

Multiplying by *s*^2^ and summing over *L*_target_ sites gives Equation (4) of the main text:

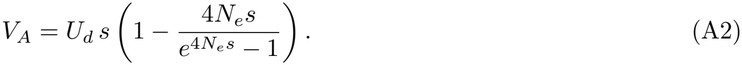

The factor in parentheses lies between 0 and 1. For *N*_*e*_*s ≪* 1, it expands to 2*N*_*e*_*s*, recovering the neutral mutation–drift variance *V*_*A*_ *≈* 2*N*_*e*_*U*_*d*_*s*^2^ [108]. For *N*_*e*_*s ≫* 1, it approaches 1, because each allele is held near *q* = *μ/s* and *V*_*A*_ *≈ U*_*d*_*s* at mutation–selection balance [22, 32].

### A.2 Inverting for *s*^*\**^

Because Equation (A2) is strictly increasing in *s* from 0 to *∞*, any observed *V*_*A*_ *>* 0 determines a unique single-effect solution *s*. In the strong-selection limit, Equation (5) inverts to 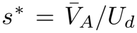 (Equation (6)), which is self-consistent whenever 2*N*_*e*_*s*^*\**^ *≫* 1. A solution in the weak-selection regime (2*N*_*e*_*s ≪* 1), where *V*_*A*_ *≈* 2*N*_*e*_*U*_*d*_*s*^2^, would require

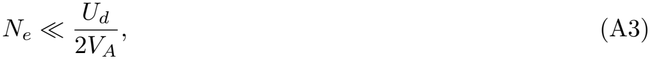

corresponding to an effective size of only a few individuals for *U*_*d*_ of order 1 and *V*_*A*_ of order 0.1. Both 2*N*_*e*_*s*^*\**^ and *U*_*d*_*/*(2*V*_*A*_) are reported for each study population in the Results.

### A.3 Distributions of fitness effects

Summing Equation (A2) over selective classes gives the stationary variance maintained under a DFE:

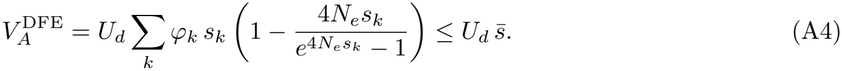

Equality is attained in the strong-selection limit, where each class contributes *U*_*d*_ *φ*_*k*_*s*_*k*_ regardless of the shape of *φ*(*s*), because a class of larger effect adds more variance per generation and is removed proportionally sooner (subsection 2.2); consequently, *s*^*\**^ estimates the arithmetic mean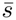. The shape of the DFE matters only through nearly neutral classes (4*N*_*e*_*s*_*k*_ ≲ 1), which contribute less than *U*_*d*_ *φ*_*k*_*s*_*k*_ and therefore lower 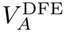 for a fixed mean 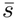. Using the upper bound 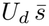 with the DFE assigned to each population [40] (Methods) is therefore biased toward passing the maintenance test, because it overstates the variance that mutation–selection–drift balance maintains. The class-specific retention factor *Z*_*k*_ = 1 *− s*_*k*_ and variance share 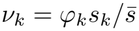 are derived in Appendix B.4.

### A.4 Unlinked reduction under fast purging

Alleles of large deleterious effect are purged within a few generations, so their associations with unlinked neutral sites decay faster than under Mendelian segregation alone. At retention factor *Z*_*k*_ = 1 *− s*_*k*_, the unlinked transmission factor (Equation (9)) becomes *Q*_*k*_ = 2*/*(1 + *s*_*k*_) in place of 2 [17, 22]. Weighting by each class’s variance share 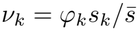 of 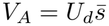 gives

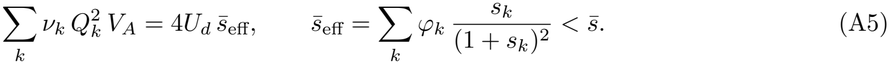

Large effects are discounted because their carriers are removed before the association has lasted its two generations. Equation (A5) replaces 4*κV*_*A*_ by 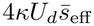 in the reduction at unlinked sites, in the multiplicative model or its small-variance approximation. The substitution is exact under random mating and under strict monogamy. With extra-pair paternity, *κ* itself depends on *Z* (Equation (C5)), and holding it at its value for *Z* = 1 understates the reduction in *N*_*e*_, by at most 2.3% for *s*_*k*_ *≤* 0.1. Like truncation of the upper tail of log-fitness (Appendix B.3), purging lowers the reduction in *N*_*e*_ below the multiplicative bound used in the main text. Results give the size of this purging correction at *s*^*\**^.

### A.5 Equivalence to the unlinked limit of background selection

Equation (A5) coincides with the unlinked limit of classical background selection. Consider a diploid selected locus with haploid (per-gamete) deleterious mutation rate *μ*_locus_ and heterozygous effect *s*. At mutation– selection balance (*q ≈ μ*_locus_*/s*), the locus contributes 2*qs*^2^ = 2*μ*_locus_*s* to *V*_*A*_, or *μ*_locus_*s* per homolog. For a neutral site at recombination fraction *r*, the background-selection reduction from the single *cis* copy is exp *−μ*_locus_*s/*(*s* + *r*(1 *− s*))^2^ [24, 25], which at 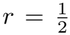 reduces to exp {*−*[2*/*(1 + *s*)]^2^*μ*_locus_*s*}, which gives the transmission factor *Q*_*k*_ = 2*/*(1 + *s*_*k*_) of Equation (A5).

At an unlinked locus, however, the *trans* (homologous) copy recombines away at the same rate 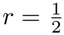 as the *cis* copy and makes an identical contribution, so the diploid locus lowers *N*_*e*_*/N* by exp *−*[2*/*(1 + *s*)]^2^ 2*μ*_locus_*s* . Summing over all loci with *U*_*d*_ = 2*μ*_locus_ yields the exponent 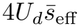 of Equation (A5) under random mating; summing over *cis* copies alone gives only half of the unlinked reduction. Equation (19) applies the same accounting within a chromosome. The *cis* half of the variance (*v*_*h*_ = *v*_*i*_*/*2) is assigned to the linked term and the homologous half stays in the unlinked term, whereas Buffalo and Kern [29] track only the *cis* half. Similarly, Marsh, Daigle, and Johri [50, Equation 4] evaluate the unlinked background-selection contribution from other chromosomes across uniform DFE bins by averaging ln(1 + *s*) + (1 + *s*)^*−*1^ over each bin. Because 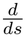 ln(1 + *s*) + (1 + *s*)^*−*1^ = *s/*(1 + *s*)^2^, their unlinked exponent equals 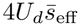 (both homologs included).

Their simulation result, that expressions derived for slow purging (*s ≪* 1) overestimate the loss of diversity at unlinked sites when purging is fast, is explained by the factor *Q*_*k*_ = 2*/*(1 + *s*_*k*_) derived above.

## Appendix B: Effective size under heritable variance in log-fitness

### B.1 Long-term genetic contributions and effective size

Following Barton and Etheridge [20], consider a diploid population of *N* breeding adults in which a parent with additive breeding value *z* for log-fitness produces a Poisson number of offspring with mean proportional to *e*^*z*^, where 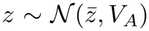. Dominance and epistatic deviations are omitted because they do not transmit stably across generations. Assume initially that mating is random with respect to *z*, and consider a neutral locus unlinked to all selected loci (pair bonds and extra-pair paternity are treated in Appendix C, and physical linkage in Appendix D).

Track a neutral gene copy carried by an ancestor with breeding value *z*. Tracking a gene copy rather than an individual allows the same bookkeeping to extend directly to pair bonds and linked sites; at an unlinked site the two perspectives are equivalent because a gene copy’s survival probability is proportional to the long-term genetic contribution of its carrier [20]. Under the infinitesimal model, offspring of carrier *z* inherit a mean deviation 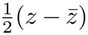 plus a Mendelian segregation residual of variance 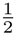. Because this segregation variance is homoscedastic (independent of *z*) and expected family sizes multiply along a lineage, deviations in each subsequent generation decay by a factor of 1*/*2 and sum on the log scale to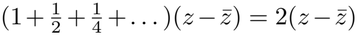.

The exact reproductive-value recursion [20, Appendix B] gives the expected long-term contribution of the ancestral copy, normalized so that E[*R*] = 1, as

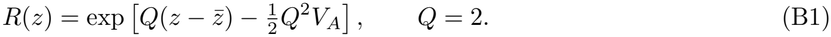

Two neutral copies sampled in a distant future generation coalesce in the ancestral generation if both descend from the same ancestral copy. Ancestral copy *j* (*j* = 1, …, 2*N* ), carried by an individual with breeding value *z*, contributes an expected fraction 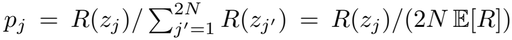 of the descendant gene pool. Conditional on the carrier’s breeding value, the two homologous copies within a diploid parent segregate independently into Poisson family sizes, so two sampled lineages trace to copy *j* with probability 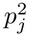 [20]. Summing over all 2*N* ancestral copies gives the per-generation rate of inbreeding [16],

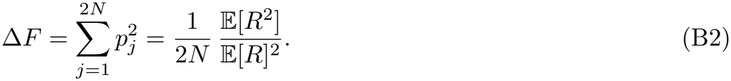

Without heritable variance, every copy has the same contribution, *R* = E[*R*], and Δ*F* = 1*/*(2*N* ), the Poisson baseline. Heritable variance replaces E[*R*]^2^ by E[*R*^2^], so the more unequal the shares *p*_*j*_, the more often two lineages trace to the same copy. Because Δ*F* is the probability that two lineages coalesce in one generation, setting Δ*F* = 1*/*(2*N*_*e*_) defines the inbreeding effective size, which here is also the coalescent effective size *N*_*eC*_ that *π* measures. The inbreeding, variance, and coalescent effective sizes coincide whenever the size and pattern of reproduction of a population remain constant, not only under Wright–Fisher sampling [109], and differ when its size changes [110]. The inbreeding effective size is not the variance effective size of a single generation. That size reflects only one generation’s family-size variance and so omits the persistence of the carriers’ advantage along the lineage that produces the factor *Q*^2^. With *N* and *V*_*A*_ constant, the long-term *N*_*eV*_ converges to the same value, and

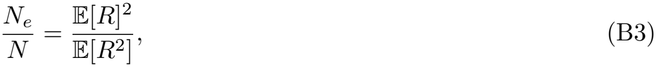

giving Equation (8) of the main text. Because log *R* is Gaussian, E[*R*]^2^*/*E[*R*^2^] = exp[*−*Var(log *R*)]. Under random mating, Var(log *R*) = *Q*^2^*V*_*A*_ = 4*V*_*A*_, yielding Equation (12):

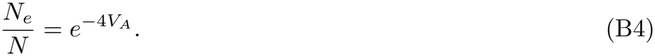

Under strict monogamy, pairs form at random and a pair with breeding values *z* and *z*′ produces offspring in proportion to 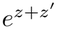 [23]. Every offspring that inherits the focal neutral copy from parent *z* simultaneously inherits half of its unlinked background from parent *z* and half from the monogamous partner *z*′. Consequently, the partner’s deviation 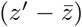 is co-transmitted along the focal gene’s lineage through the same geometric series 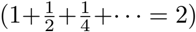 as the carrier’s own deviation, which gives log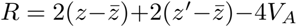. Because *z* and *z*′ are independent, Var(log *R*) = 8*V*_*A*_ and

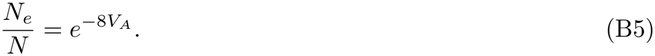

Equations (B4) and (B5) match the 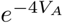 and 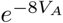 reductions in the fixation probability of a weakly beneficial allele derived by Weissman and Barton [23], because the coalescence rate increases, and the fixation probability decreases, in proportion to E[*R*^2^]*/*E[*R*]^2^ [111].

An ancestor’s expected contribution is settled within about ten generations, and mostly within the first two, because the initial terms of the *Q* series dominate [20]. The advantage therefore has to keep its sign for only a few generations to increase drift, as it does under MSDB, during sweeps of positively selected alleles, and under fluctuating selection that is positively autocorrelated between generations. Selection that reverses direction every generation instead makes the terms alternate in sign and gives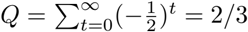. Over the genealogy, *V*_*A*_ and *N* may both change. If *N* (*t*) fluctuates, *N* in Equation (11) is replaced by its harmonic mean provided *V*_*A*_(*t*) and 1*/N* (*t*) are uncorrelated; if they covary, 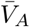 becomes the mean of *V*_*A*_(*t*) weighted by 1*/N* (*t*), because generations of small size contribute most to both the harmonic mean and the drift that heritable variance in fitness causes.

### B.2 The small-variance approximation

When *V*_*A*_ *≪* 1, linearizing 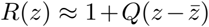 gives E[*R*^2^] = 1+Var(*R*) *≈* 1+*Q*^2^*V*_*A*_, reducing Equation (B3) to

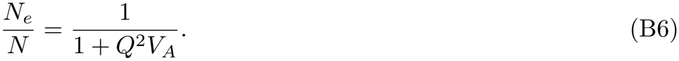

Robertson [2] and Santiago and Caballero [14] (reviewed in Wang et al. [112]) derived Equation (B6) for any smooth mapping from breeding value to fitness, because all such mappings reduce to the same linear form as *V*_*A*_*→*0. In their general formulation, the cumulative transmission factor is *Q* = 2*/*[2 *− Z*(1 + *r*_*mf*_ )] (Equation (9)), where *Z* is the retention factor (Appendix B.4) and *r*_*mf*_ is the correlation between mates’ breeding values. At *Z* = 1 and *r*_*mf*_ = 0, *Q* = 2 and *N*_*e*_*/N* = 1*/*(1 + 4*V*_*A*_); under strict monogamy, expanding Equation (B5) gives 1*/*(1 + 8*V*_*A*_).

Incorporating the general mating-system factor *κ*(*α*) (Equation (18)), setting *N*_*e*_*/N* = *δ*, and inverting the small-variance form yields the small-variance ceiling

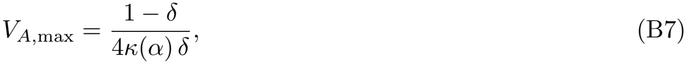

above which heritable variance in fitness alone would remove more diversity than observed. Because the small-variance expansion holds only for *V*_*A*,max_ *≤* 1, Equation (B7) applies only when *δ* ≳ 1*/*(1 + 4*κ*), that is, *δ* ≳ 0.20 under random mating (*κ* = 1), 0.12–0.16 under social monogamy with extra-pair paternity (*κ*_*B*_ = 1.32–1.86), and 0.11 under strict monogamy (*κ* = 2). For larger deficits, the required variance is *V*_*A,δ*_, from the multiplicative bound (Equation (14)).

At each population’s *κ*, none of the 19 pedigree estimates exceeds this ceiling at *δ ≤* 0.2; two do at *δ* = 0.25–0.3 (the blue tits at Pirio and Muro), five at *δ* = 0.5 and 15 at *δ* = 0.9 (Figure S2). Every 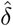 in the deficit set lies two to four and a half orders of magnitude below the *δ* at which its own estimate would reach the ceiling. Neutral diversity in these populations therefore sets no upper limit on their heritable variance in fitness.

### B.3 The upper tail of log-fitness and finite-population truncation

Under random mating, E[*R*^2^] *∝* E[*e*^4*z*^]. Exponentially tilting the Gaussian density of *z* by *e*^4*z*^ shifts its mean to 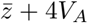 without altering its variance, so the ancestors dominating 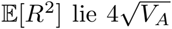 standard deviations above the population mean. A 10^2^-fold reduction in *N*_*e*_ (*V*_*A*_ = ln(100)*/*4 = 1.15) comes from ancestors 4.3 standard deviations above the mean (roughly 1 in 10^5^ individuals), whereas a 10^4^-fold reduction (*V*_*A*_ = 2.30) requires ancestors at 6.1 standard deviations (roughly 1 in 10^9^).

Realizing the analytical expectation E[*R*^2^] therefore requires both an untruncated biological upper tail and a census size large enough to sample that tail. Ecological limits on reproductive output, such as territory saturation, clutch-size constraints, or the upper limit of one on the probability of breeding in the zero-inflation component of the animal model [9] (subsection 2.4), truncate the tilted density and lower E[*R*^2^]. Consequently, Equations (B4), (B5), and (C12) give the largest reduction in *N*_*e*_ that a given *V*_*A*_ can cause.

Finite population size sets a second limit. For a general mating structure with Var(log *R*) = 4*κ*_max_*V*_*A*_, the lineages dominating E[*R*^2^] originate from ancestors 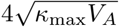 standard deviations above the mean. To leading exponential order in the Gaussian tail, such individuals make up a fraction 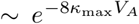 of the cohort. A population of *N* breeding adults (or *N/*2 monogamous pairs) samples these ancestors in a typical generation only while

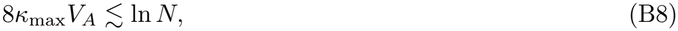

at which limit 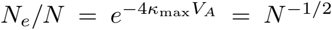. Beyond this limit, most generations contain none of the ancestors that E[*R*^2^] depends on. The analytical expectation is then reached only as an average over very many generations, a few of which contain such an ancestor, and lineages coalesce in bursts in those ancestors and not steadily in pairs [100–102]. For example, in single-generation Monte Carlo draws of *N* = 10^6^ random-mating individuals (where the limit is *V*_*A*_ = ln(10^6^)*/*8 = 1.73), the median realized pairwise coalescence rate at *V*_*A*_ = 1.5 is only 0.56 of the analytical expectation (range 0.36–5.5*×*), and 0.39 at *V*_*A*_ = 1.73, where the single fittest individual accounts for *∼* 1% of all long-term genetic contributions.

### B.4 The retention factor *Z*

In an ancestor-centered perspective, the retention factor *Z* is the additive genetic variance among successfully reproducing parents (weighted by their offspring output) divided by the cohort variance *V*_*A*_. In a gene-centered perspective, *Z* is the per-generation retention of a neutral allele’s statistical association with its selected background (the factor *G* of Santiago and Caballero [14]). Defining a stationary retention factor *Z* requires that allele frequencies at selected loci change negligibly over the timescale on which the background association decays [21, 113], as holds at mutation–selection–drift balance.

Under multiplicative fitness, weighting parents by their expected offspring *e*^*z*^ applies an exponential tilt with parameter *ψ* = 1 to the distribution of *z*. Letting *K*(*ψ*) = ln E[*e*^*ψz*^] denote the cumulant-generating function of *z*, the variance of the offspring-weighted parent distribution is *K*^*′′*^(1), while *V*_*A*_ = *K*^*′′*^(0), giving

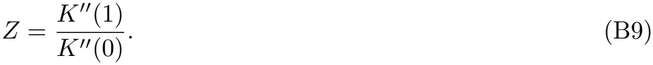

When *z* is strictly Gaussian, 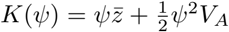, so *K*^*′′*^(*ψ*) = *V*_*A*_ for all *ψ* and *Z* = 1 identically, because exponential selection shifts the parental mean by *V*_*A*_ while leaving the variance unchanged. Expanding *K*^*′′*^(1) around *ψ* = 0 for an arbitrary distribution gives

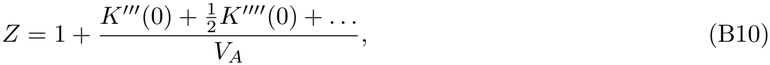

showing that positive skewness in breeding values (*K*^*′′′*^(0) *>* 0) raises *Z* above 1 (subsection 2.7).

Now suppose *V*_*A*_ is maintained by rare deleterious alleles at mutation–selection balance. A heterozygous carrier of an allele in selective class *k* has relative fitness 1 *− s*_*k*_, a log-fitness deviation ln(1 *− s*_*k*_), and equilibrium frequency *q*_*k*_ *≈ μ/s*_*k*_. Because cumulants sum across unlinked loci, a rare locus in class *k* Contributes 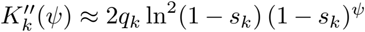. Applying Equation (B9) across classes to leading order in *s*_*k*_ gives

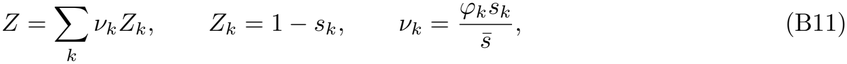

where *φ*_*k*_ is the proportion of new deleterious mutations in class *k* and *ν*_*k*_ is that class’s share of total *V*_*A*_. Each mutational class is retained in proportion to its carriers’ relative survival 1 *− s*_*k*_ (Equation (10)). While the class-weighted average reproduces the aggregate retention factor 1 *−* E[*s*^2^]*/*E[*s*] of Santiago and Caballero [17], the main text evaluates *Q* and 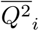 separately by class because both quantities are nonlinear in *Z*_*k*_. A larger effect *s*_*k*_ speeds purging, which lowers *Z*_*k*_, *Q*_*k*_, and the resulting reduction in *N*_*e*_ below the Gaussian (*Z* = 1) limit; by contrast, rare beneficial alleles (*s*_*k*_ *<* 0) yield *Z*_*k*_ *>* 1.

Finally, Equation (B9) relies on the exponential map *w ∝ e*^*z*^. Under a linear map *w* = 1+*a* with E[*a*] = 0 and Var(*a*) = *V*_*A*_, the offspring-weighted parental variance is 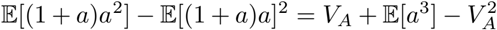, giving

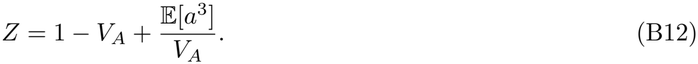

For a Gaussian breeding value (E[*a*^3^] = 0), *Z* = 1*−V*_*A*_, *Q* = 2*/*(1+*V*_*A*_), and 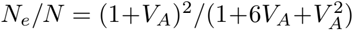. This ratio reaches a minimum of 1*/*2 at *V*_*A*_ = 1 and increases toward 1 at larger *V*_*A*_. That increase is an artifact of the linear map, which assigns negative expected offspring (1 + *a <* 0) to a fraction Pr(*a < −*1) of the cohort (8% of individuals at *V*_*A*_ = 0.5 and 16% at *V*_*A*_ = 1); we therefore do not consider the linear fitness map further.

## Appendix C: Effective size with separate sexes, pair bonds, and extra-pair paternity

This appendix derives the mating-system scalar *κ* of Equation (18) (Model B, Equation (C6)), its split into the components used in Equation (19), and the multiplicative-model bracket of Equation (C12). It applies the expected long-term contributions of Woolliams, Bijma, and Villanueva [19] and the two-sex combination of Santiago and Caballero [14].

Consider a diploid population with discrete generations, *N* breeding adults, and an equal sex ratio. Each individual has an additive breeding value *a* for relative lifetime breeding success, with variance Var(*a*) = *V*_*A*_ assumed equal in both sexes. For small advantages, 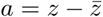 (Appendix B). Bonnet et al. [9] estimate this single variance from an animal model that includes both sexes with sex fitted as a fixed effect.

Under random mating, a parent’s relative family size is 1 + *a*. Under social monogamy, the nest is the unit of productivity. A female paired with social partner *m*′ has relative family size 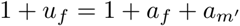. Extra-pair paternity at rate *α* moves paternity out of the nest without altering its overall productivity, so a male paired with social partner *f*′ has relative success

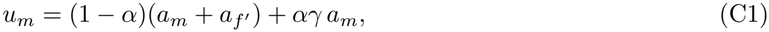

where *γ* indexes the dependence of extra-pair siring success on the male’s own breeding value. Model A sets *γ* = 0 (extra-pair siring as a random lottery), whereas Model B sets *γ* = 1 (extra-pair gains exactly offset within-pair losses). Model B is the unique parameterization in which a male’s breeding value contributes to his total fitness with the same average weight as a female’s across all *α*, satisfying the single-variance assumption of the animal model. We therefore use Model B for all primary results.

A carrier’s own breeding value therefore contributes to its advantage with weight 1 in females and *β* = 1 *− α* + *αγ* in males, averaging 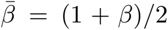 across the sexes. Social partners’ breeding values are correlated by *r*_*mf*_ . Each generation of selection multiplies a gene copy’s association with its selected background by the retention factor *Z* (Appendix B.4). Except in Section C.4, the following derivations use the small-variance approximation (*V*_*A*_ *≪* 1).

### C.1 Long-term contributions and *κ*

Consider a neutral gene copy in individual *X* with social partner *X*′. The copy’s expected number of transmitted descendants is proportional to 1 + *u*_*X*_ . In a descendant carrier with expected breeding value *h*, the partner’s expected breeding value is *r*_*mf*_ *h*, contributing to the carrier’s advantage with weight 1 in female nests and 1 *− α* in male nests. The carrier’s expected advantage is thus 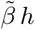, where

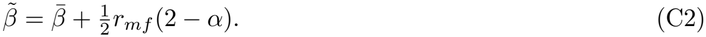

Offspring that inherit the copy receive half of *h* and half of the other parent’s expected breeding value. The other parent’s value is *r*_*mf*_ *h* when that parent is the social partner, with probability 1 *− α*, and zero when it is an extra-pair parent. Because selection then multiplies the association by *Z*, expected breeding values decline geometrically, *h*_*t*+1_ = *θh*_*t*_ with *θ* = (*Z/*2)[1 + (1 *− α*)*r*_*mf*_ ], the fraction of a carrier’s expected breeding value that the next carrier keeps (half by segregation, raised by a correlated partner, times the retention factor). In the first generation the partner’s breeding value is known,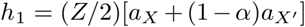. Summing 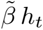 over *t ≥* 1 gives the cumulative advantage of the copy, 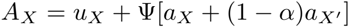 with 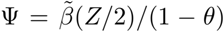, the descendants’ part of the cumulative advantage, so that 1 + Ψ = *Q* under strict monogamy and, at *r*_*mf*_ = 0, under random mating:

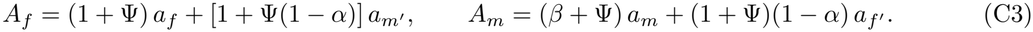

For small *V*_*A*_, 1 + *A*_*X*_ approximates the expected long-term contribution *R* (Equation (B1)), which is the regression of contribution on ancestral selective advantages [19, Equation 6b]. In each sex, E[*R*^2^] = 1 + Var(*A*), which gives *N*_*e,X*_ */N* = 1*/*[1 + Var(*A*_*X*_ )]. Combining sexes via Santiago and Caballero’s [14] formula 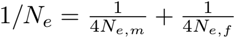 (following Crow and Denniston [110]) at an equal sex ratio makes *N/N*_*e*_ the mean of 1 + Var(*A*) across sexes:

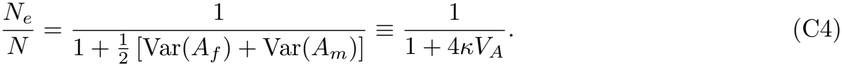

Here 4*κV*_*A*_ is the sex-averaged variance of long-term contributions, and *κ* scales this variance relative to its random-mating baseline (*Z* = 1, *r*_*mf*_ = 0, 4*V*_*A*_). For a weighted sum *ξ*_1_*a* + *ξ*_2_*a*′ of partner breeding values with correlation *r*_*mf*_, the variance is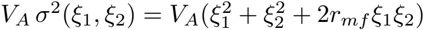, yielding

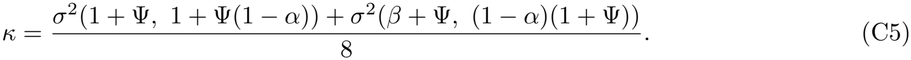

### C.2 *κ* under Models A and B

When *Z* = 1 and *r*_*mf*_ = 0, *θ* = 1*/*2 and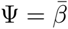. Model B 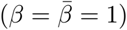 reduces Equation (C5) to

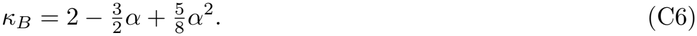

For Model 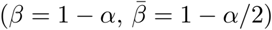,

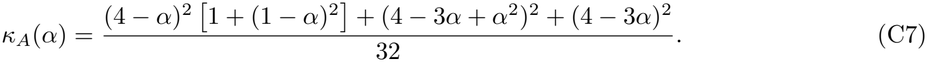

As *α* increases to 1, *κ*_*B*_ decreases monotonically to 9*/*8 (above 1 because a female’s family size still depends on her social partner), whereas *κ*_*A*_ decreases to 7*/*16 and is below 1 once *α >* 0.44 (Figure S3).

Under strict monogamy (*α* = 0), both models yield 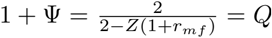, giving 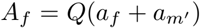 and recovering the pair-mated model of Santiago and Caballero [14] with *κ* = 2 at *Z* = 1, *r*_*mf*_ = 0. Under random mating, *u* = *a* in both sexes 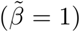, so that 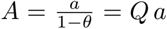 and *κ* = *Q*^2^*/*4 (1 at *Z* = 1, *r*_*mf*_ = 0).

Positive assortative mating (*r*_*mf*_ *>* 0) lengthens the association through *θ* and so raises *κ*; we set *r*_*mf*_ = 0 in the absence of empirical estimates. Under random mating and Model B, carrier weights equal 1 in both sexes, preserving the single-sex retention factor *Z* = 1 (Gaussian) or *Z*_*k*_ = 1 *− s*_*k*_ (deleterious classes;Equation (B11)).

### C.3 Own, partner, and cis components

At linked sites, *κ* must be split into the part due to the carrier and the part due to its social partner, because the ancestor-centered and gene-centered perspectives no longer coincide there (subsection 2.3). Neither the homologous (*trans*) haplotype nor a social partner’s breeding value co-segregates with the focal gene copy, so the individual ancestor’s long-term contribution no longer sets drift at linked sites. Because meiosis separates the homologous chromosomes every generation, a neutral allele remains linked only to the selected alleles in *cis* on its own haplotype [29], which carries half the chromosome’s diploid variance (*v*_*h*_ = *v*_*i*_*/*2). Similarly, only the carrier’s own breeding value, not its mate’s, acts in *cis*. Setting *r*_*mf*_ = 0, Var(*A*_*f*_ ) + Var(*A*_*m*_) separates into carrier and partner variances:

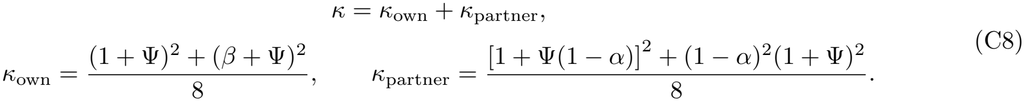

The partner’s genes are never on the focal copy’s haplotype, so the partner component is unlinked. The share of the own component that lies on the focal haplotype, *v*_*h*_, is assigned to the linked term. In the linked term, a selected locus keeps its association with the copy with probability *ρ* = (1 *− r*)*Z* per generation. When every carrier has weight 1, the cumulative advantage per unit of the locus’s effect is the linked kernel 1*/*(1 *−ρ*) = 1 + *χ* of subsubsection 2.6.2, with *χ* = *ρ/*(1 *−ρ*). With the carrier’s own weight *β*_*X*_ (1 in females, *β* in males) and later carriers’ average weight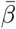, the cumulative advantage is 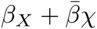 instead. The square of this weighted kernel relative to the square of the unweighted one, averaged over the sexes, is

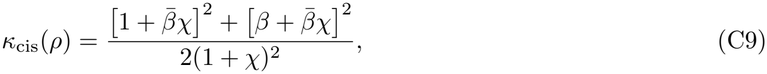

which decreases from (1 + *β*^2^)*/*2 as *ρ→*0 to 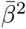 under tight linkage. Under random mating, and under Model B, *β* = 1, so *κ*_own_ = *κ*_cis_ = 1 and *κ*_partner_ = *κ −* 1. The remainder of the genome (the homologous haplotype and all other chromosomes) assorts independently of the focal neutral site, so Equation (19) uses this split, applying *κ*_cis_ to the linked term and *Q*^2^(*κV*_*A*_ *− κ*_own_*v*_*h*_) to the exponent of the unlinked background, with *Q* = 2*/*(2 *− Z*) at the same retention factor as the linked term. Because the *cis* haplotype variance *v*_*h*_ moves from the unlinked to the linked term, no variance is counted twice. The two terms combine additively in the exponent if selected loci are distributed independently along the genetic map, so that their effects multiply [24, 114], and fitness is multiplicative across loci [29, Supplementary Equations 28–29] (cf. [17]). Across a DFE, 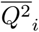 and *Q*^2^ are replaced by their averages over the variance weights *ν*_*k*_, and the multiplicative bound replaces *κ* by *κ*_max_ (Appendix D.5 gives the small-variance form).

### C.4 The multiplicative model with pair bonds

Under Model B with multiplicative fitness (*w ∝ e*^*z*^), a gene copy’s long-term contribution arises through within-pair and extra-pair routes. In females, productivity is 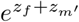, with a fraction 1 *− α* of offspring sired by *m*′ and *α* by extra-pair sires. In males, contributions arise from within-nest and extra-pair offspring proportional to 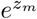. Following Appendix B.1, expected contributions are

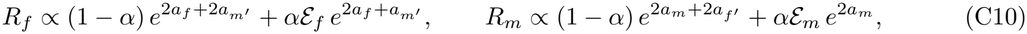

where *ℰ*_*f*_ and *ℰ*_*m*_ collect expectations over outside individuals and equal 1 at *V*_*A*_ = 0. Letting 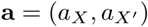 denote independent carrier and partner deviations, contributions take the mixture form 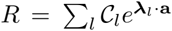, with exponent vectors ***λ*** *∈ {*(2, 2), (2, 1)*}* in females and *{*(2, 2), (2, 0)*}* in males, and coefficients *C*_1_ = 1 *− α, C*_2_ = *α ℰ*_*X*_ . Using the Gaussian moment-generating function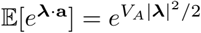,

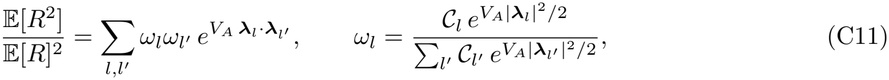

where *ω*_*l*_ is route *l*’s share of expected long-term contribution. Averaging over sexes per Equation (B3) gives *N*_*e*_*/N* . Because all dot products 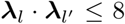, the ratio in Equation (C11) is bounded above by 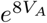 in either sex. From below, Jensen’s inequality bounds the sex mean by exp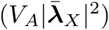. At baseline weights 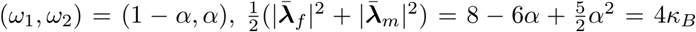. Consequently, provided extra-pair routes carry no more than their proportional share *α*,

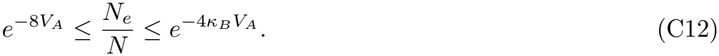

This bound holds for extra-pair multipliers *ℰ*_*f*_, *ℰ*_*m*_ up to 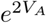. At *α* = 0, *κ*_*B*_ = 2 and both bounds collapse to Weissman and Barton’s [23] strict-monogamy result (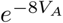; Equation (B5)). For small *V*_*A*_, the upper bound reduces to 1*/*(1 + 4*κ*_*B*_*V*_*A*_), whereas for larger *V*_*A*_ the within-pair route dominates E[*R*^2^], shifting the exact ratio toward the lower bound.

## Appendix D: The linked factor along a Haldane map

This appendix derives the chromosome-wide average 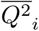 of subsubsection 2.6.2 (Equation (D4)) and evaluates it.

### D.1 The kernel and chromosome average

Consider a single mutational effect class with retention factor *Z* (Equation (10)); under a distribution of fitness effects, each expression below is averaged over classes using the variance weights *ν*_*k*_ of subsubsection 2.6.2. Take a chromosome of map length *M* Morgans (dropping the chromosome index *i* for clarity) on which two sites separated by *ℓ* Morgans recombine with probability *r*(*ℓ*). Unless stated otherwise (Section D.6), both selected variance and neutral sites are distributed uniformly in map distance. Let *Q*^2^ = [2*/*(2 *− Z*)]^2^ denote the unlinked transmission factor squared (Equation (9)), 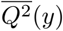 the linked value at a neutral site at position *y* averaged over selected loci, and 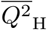 and 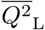 the chromosome-wide averages under the Haldane and linear maps (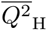 is the 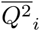 of the main text).

An association between a neutral site and a selected locus *ℓ* Morgans away survives one generation with probability (1*−r*(*ℓ*))*Z*, as recombination breaks it with probability *r*(*ℓ*) and selection multiplies the remaining association by *Z*. Summing over generations gives the cumulative advantage per unit effect [29, 114],

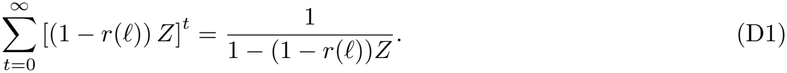

It is the counterpart of *Q* for the chromosome segment that carries the neutral site, counting the advantage over every generation in which the segment keeps the selected locus, so it is the expected number of generations that the association lasts (subsubsection 2.6.2). Because the cumulative factor appears squared in the variance of the segment’s long-term contributions, as *Q* does in Equation (B6), a locus at distance *ℓ* contributes *K*(*ℓ*) = [1 *−* (1 *− r*(*ℓ*))*Z*]^*−*2^. Here and below, *y* is always the position of a neutral site and *x* that of a selected locus. With 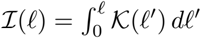, the profile and its chromosome average are

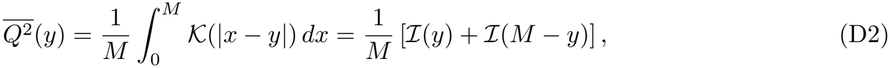

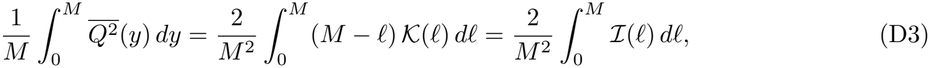

where the factor (*M −ℓ*) weights each distance *ℓ* by the density of site pairs separated by *ℓ*. Under Haldane’s mapping function, 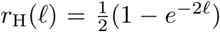, which saturates at 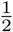 for sites far apart on the same chromosome, Equation (D3) becomes, for chromosome *i* of map length *M*_*i*_,

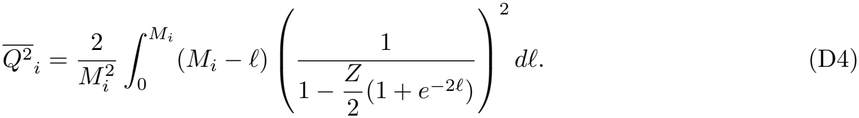

Substituting *Z* = 1 *− s* reduces the term inside the integral to 1*/*[*s* + *r*(*ℓ*)(1 *− s*)], the familiar kernel from classical background selection theory [17, 24, 25, 29, 114]. Whereas Buffalo and Kern [29] evaluate this kernel locally along a high-resolution recombination map, our spatial average yields the expected chromosome-wide reduction when only total map length *M*_*i*_ is known. Santiago and Caballero [17] and Buffalo and Kern [29, Supplementary Equations 32–33] derived similar chromosome-wide averages for a central or end site under a linear map. Equation (D4) uses Haldane’s mapping function and also averages over the position of the neutral site.

### D.2 Closed form under the Haldane map

Under Haldane’s map, 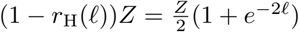, so

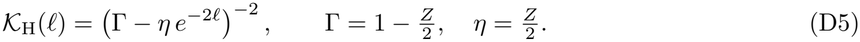

Substituting *τ* = *e*^*−*2*ℓ*^ and decomposing into partial fractions gives the arm integral in elementary functions:

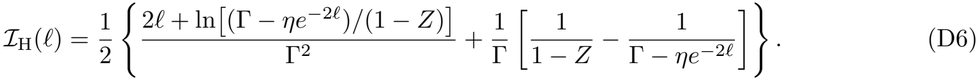

Integrating once more over [0, *M* ], using 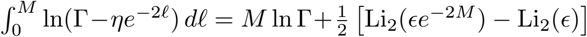 with *ϵ* = *η/*Γ and Li_2_ the dilogarithm, gives

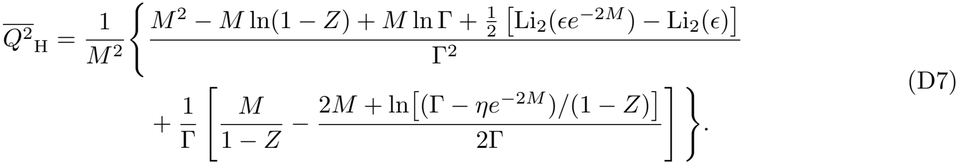

Equation (D7) is the exact form evaluated in the main analysis. It agrees with direct numerical quadrature of Equation (D3) in R to better than 10^*−*11^ relative error across *Z ∈* [10^*−*9^, 1 *−* 10^*−*9^] and *M ∈* [0.005, 60] Morgans, spanning all selection classes and map lengths analyzed.

### D.3 Short- and long-map limits

As 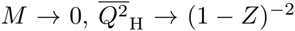, the non-recombining limit of background selection. Approaching this limit requires recombination across the entire chromosome to be slower than purging (*M ≪* 1*−Z*), not only a short physical map. At *Z* = 0.99 and *M* = 0.5, 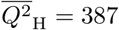 compared with (1 *− Z*)^*−*2^ = 10^4^. At the other extreme, as *ℓ→∞, K*_H_(*ℓ*)→Γ^*−*2^ = *Q*^2^, recovering the unlinked factor, and on a freely recombining chromosome (*M→∞*) the linked and unlinked terms of Equation (19) combine into the reduction at unlinked sites, 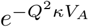. Convergence to this free-recombination limit is slow:

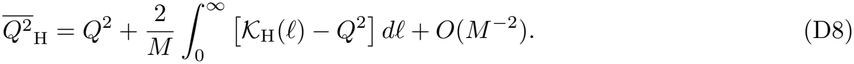

From Equation (D6), the leading integral evaluates to 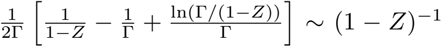 as *Z→*1 (10.3 at *Z* = 0.9 and 105 at *Z* = 0.99). Free recombination therefore requires *M ≫* 1*/*(1 *− Z*); even at *M* = 10 and *Z* = 0.99, 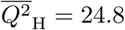 against *Q*^2^ = 3.9.

Empirical chromosomes occupy the intermediate regime 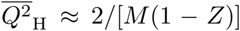 [17], because an allele removed at rate *s* = 1 *− Z* stays linked to sites within about *s* Morgans on either side, a fraction 2*s/M* of the chromosome, for about 1*/s* generations, which gives (1*/s*)^2^ *×* 2*s/M* . For the same reason the reduction depends on the selected target per Morgan, *f*_*i*_*/M*_*i*_. Even avian microchromosomes rarely approach the non-recombining limit, because an obligate crossover per bivalent implies *M* ≳ 0.5 Morgans, and the shortest empirical maps analyzed here (0.04–0.2 Morgans in great tits and song sparrows) still greatly exceed 1 *− Z* for all but the classes purged fastest (Table S4).

### D.4 The linear map as a lower bound

Under a linear map, *r*_L_(*ℓ*) = *ℓ*, the kernel is *K*_L_(*ℓ*) = (1 *− Z* + *ℓZ*)^*−*2^ and the arm integral is *ℐ*_L_(*ℓ*) = *ℓ/*[(1 *− Z*)(1 *− Z* + *ℓZ*)], whose mid-chromosome value matches Supplementary Equation 33 of Buffalo and Kern [29]. Equation (D3) gives

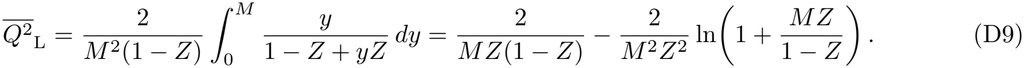

As 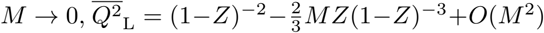, matching 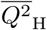 to first order because *r*_H_(*ℓ*) = *ℓ*+*O*(*ℓ*^2^). As *M→∞*, however, 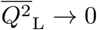 rather than *Q*^2^, because *r*_L_(*ℓ*) exceeds 1*/*2 for *ℓ >* 1*/*2.

Because *e*^*−*2*ℓ*^ *>* 1 *−* 2*ℓ* implies *r*_H_(*ℓ*) *< ℓ* for all *ℓ >* 0, it follows that 0 *<* 1 *− Z* + *r*_H_(*ℓ*)*Z <* 1 *− Z* + *ℓZ* everywhere for *Z ∈* (0, 1), so *K*_L_(*ℓ*) *< K*_H_(*ℓ*) at every distance and

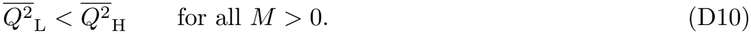

Equation (D9) is therefore a lower bound on 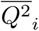, and hence on the reduction in *N*_*e*_ at linked sites. The gap grows with *M* and with 1 *− Z*; the ratio 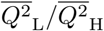 is 0.98, 0.95, and 0.91 at *M* = 0.1, 1, and 3 Morgans for *Z* = 0.99, and 0.99, 0.85, and 0.67 for *Z* = 0.9. Buffalo and Kern [29] use instead the linear-map profile at the middle of the segment, 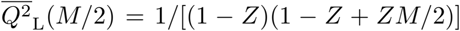, which is the largest value along the segment. The mid-segment value exceeds 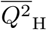 by at most about 12%, on short maps with *M* a few times 1 *− Z*, and is below it once *M* exceeds about one Morgan and purging is fast, to 0.81 at *M* = 10 for *Z* = 0.99 and 0.41 for *Z* = 0.9 (Figure S11). Both linear-map expressions are close to the Haldane average over the rest of the range, which includes the short, densely mapped segments of human data for which the mid-segment value was designed. The main text uses the Haldane closed form, Equation (D7).

### D.5 Upper bound on the linked factor (Ω_*i*_ *≤* 1)

Linkage can only lengthen the association between a neutral site and the selected alleles on its haplotype, so their variance never counts for less than it would at an unlinked site. The argument below makes this exact. Because *K*_H_(*ℓ*) decreases monotonically from (1 *− Z*)^*−*2^ to *Q*^2^, every local profile satisfies 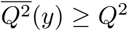, and therefore 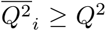 regardless of how selected variance is distributed along the chromosome (Section D.6). We define three compound quantities

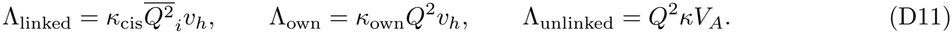

In the primary model *κ*_cis_ = *κ*_own_ (Appendix C.3), so Λ_linked_ *≥* Λ_own_; furthermore, Λ_unlinked_ *≥* Λ_own_ since *v*_*h*_ *≤ V*_*A*_ and *κ ≥ κ*_own_. Under the multiplicative form of Equation (19), 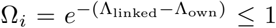 follows immediately, with equality on a freely recombining chromosome 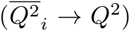, because there no association outlasts an unlinked one, so linkage adds nothing.

The bound also holds when the unlinked factor uses the small-variance reciprocal form 1*/*(1 +Λ_unlinked_ *−* Λ_own_), because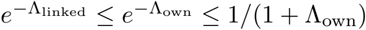,

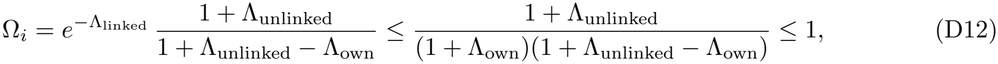

where the final inequality holds because (1 +Λ_own_)(1 +Λ_unlinked_ *−* Λ_own_) *−* (1 +Λ_unlinked_) = Λ_own_(Λ_unlinked_ *−* Λ_own_) *≥* 0. Under free recombination, this mixed form gives 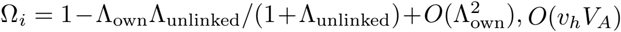, an *O*(*v*_*h*_*V*_*A*_) discrepancy arising solely from combining exponential and reciprocal approximations for the same variance. Under a distribution of fitness effects, the bound applies class by class.

With linkage, the minimum variance that accounts for *δ* solves Equation (21) set equal to *δ* (Methods), and because 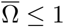 it lies below the unlinked value at the same retention factor, ln(1*/δ*)*/*(*Q*^2^*κ*), which equals *V*_*A,δ*_ at *Z* = 1 and *κ* = *κ*_max_.

### D.6 Non-uniform placement of selected variance

Because the chromosomal architecture of fitness variance in wild populations is unknown [115], uniform placement is a baseline assumption. Let 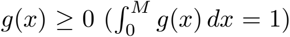 denote the density of selected variance in map distance. With neutral sites uniform in map distance, symmetry of *K*_H_(|*x − y*|) gives

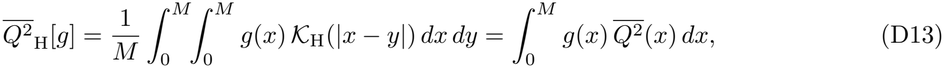

where the square brackets mark the chromosome average as a function of the whole density *g*. Because *K*_H_ depends only on the distance between the two sites, the profile of Equation (D2) has two equivalent meanings. Evaluated at a neutral site, it averages the kernel over selected loci spread uniformly; evaluated at a selected locus, as here, it averages the kernel over neutral sites spread uniformly, and so measures how much a locus at *x* reduces diversity along the whole chromosome. With *g* = 1*/M* the chromosome average equals the uniform one, 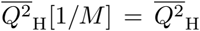, and for *J* loci of equal effect at *x*_1_, …, *x*_*J*_ it is the mean of 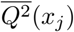. Because 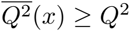 everywhere, Ω_*i*_ *≤* 1 holds for every *g*. The profile is strictly concave, because its second derivative, 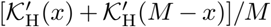, is negative. It is largest at the midpoint and smallest at the telomeres, because a central locus is linked to neutral sites on both chromosome arms and a terminal one to sites on one arm only.

Because linked diversity depends on the expectation of the convex function exp 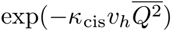, Jensen’s inequality implies that for a single locus placed uniformly at random on [0, *M* ],

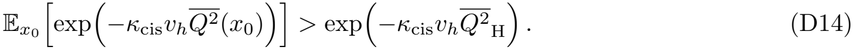

For *J* independently placed equal-effect loci, a second-order expansion gives

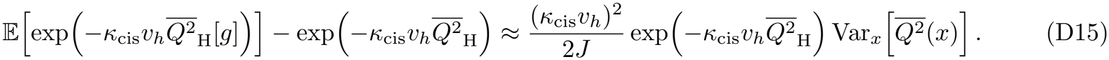

Under random placement independent of recombination rate, uniform density therefore maximizes the expected linked reduction in *N*_*e*_. This ordering can reverse, however, when selected alleles are concentrated in regions of suppressed recombination such as pericentromeres or chromosomal inversions. Even without physical clustering, a region of low recombination places long stretches of selected and neutral sequence within a short map interval, so linkage there is tighter than under uniform placement. Processes that concentrate selected alleles in low-recombination regions or inversions [116, 117] add to this effect.

## Supplementary Tables

**Table S2:**
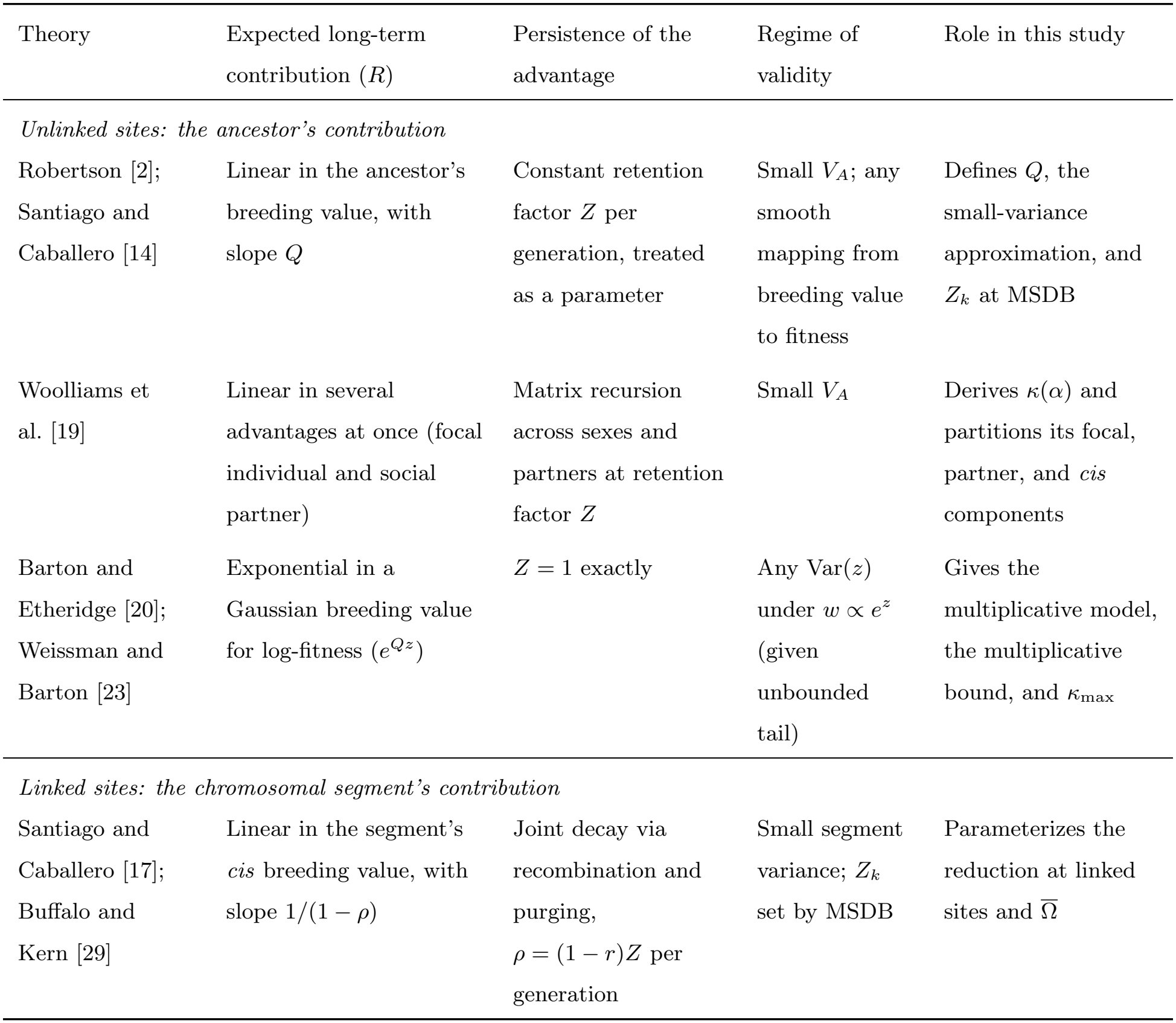
Theories of drift under selection compute the same quantity, an ancestor’s expected long-term genetic contribution *R*, from its heritable selective advantage. *R* is the ancestor’s share of the population’s genes many generations later, relative to an average ancestor’s. At linked sites, the long-term contribution that sets drift is that of the chromosomal segment carrying the focal neutral copy, not that of the individual ancestor.

| Theory | Expected long-term contribution (R) | Persistence of the advantage | Regime of validity | Role in this study |
|---|---|---|---|---|
| *Unlinked sites: the ancestor's contribution* |  |  |  |  |
| Robertson [2]; Santiago and Caballero [14] | Linear in the ancestor's breeding value, with slope Q | Constant retention factor Z per generation, treated as a parameter | Small V<sub>A</sub>; any smooth mapping from breeding value to fitness | Defines Q, the small-variance approximation, and Z<sub>k</sub> at MSDB |
| Woolliams et al. [19] | Linear in several advantages at once (focal individual and social partner) | Matrix recursion across sexes and partners at retention factor Z | Small V<sub>A</sub> | Derives κ(α) and partitions its focal, partner, and *cis* components |
| Barton and Etheridge [20]; Weissman and Barton [23] | Exponential in a Gaussian breeding value for log-fitness (e<sup>Qz</sup>) | Z = 1 exactly | Any Var(z) under w ∝ e<sup>z</sup> (given unbounded tail) | Gives the multiplicative model, the multiplicative bound, and κ<sub>max</sub> |
| *Linked sites: the chromosomal segment's contribution* |  |  |  |  |
| Santiago and Caballero [17]; Buffalo and Kern [29] | Linear in the segment's *cis* breeding value, with slope 1/(1 − ρ) | Joint decay via recombination and purging, ρ = (1 − r)Z per generation | Small segment variance; Z<sub>k</sub> set by MSDB | Parameterizes the reduction at linked sites and $\overline{\Omega}$ |

**Table S3:**
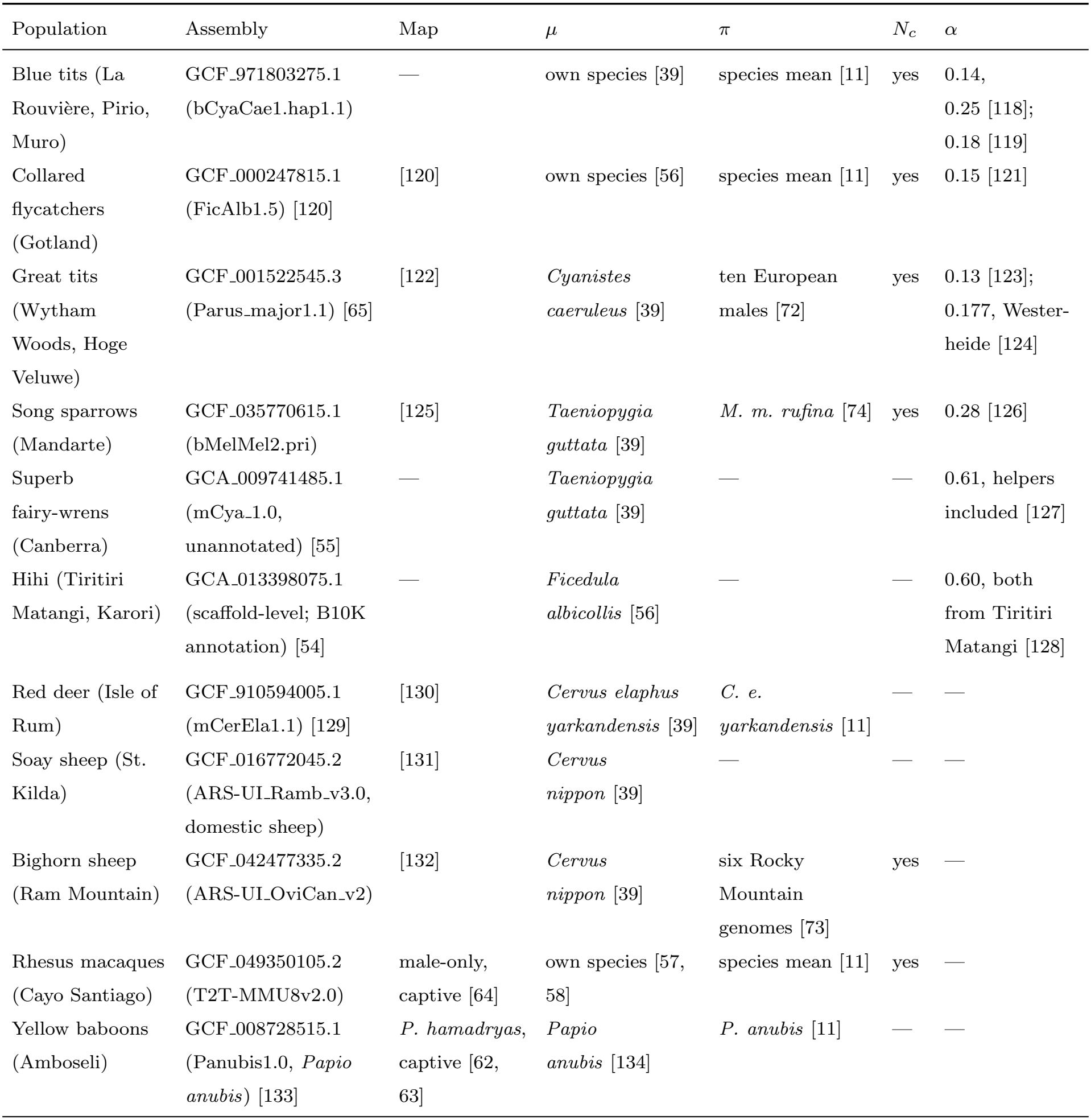
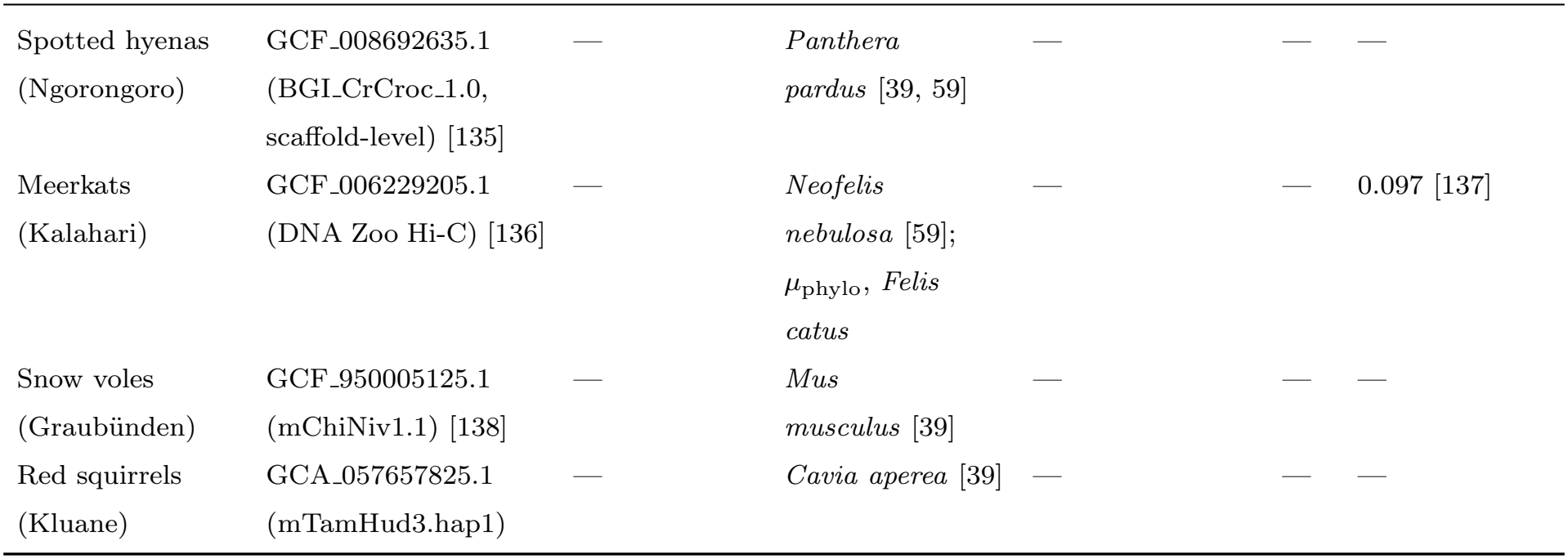
Sources of the inputs for each population. Assembly: NCBI accession (assembly name); all are of the study species except where a species is named. Map: published pedigree linkage map, where one was joined to the assembly. *μ*: species of the direct mutation-rate estimate, from the compilation of Lewin and Eyre-Walker [11], and of the phylogenetic prediction of Wang and Obbard [60], which is the same species except where noted. *π*: source of nucleotide diversity. *N*_*c*_: census size of the study species from Buffalo [10]. *α*: extra-pair paternity rate of each socially monogamous population, in the order of the sites named. A dash: none used, or random mating for *α*.

| Population | Assembly | Map | μ | π | *N*<sub>*c*</sub> | α |
|---|---|---|---|---|---|---|
| Blue tits (La Rouvière, Pirio, Muro) | GCF_971803275.1 (bCyaCae1.hap1.1) | — | own species [39] | species mean [11] | yes | 0.14, 0.25 [118]; 0.18 [119] |
| Collared flycatchers (Gotland) | GCF_000247815.1 (FicAlb1.5) [120] | [120] | own species [56] | species mean [11] | yes | 0.15 [121] |
| Great tits (Wytham Woods, Hoge Veluwe) | GCF_001522545.3 (Parus_major1.1) [65] | [122] | *Cyanistes caeruleus* [39] | ten European males [72] | yes | 0.13 [123]; 0.177, Wester-heide [124] |
| Song sparrows (Mandarte) | GCF_035770615.1 (bMelMel2.pri) | [125] | *Taeniopygia guttata* [39] | *M. m. rufina* [74] | yes | 0.28 [126] |
| Superb fairy-wrens (Canberra) | GCA_009741485.1 (mCya_1.0, unannotated) [55] | — | *Taeniopygia guttata* [39] | — | — | 0.61, helpers included [127] |
| Hihi (Tiritiri Matangi, Karori) | GCA_013398075.1 (scaffold-level; B10K annotation) [54] | — | *Ficedula albicollis* [56] | — | — | 0.60, both from Tiritiri Matangi [128] |
| Red deer (Isle of Rum) | GCF_910594005.1 (mCerEla1.1) [129] | [130] | *Cervus elaphus yarkandensis* [39] | *C. e. yarkandensis* [11] | — | — |
| Soay sheep (St. Kilda) | GCF_016772045.2 (ARS-UI_Ramb_v3.0, domestic sheep) | [131] | *Cervus nippon* [39] | — | — | — |
| Bighorn sheep (Ram Mountain) | GCF_042477335.2 (ARS-UI_OviCan_v2) | [132] | *Cervus nippon* [39] | six Rocky Mountain genomes [73] | yes | — |
| Rhesus macaques (Cayo Santiago) | GCF_049350105.2 (T2T-MMU8v2.0) | male-only, captive [64] | own species [57, 58] | species mean [11] | yes | — |
| Yellow baboons (Amboseli) | GCF_008728515.1 (Panubis1.0, *Papio anubis*) [133] | *P. hamadryas*, captive [62, 63] | *Papio anubis* [134] | *P. anubis* [11] | — | — |
| Spotted hyenas (Ngorongoro) | GCF_008692635.1 (BGI_CrCroc_1.0, scaffold-level) [135] | — | *Panthera pardus* [39, 59] | — | — | — |
| Meerkats (Kalahari) | GCF_006229205.1 (DNA Zoo Hi-C) [136] | — | *Neofelis nebulosa* [59]; μ<sub>phylo</sub>, *Felis catus* | — | — | 0.097 [137] |
| Snow voles (Graubünden) | GCF_950005125.1 (mChiNiv1.1) [138] | — | *Mus musculus* [39] | — | — | — |
| Red squirrels (Kluane) | GCA_057657825.1 (mTamHud3.hap1) | — | *Cavia aperea* [39] | — | — | — |

**Table S4:** Asymptotic limits of the chromosome-wide average 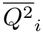 for a single class of deleterious mutations purged at rate *s* (*Z* = 1*−s*) or slowly (*s→*0). Complete linkage requires *M*_*i*_ *≪* 1*−Z* and free recombination requires *M*_*i*_ *≫* 1*/*(1 *− Z*); empirical chromosomes lie in the intermediate regime *≈* 2*/*[*M*_*i*_(1 *− Z*)].

| Purging regime | Complete linkage | Free recombination |
|---|---|---|
| At rate *s* (*Z* = 1 − *s*) | *s*<sup>−2</sup> | 4/(1 + *s*)<sup>2</sup> |
| Slow (*s* → 0, *Z* → 1) | ∞ | 4 |

## Supplementary Methods: approximations biased toward passing

On the diversity side, the approximations maximize the reduction in effective size that a given *V*_*A*_ causes, and so minimize *V*_*A,δ*_ (subsection 2.7):

- **Unbounded multiplicative fitness:** The multiplicative bound places no ceiling on family size (sub-section 2.4).
- **Maximum mating-system amplification:** The bound uses *κ*_max_ = 2 under social monogamy, not the smaller *κ*(*α*) (subsubsection 2.6.1).
- **Complete retention at unlinked sites:** The reduction at unlinked sites uses *Z* = 1, so *Q* = 2 rather than 2*/*(1 + *s*_*k*_) (subsection 2.3).
- **Pre-breeding survival treated as fecundity:** *Q* = 2 applies to all of 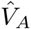, although survival to breeding age has *Q* = 1 (subsection 3.1).
- **Absence of crossover interference:** Haldane’s map omits interference, which raises recombination at intermediate distances and weakens linked selection.
- **Empirical scaling of linked variance:** The linked term carries all of 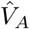, not the smaller 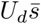 that MSDB maintains (subsubsection 2.6.2). On the maintenance side, the approximations maximize the variance that recurrent mutation sustains, and so maximize *c*_MSD_ and minimize *s*^*\**^:
- **Strong-selection limit:** 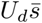 is the most variance MSDB maintains at any effective size (subsection 2.2).
- **Deleterious target:** Every mutation in the primary target, twice the autosomal exome, is treated as deleterious, synonymous sites included (subsection 3.3).

## Supplementary Results: linkage

In our predictions, the contrast between birds and mammals does not follow the recombination maps. At the same selection coefficient and *V*_*A*_, the maps alone give similar linked factors 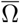 (Equation (20)) in the two taxa (0.55–0.73; Figure S4b), although the bird genomes recombine about twice as much per megabase. The contrast follows instead the DFEs assigned to the two taxa, whose mean heterozygous effects are 0.0013– 0.0021 in birds, an order of magnitude lower than the 0.014–0.020 assigned to the mapped mammals. Because each DFE was assigned by taxon from related species (Methods), the contrast is attributable to the assigned DFEs within the model and is not evidence that the DFEs of these populations differ. The assigned DFEs differ in how fast they purge. Slow purging, in turn, gives a larger reduction at linked sites from the same *V*_*A*_ (subsubsection 2.6.2).

The reduction at linked sites is also steeply nonlinear in the amount of variance. The two great tit populations share maps and a DFE, yet 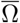 is 0.031 at Wytham Woods, where 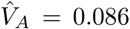, and 0.49 at Hoge Veluwe, where 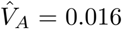 (Figure S4a). Because the variance on the haplotype multiplies the excess persistence of its associations in the exponent of Equation (20), the linked factor of each chromosome decreases geometrically as 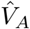 increases. Partitioning the variance among chromosomes by gene count rather than by coding length does not change these results (Figure S5). Partitioning it by the annotated functional target with up to 15% of the unannotated sequence changes the linked *N*_*e*_*/N* by at most 13% in eight of the nine mapped populations and by 40% in the great tits at Wytham Woods, and each of the six with an observed deficit remains at least 50-fold below it (Figure S7).

## Supplementary Results: maintenance

The mammals other than the spotted hyenas fail the maintenance test by 9- to 22-fold at the primary target (the meerkats by 2.0-fold) and by 2- to 5-fold when 8% of the unannotated sequence is counted as target, at which the meerkats pass (Figure S9).

## Supplementary Figures

**Figure S1:**
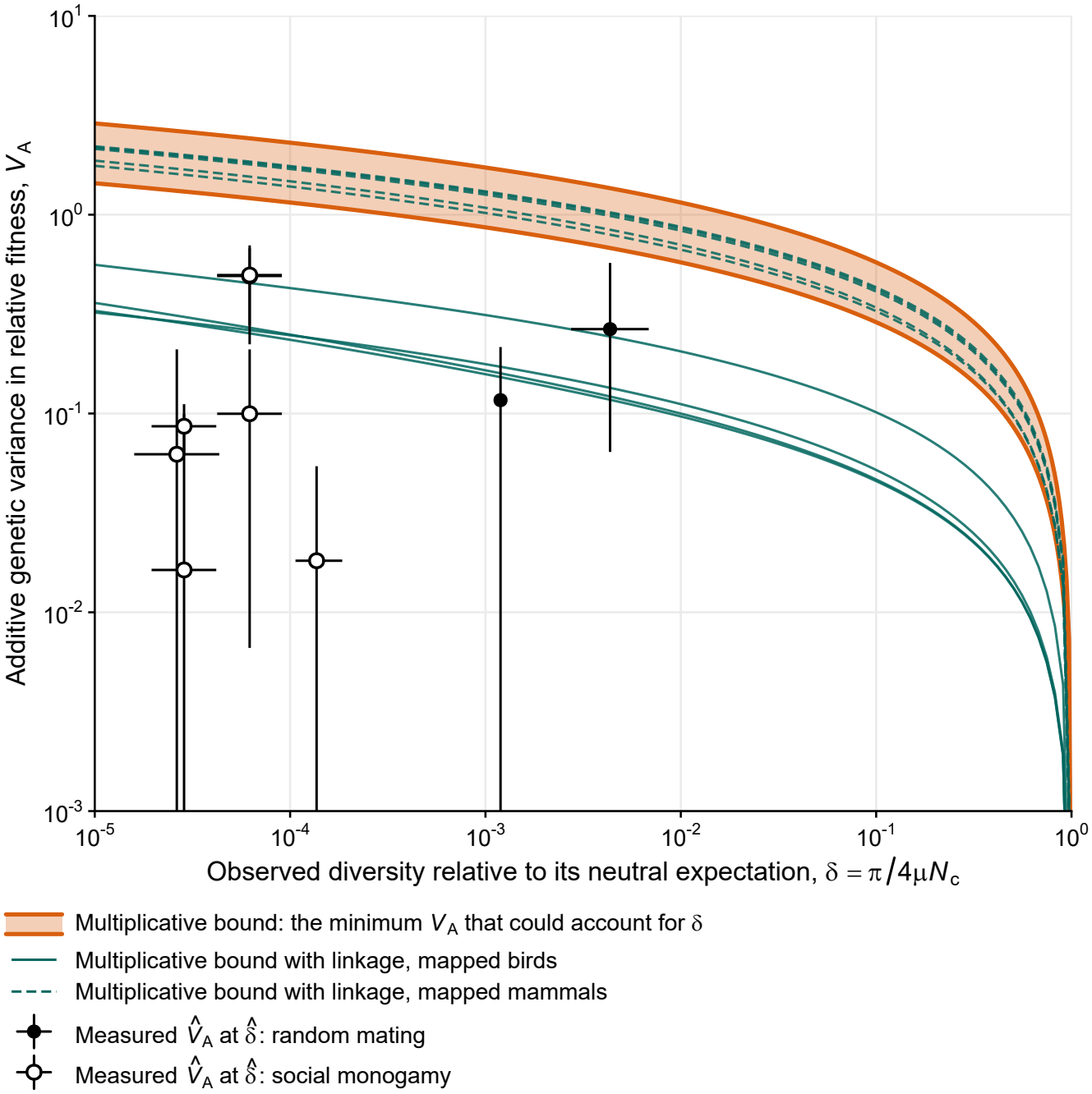
The minimum variance that could account for a diversity deficit increases only with the logarithm of the deficit. Orange band: the minimum *V*_*A*_ that could account for a diversity ratio *δ*, the reciprocal of the deficit, under the multiplicative bound, between *κ*_max_ = 1 (upper line, random mating) and 2 (lower line, strict monogamy). Teal lines: the same with linkage for each of the nine mapped populations at its assigned empirical DFE (solid, birds; dashed, mammals). Points: the nine populations with *π* and a census size, at 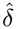 and the posterior mode of 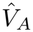, with bars over the 95% HPD interval of 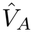 (vertical) and the 95% CI of the mutation rate (horizontal); filled, random mating; open, social monogamy. A population would pass the diversity test where its estimate reaches the line for its mating system, or with linkage its own teal line.

**Figure S2:**
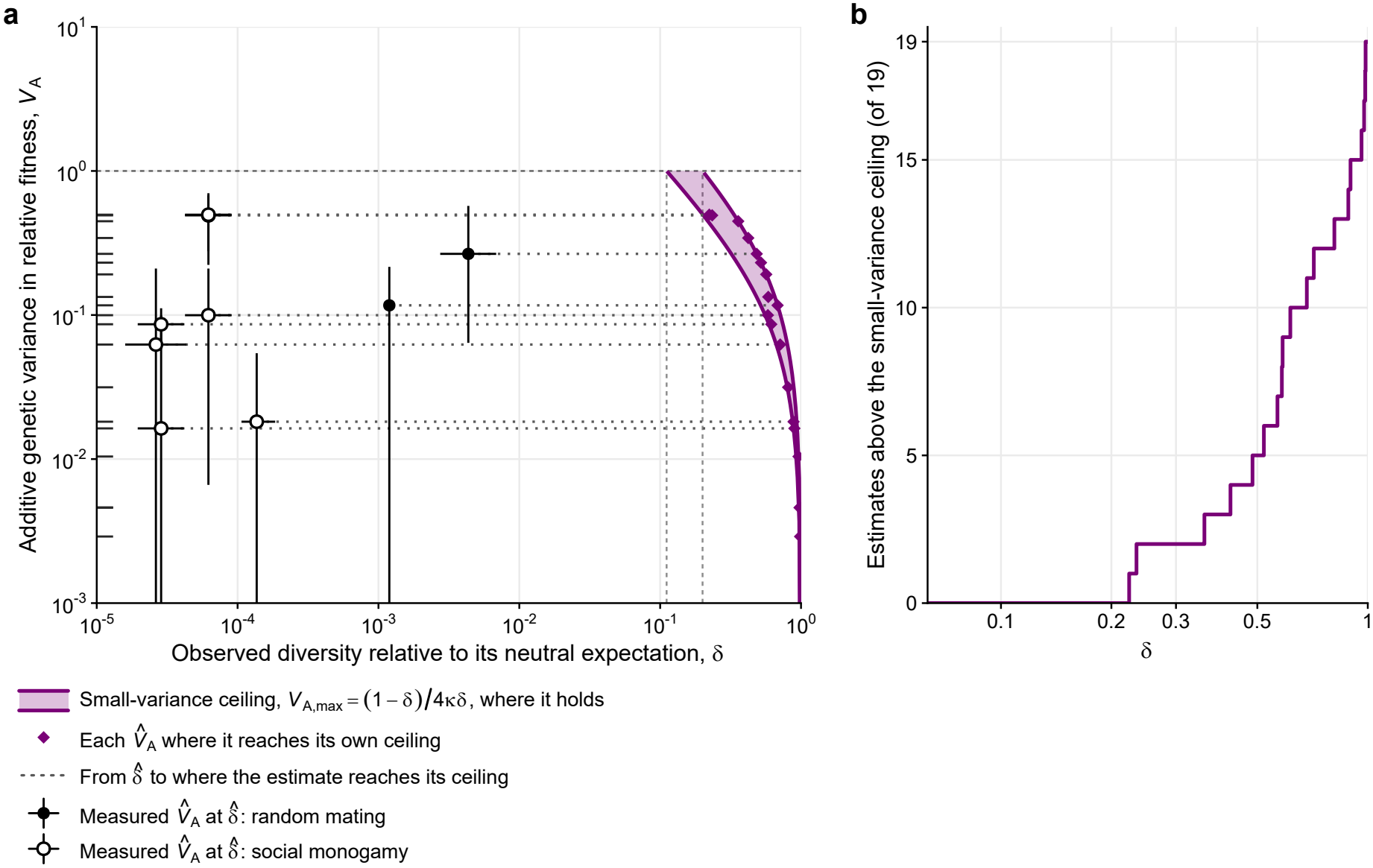
No pedigree estimate exceeds the small-variance ceiling that its population’s diversity implies. (a) Purple band: the small-variance ceiling, *V*_*A*,max_, for *κ* = 1 (upper line) to 2 (lower line), drawn only where it is at most 1. The horizontal dashed line marks *V*_*A*_ = 1, the limit of the small-variance approximation; the vertical dashed lines mark where the ceiling reaches 1, at *δ* = 0.11 (*κ* = 2) and 0.2 (*κ* = 1). Purple diamonds: each of the 19 posterior modes of 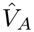 at the *δ* where it reaches the ceiling for its population’s *κ*. Points: the nine populations with *π* and a census size, at their 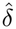 and the posterior mode of 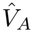, with bars over the 95% HPD interval of 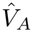 (vertical) and the 95% CI of the mutation rate (horizontal); filled, random mating; open, social monogamy. A dotted line joins each point to its diamond. Ticks on the y-axis: the posterior modes of all 19 populations. (b) The number of the 19 posterior modes above the ceiling, each at its population’s *κ*, as a function of *δ*.

**Figure S3:**
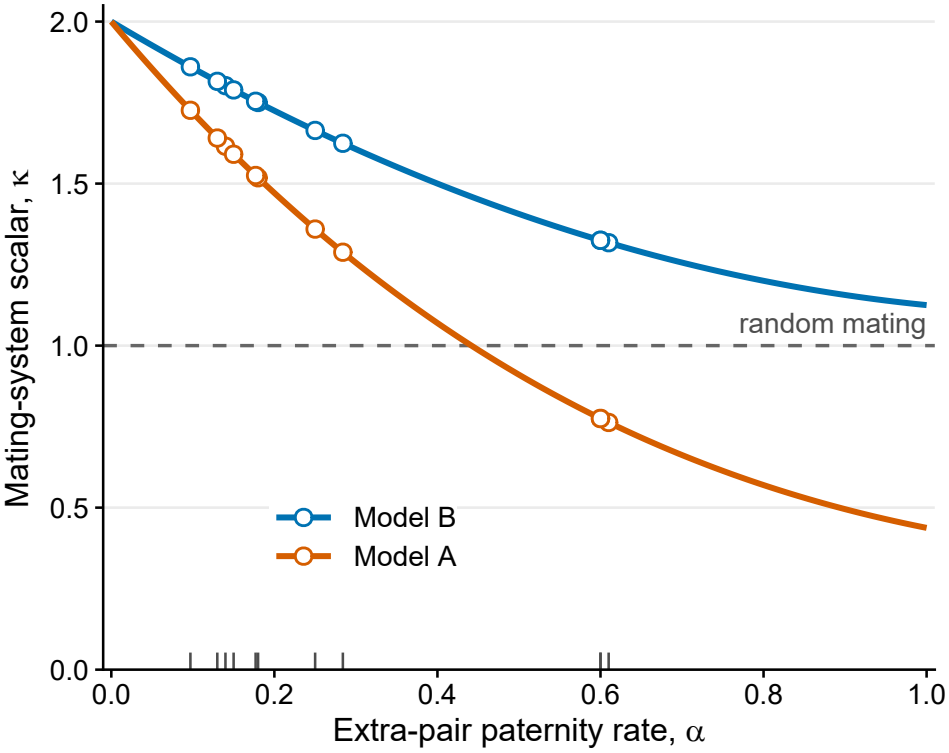
Extra-pair paternity lowers the mating-system scalar, and under Model B it stays above the random-mating value. The scalar *κ* against the extra-pair paternity rate *α*, with Model B (extra-pair success proportional to a male’s breeding value; primary) in blue and Model A (extra-pair siring independent of it; sensitivity) in orange. Both models give *κ* = 2 under strict monogamy. Under Model A, *κ* drops below the random-mating value (dashed line) once *α* exceeds 0.44. Points and ticks mark the eleven socially monogamous populations at their extra-pair paternity rates.

**Figure S4:**
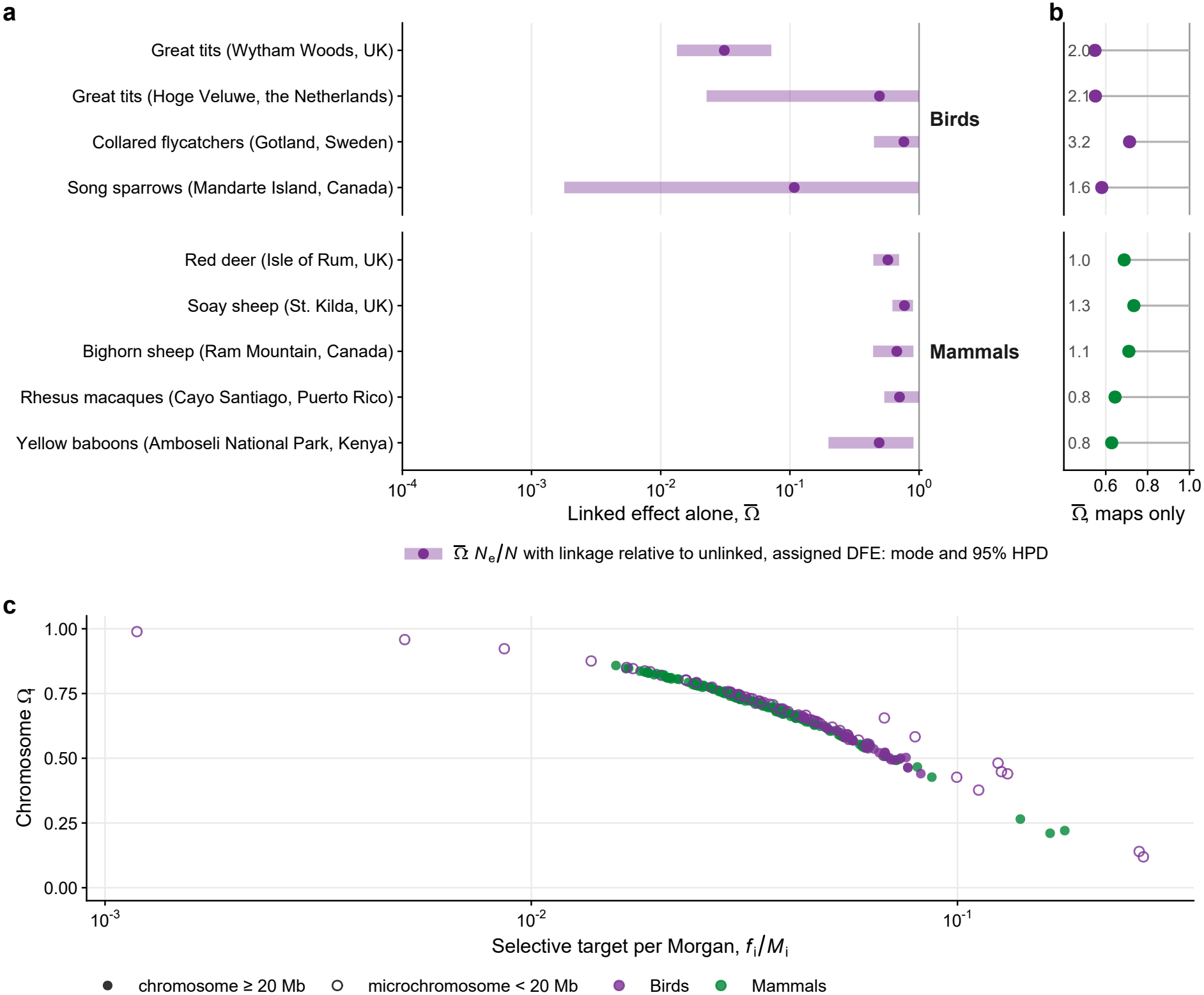
The linked factor is steeply nonlinear in the measured variance, and from the maps alone it is similar in birds and mammals. (a) The linked factor alone, 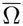, is the share of diversity kept once linkage is added (genome-wide *N*_*e*_*/N* with linkage relative to the unlinked value at the same retention factor), for the nine populations with linkage maps, at the empirical DFE assigned to each population. Dot: at the posterior mode of 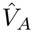; bar: over its 95% HPD. (b) The linked factor 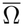 from the maps alone, at a uniform selection coefficient *s* = 0.01 and *V*_*A*_ = 0.1, with each genome’s map length per megabase (cM/Mb) on the left. (c) The linked factor of each chromosome, Ω_*i*_, against its selected target per Morgan, *f*_*i*_*/M*_*i*_, which sets how much of the chromosome’s variance lies close enough to a neutral site to stay associated with it for longer than at an unlinked site, at the same values as (b). Open circles: microchromosomes (*<* 20 Mb).

**Figure S5:**
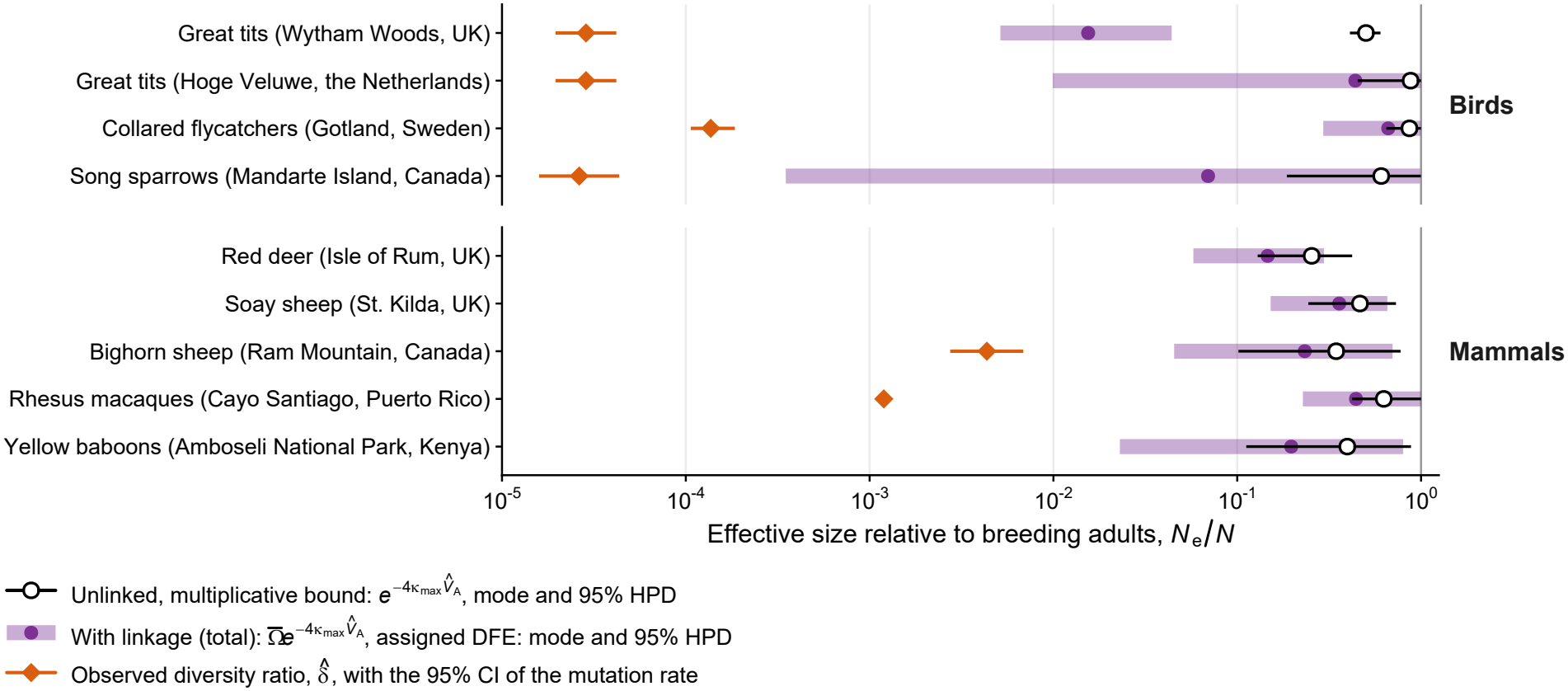
Partitioning the variance by gene count leaves the linked results unchanged. As Figure 3, with the additive variance partitioned among chromosomes by annotated protein-coding gene count rather than by coding length.

**Figure S6:**
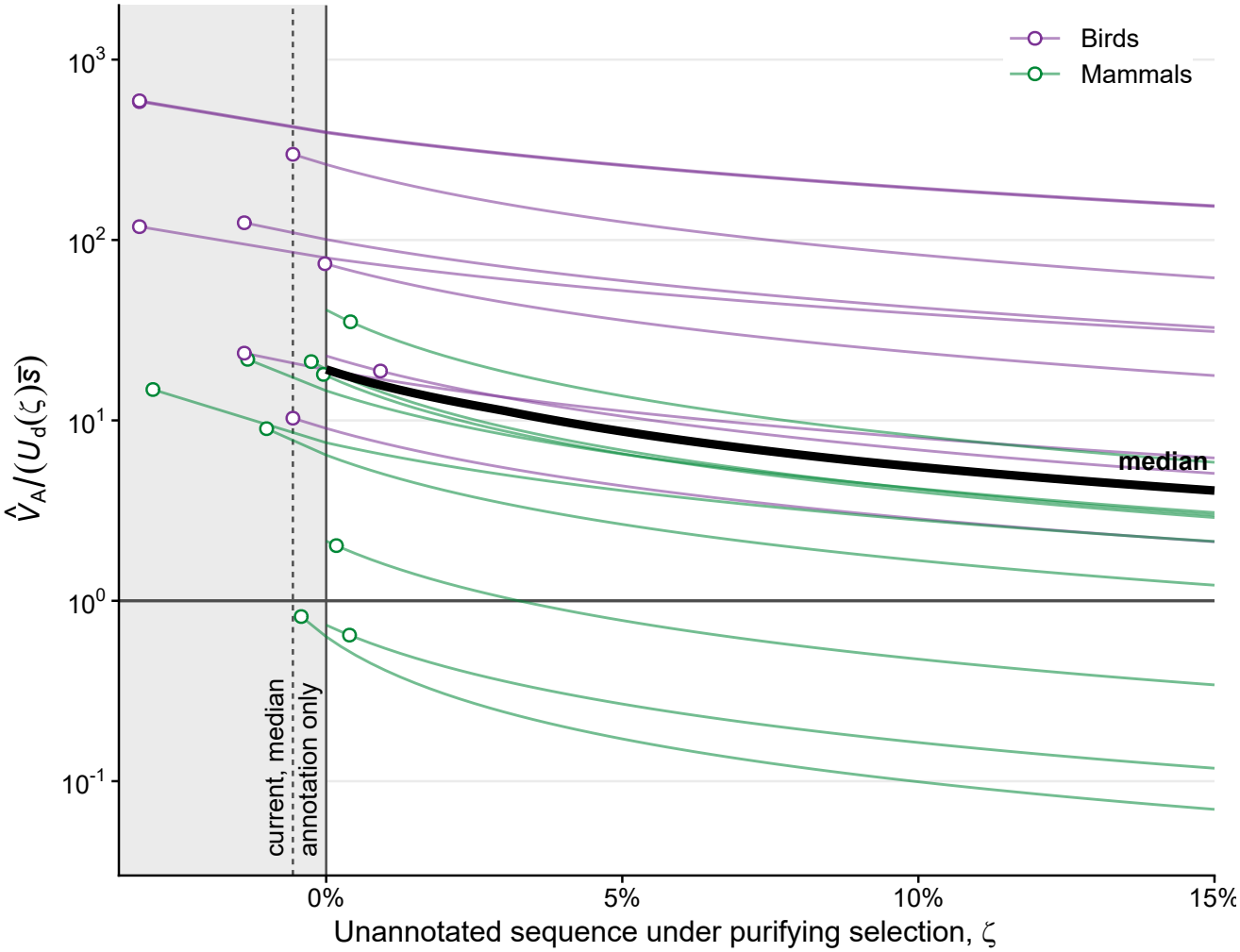
A larger functional target eases but does not remove the maintenance excess. The deleterious target is the annotated functional sequence (coding sequence, untranslated regions and non-coding RNA exons) plus a fraction *ζ* of the remaining, unannotated sequence. The measured variance relative to the variance MSDB maintains at the assigned empirical DFE, 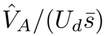, at the posterior mode of 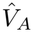 and the direct mutation rate, for the 18 populations with an annotated assembly. Faint lines: populations, colored by taxon; bold line: median; open circles: the value of *ζ* at which each population’s target equals the doubled autosomal exome used in the main text, which for most populations is smaller than the annotation alone (gray). Solid vertical line: annotation only; dashed vertical line: the median of the open circles; horizontal line: the variance MSDB maintains.

**Figure S7:**
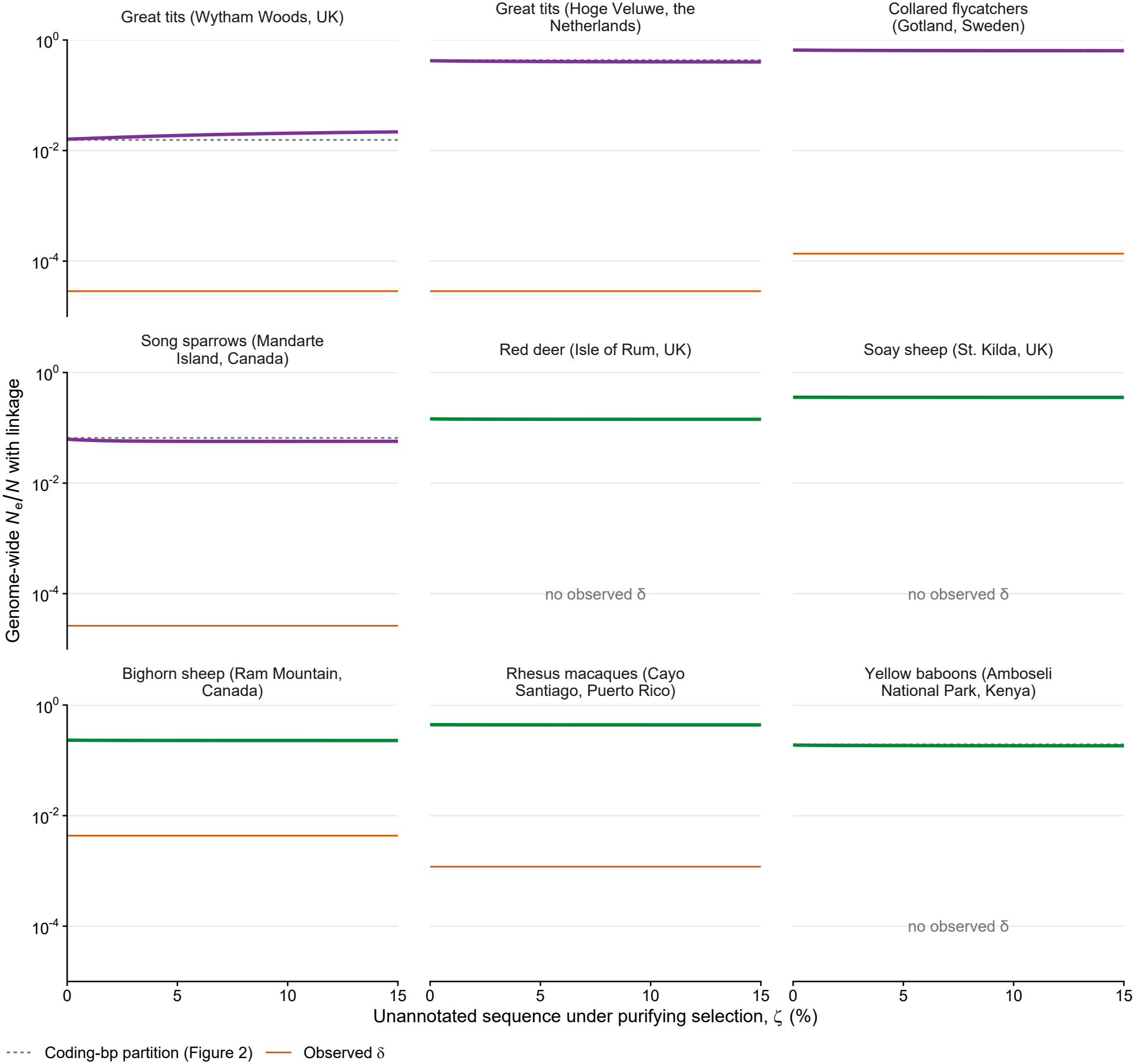
A larger functional target barely moves the linked prediction. Genome-wide *N*_*e*_*/N* with linkage for the nine mapped populations, with the variance partitioned among chromosomes by the target of Figure S6 (annotated functional sequence plus a fraction *ζ* of the unannotated sequence), at the posterior mode of 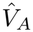 and the assigned DFE. Dashed gray: the coding-length partition of Figure 3; orange: observed diversity ratio, where available.

**Figure S8:**
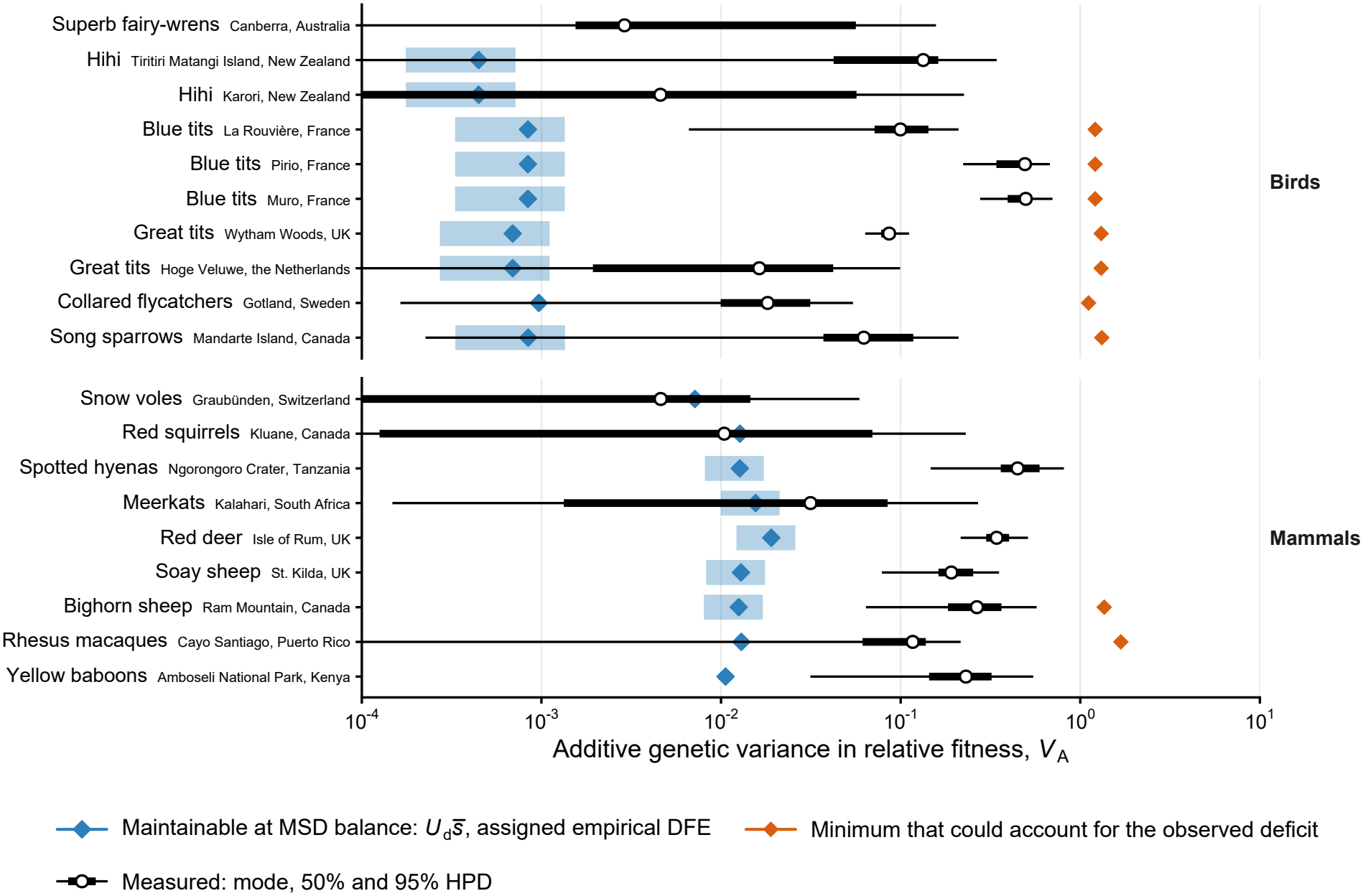
The measured variances lie mostly above what MSDB maintains and below what the observed deficits require. One row per population, on a shared *V*_*A*_ axis. Blue diamonds: the variance MSDB maintains in the strong-selection limit, 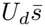, with *U*_*d*_ from the direct mutation rate and 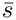 the mean of the empirical DFE assigned to the population; where the assignment is an equal mixture of two species, the band spans *U*_*d*_ times their means. Circles and bars: measured 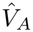, posterior mode, 50% (thick) and 95% (thin) HPD. Orange diamonds: the minimum variance that could account for the observed deficit at unlinked sites, *V*_*A,δ*_, with bars over the 95% CI of the mutation rate. The superb fairy-wren has no annotated assembly. Orange diamonds are drawn for the nine populations with *π* and a census size; red deer and yellow baboons have no census size for their own species, Soay sheep have no diversity estimate, and the other seven populations have neither.

**Figure S9:**
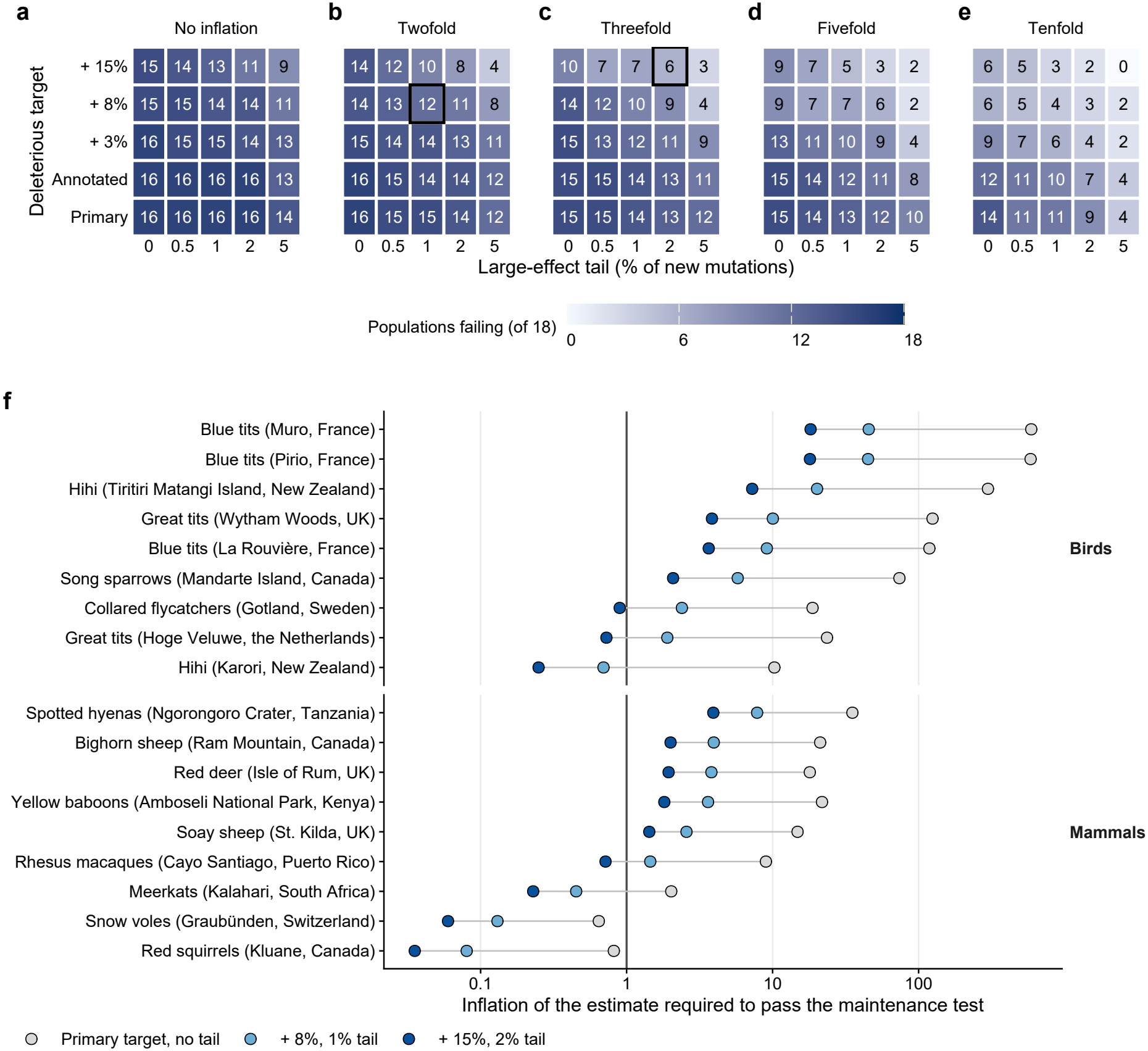
Combined departures in the target, the tail of large effects and the estimate bring the median population close to passing the maintenance test, but most populations pass only when all three are large. (a–e) Number of the 18 populations with an annotated assembly in which the measured variance, divided by the inflation factor, still exceeds the variance MSDB maintains, at the posterior mode of 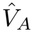 and the point value of the direct mutation rate. Rows: deleterious target, either the primary doubled autosomal exome or the annotated functional sequence plus a fraction *ζ* of the unannotated sequence (Annotated, *ζ* = 0; +3%, +8% and +15%). Columns: percentage of new mutations given a heterozygous effect of 0.5, the rest following the assigned DFE. Panels: factor by which the pedigree estimate is taken to overstate the long-term variance. Outlined cells: the moderate (b) and stronger (c) joint scenarios reported in Results. (f) Inflation of the estimate that each population would require to pass, at three combinations of target and tail; vertical line: no inflation.

**Figure S10:**
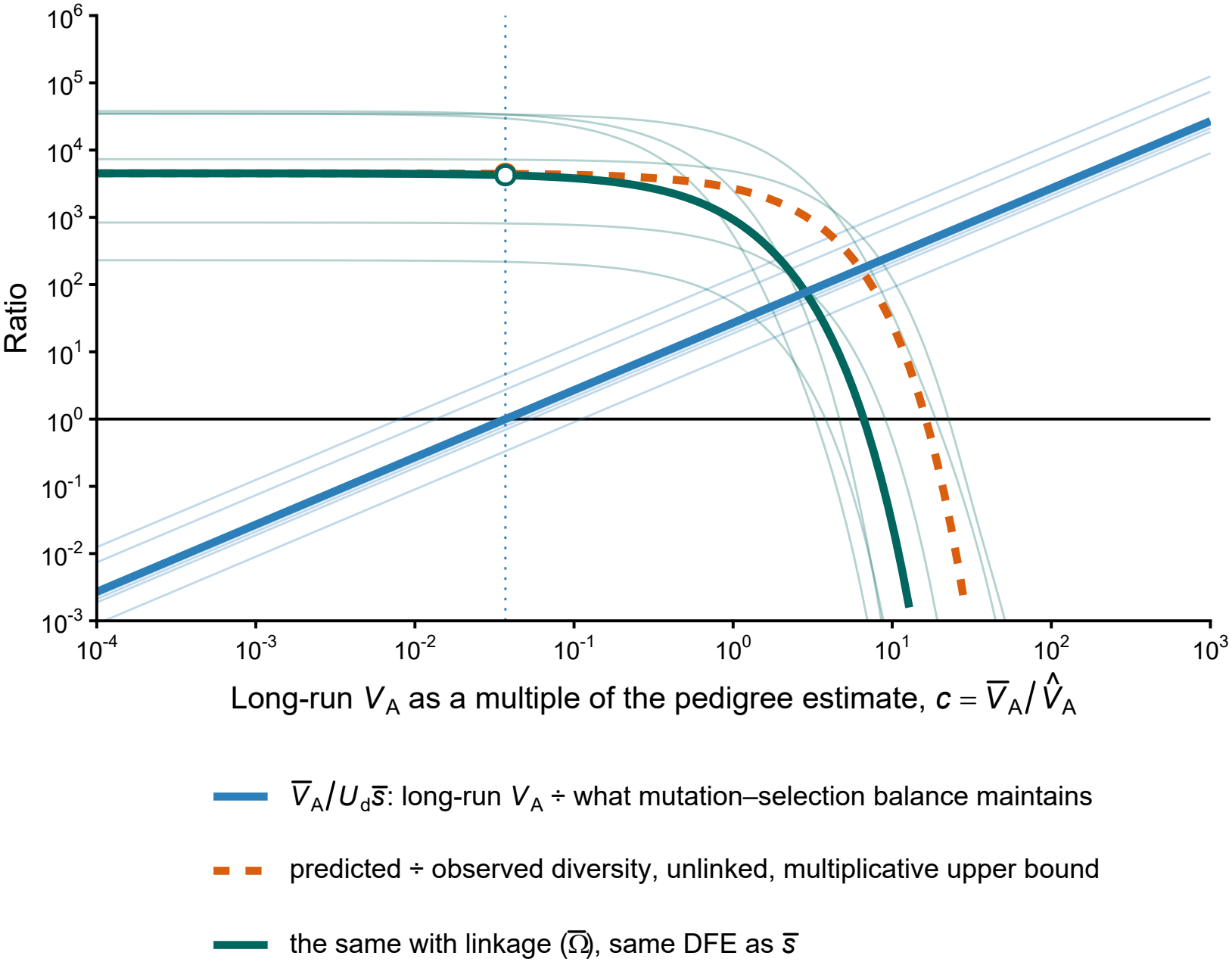
Linkage narrows the gap between the two tests but does not resolve it. As Figure 5, with linkage, for the six populations with both an observed deficit and a linkage map (collared flycatchers, rhesus macaques, song sparrows, bighorn sheep and both great tit populations). Blue: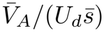. Orange dashed: predicted relative to observed diversity, unlinked, under the multiplicative bound. Green: the same with linkage, multiplied by 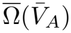 at the same assigned DFE, so that both tests share a single distribution of effects. Open circle: the average at 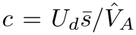, where the variance equals what that DFE maintains, the self-consistent prediction of MSDB with background selection. Bold: species averages of the log_10_ ratio (great tits averaged first); faint: populations.

**Figure S11:**
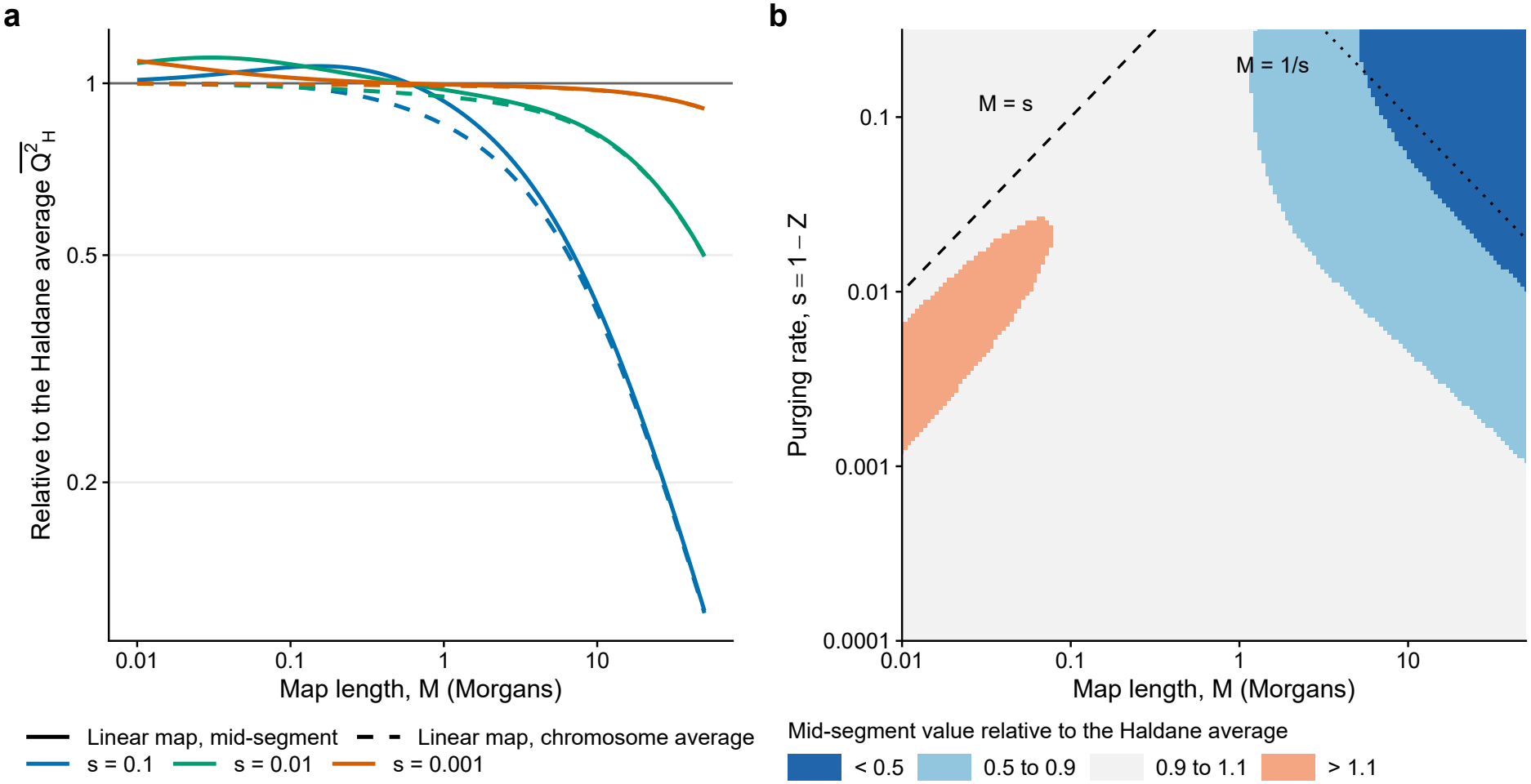
The linear map and the mid-segment value depart from the Haldane average only on long maps with fast purging, and the mid-segment value briefly overshoots on short ones. Each value is relative to the Haldane chromosome average 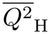 used in the analysis, for a single class of deleterious mutations purged at rate *s*. (a) Against map length, at three purging rates. Solid: the linear-map value at the middle of the segment, as used by Buffalo and Kern; dashed: the linear-map chromosome average. (b) The mid-segment value over map length and purging rate. Dashed line: *M* = *s*, below which the chromosome approaches complete linkage; dotted line: *M* = 1*/s*, above which it approaches free recombination.

## References

[1] James F. Crow. “Some possibilities for measuring selection intensities in man”. In: Human Biology 30.1 (1958), pp. 1–13. url: https://www.jstor.org/stable/41449168.

[2] Alan Robertson. “Inbreeding in artificial selection programmes”. In: Genetical Research 2.2 (1961), pp. 189–194. doi: 10.1017/S0016672300000690.

[3] Loeske E. B. Kruuk. “Estimating genetic parameters in natural populations using the ‘animal model’”. In: Philosophical Transactions of the Royal Society of London. Series B: Biological Sciences 359.1446 (2004), pp. 873–890. doi: 10.1098/rstb.2003.1437.

[4] Erik Postma. “Four decades of estimating heritabilities in wild vertebrate populations: improved methods, more data, better estimates?” In: Quantitative Genetics in the Wild. Ed. by Anne Charmantier, Dany Garant, and Loeske E. B. Kruuk. Oxford: Oxford University Press, 2014, pp. 16–33. doi: 10.1093/acprof:oso/9780199674237.003.0002.

[5] Alan Robertson. “A mathematical model of the culling process in dairy cattle”. In: Animal Production 8.1 (1966), pp. 95–108. doi: 10.1017/S0003356100037752.

[6] George R. Price. “Selection and covariance”. In: Nature 227.5257 (1970), pp. 520–521. doi: 10.1038/227520a0.

[7] Juha Merilä, Ben C. Sheldon, and Loeske E. B. Kruuk. “Explaining stasis: microevolutionary studies in natural populations”. In: Genetica 112–113 (2001), pp. 199–222. doi: 10.1023/A:1013391806317.

[8] Walid Mawass and Emmanuel Milot. “Contrasting evolutionary outcomes in a human life history trait which is heritable and under consistent unbiased directional selection”. In: bioRxiv (2025). doi: 10.1101/2025.10.13.682154. url: https://www.biorxiv.org/content/10.1101/2025.10.13.682154v1.

[9] Timothée Bonnet, Michael B. Morrissey, Pierre de Villemereuil, Susan C. Alberts, Peter Arcese, Liam D. Bailey, Stan Boutin, Patricia Brekke, Lauren J. N. Brent, Glauco Camenisch, et al. “Genetic variance in fitness indicates rapid contemporary adaptive evolution in wild animals”. In: Science 376.6596 (2022), pp. 1012–1016. doi: 10.1126/science.abk0853.

[10] Vince Buffalo. “Quantifying the relationship between genetic diversity and population size suggests natural selection cannot explain Lewontin’s paradox”. In: eLife 10 (2021), e67509. doi: 10.7554/eLife.67509.

[11] Loveday Lewin and Adam Eyre-Walker. “A comparative analysis of long-term effective population sizes across eukaryotes”. In: Molecular Ecology 35.4 (2026), e70265. doi: 10.1111/mec.70265.

[12] Richard C. Lewontin. The Genetic Basis of Evolutionary Change. New York: Columbia University Press, 1974.

[13] Ellen M. Leffler, Kevin Bullaughey, Daniel R. Matute, Wynn K. Meyer, Laure Ségurel, Aarti Venkat, Peter Andolfatto, and Molly Przeworski. “Revisiting an old riddle: what determines genetic diversity levels within species?” In: PLoS Biology 10.9 (2012), e1001388. doi: 10.1371/journal.pbio.1001388.

[14] Enrique Santiago and Armando Caballero. “Effective size of populations under selection”. In: Genetics 139.2 (1995), pp. 1013–1030. doi: 10.1093/genetics/139.2.1013.

[15] Sewall Wright. “Size of population and breeding structure in relation to evolution”. In: Science 87.2263 (1938), pp. 430–431. doi: 10.1126/science.87.2263.425.b.

[16] Naomi R. Wray and Robin Thompson. “Prediction of rates of inbreeding in selected populations”. In: Genetical Research 55.1 (1990), pp. 41–54. doi: 10.1017/S0016672300025180.

[17] Enrique Santiago and Armando Caballero. “Effective size and polymorphism of linked neutral loci in populations under directional selection”. In: Genetics 149.4 (1998), pp. 2105–2117. doi: 10.1093/genetics/149.4.2105.

[18] John A. Woolliams, Naomi R. Wray, and Robin Thompson. “Prediction of long-term contributions and inbreeding in populations undergoing mass selection”. In: Genetical Research 62.3 (1993), pp. 231–242. doi: 10.1017/S0016672300031943.

[19] John A. Woolliams, Piter Bijma, and Beatriz Villanueva. “Expected genetic contributions and their impact on gene flow and genetic gain”. In: Genetics 153.2 (1999), pp. 1009–1020. doi: 10.1093/genetics/153.2.1009.

[20] Nicholas H. Barton and Alison M. Etheridge. “The relation between reproductive value and genetic contribution”. In: Genetics 188.4 (2011), pp. 953–973. doi: 10.1534/genetics.111.127555.

[21] Nicholas H. Barton. “Genetic hitchhiking”. In: Philosophical Transactions of the Royal Society of London. Series B: Biological Sciences 355.1403 (2000), pp. 1553–1562. doi: 10.1098/rstb.2000.0716.

[22] Brian Charlesworth. “The effects of deleterious mutations on evolution at linked sites”. In: Genetics 190.1 (2012), pp. 5–22. doi: 10.1534/genetics.111.134288.

[23] Daniel B. Weissman and Nicholas H. Barton. “Limits to the rate of adaptive substitution in sexual populations”. In: PLoS Genetics 8.6 (2012), e1002740. doi: 10.1371/journal.pgen.1002740.

[24] Richard R. Hudson and Norman L. Kaplan. “Deleterious background selection with recombination”. In: Genetics 141.4 (1995), pp. 1605–1617. doi: 10.1093/genetics/141.4.1605.

[25] Magnus Nordborg, Brian Charlesworth, and Deborah Charlesworth. “The effect of recombination on background selection”. In: Genetical Research 67.2 (1996), pp. 159–174. doi: 10.1017/S0016672300033619.

[26] Vince Buffalo and Graham Coop. “The linked selection signature of rapid adaptation in temporal genomic data”. In: Genetics 213.3 (2019), pp. 1007–1045. doi: 10.1534/genetics.119.302581.

[27] Vince Buffalo and Graham Coop. “Estimating the genome-wide contribution of selection to temporal allele frequency change”. In: Proceedings of the National Academy of Sciences 117.34 (2020), pp. 20672–20680. doi: 10.1073/pnas.1919039117.

[28] David A. Murphy, Eyal Elyashiv, Guy Amster, and Guy Sella. “Broad-scale variation in human genetic diversity levels is predicted by purifying selection on coding and non-coding elements”. In: eLife 12 (2023), e76065. doi: 10.7554/eLife.76065.

[29] Vince Buffalo and Andrew D. Kern. “A quantitative genetic model of background selection in humans”. In: PLOS Genetics 20.3 (2024), e1011144. doi: 10.1371/journal.pgen.1011144.

[30] J. B. S. Haldane. “The effect of variation of fitness”. In: The American Naturalist 71.735 (1937), pp. 337–349. doi: 10.1086/280722.

[31] James F. Crow. “Genetic loads and the cost of natural selection”. In: Mathematical Topics in Population Genetics. Ed. by Ken-ichi Kojima. Berlin, Heidelberg: Springer, 1970, pp. 128–177. doi: 10.1007/978-3-642-46244-3_5.

[32] Terumi Mukai, Sadao I. Chigusa, L. E. Mettler, and James F. Crow. “Mutation rate and dominance of genes affecting viability in Drosophila melanogaster “. In: Genetics 72.2 (1972), pp. 335–355. doi: 10.1093/genetics/72.2.335.

[33] Brian Charlesworth. “Causes of natural variation in fitness: evidence from studies of Drosophila populations”. In: Proceedings of the National Academy of Sciences 112.6 (2015), pp. 1662–1669. doi: 10.1073/pnas.1423275112.

[34] Toby Johnson and Nick Barton. “Theoretical models of selection and mutation on quantitative traits”. In: Philosophical Transactions of the Royal Society B: Biological Sciences 360.1459 (2005), pp. 1411–1425. doi: 10.1098/rstb.2005.1667.

[35] Brian Charlesworth and Jeffrey D. Jensen. “How can we resolve Lewontin’s paradox?” In: Genome Biology and Evolution 14.7 (2022), evac096. doi: 10.1093/gbe/evac096.

[36] Tim Connallon and Peter Czuppon. “Additive genetic variance for fitness maintained by balancing selection”. In: Evolution 80.5 (2026), pp. 926–935. doi: 10.1093/evolut/qpag006.

[37] Adam Eyre-Walker and Peter D. Keightley. “The distribution of fitness effects of new mutations”. In: Nature Reviews Genetics 8.8 (2007), pp. 610–618. doi: 10.1038/nrg2146.

[38] Christian D. Huber, Bernard Y. Kim, Clare D. Marsden, and Kirk E. Lohmueller. “Determining the factors driving selective effects of new nonsynonymous mutations”. In: Proceedings of the National Academy of Sciences 114.17 (2017), pp. 4465–4470. doi: 10.1073/pnas.1619508114.

[39] Lucie A. Bergeron, Søren Besenbacher, Jiao Zheng, Panyi Li, Mads Frost Bertelsen, Benoit Quintard, Joseph I. Hoffman, Zhipeng Li, Judy St. Leger, Changwei Shao, et al. “Evolution of the germline mutation rate across vertebrates”. In: Nature 615.7951 (2023), pp. 285–291. doi: 10.1038/s41586-023-05752-y.

[40] Meixi Lin, Sneha Chakraborty, Carlos Eduardo G. Amorim, Sergio F. Nigenda-Morales, Annabel C. Beichman, Paulina G. Nuñez-Valencia, Jonathan C. Mah, Jacqueline A. Robinson, Christopher C. Kyriazis, Christian D. Huber, et al. “The distribution of fitness effects of nonsynonymous mutations varies phylogenetically across animals”. In: PLOS Biology 24.9 (2026), e3003976. doi: 10.1371/journal.pbio.3003976.

[41] Joseph Matheson and Joanna Masel. “Background selection from unlinked sites causes nonindependent evolution of deleterious mutations”. In: Genome Biology and Evolution 16.3 (2024), evae050. doi: 10.1093/gbe/evae050.

[42] Anne Charmantier and Dany Garant. “Environmental quality and evolutionary potential: lessons from wild populations”. In: Proceedings of the Royal Society B: Biological Sciences 272.1571 (2005), pp. 1415–1425. doi: 10.1098/rspb.2005.3117.

[43] Alastair J. Wilson, Josephine M. Pemberton, Jill G. Pilkington, David W. Coltman, D. V. Mifsud, Tim H. Clutton-Brock, and Loeske E. B. Kruuk. “Environmental coupling of selection and heritability limits evolution”. In: PLoS Biology 4.7 (2006), e216. doi: 10.1371/journal.pbio.0040216.

[44] Walid Mawass, Francine M. Mayer, and Emmanuel Milot. “Genotype-by-environment interactions modulate the rate of microevolution in reproductive timing in humans”. In: Evolution 76.7 (2022), pp. 1391–1405. doi: 10.1111/evo.14504.

[45] Reinhard Bürger, Günter P. Wagner, and Franz Stettinger. “How much heritable variation can be maintained in finite populations by mutation-selection balance?” In: Evolution 43.8 (1989), pp. 1748–1766. doi: 10.1111/j.1558-5646.1989.tb02624.x.

[46] Motoo Kimura. “The number of heterozygous nucleotide sites maintained in a finite population due to steady flux of mutations”. In: Genetics 61.4 (1969), pp. 893–903. doi: 10.1093/genetics/61.4.893.

[47] Masatoshi Nei and Motoi Murata. “Effective population size when fertility is inherited”. In: Genetical Research 8.2 (1966), pp. 257–260. doi: 10.1017/S0016672300010119.

[48] Pierre de Villemereuil, Holger Schielzeth, Shinichi Nakagawa, and Michael Morrissey. “General methods for evolutionary quantitative genetic inference from generalized mixed models”. In: Genetics 204.3 (2016), pp. 1281–1294. doi: 10.1534/genetics.115.186536.

[49] Richard Frankham. “Effective population size/adult population size ratios in wildlife: a review”. In: Genetical Research 66.2 (1995), pp. 95–107. doi: 10.1017/S0016672300034455.

[50] Jacob I. Marsh, Austin T. Daigle, and Parul Johri. “The B-value calculator: expected diversity with background selection”. In: bioRxiv (2026). Preprint. doi: 10.64898/2026.03.04.709642.

[51] Charles R. Henderson. “Best linear unbiased estimation and prediction under a selection model”. In: Biometrics 31.2 (1975), pp. 423–447. doi: 10.2307/2529430.

[52] Alastair J. Wilson, Denis Réale, Michelle N. Clements, Michael B. Morrissey, Erik Postma, Craig A. Walling, Loeske E. B. Kruuk, and Daniel H. Nussey. “An ecologist’s guide to the animal model”. In: Journal of Animal Ecology 79.1 (2010), pp. 13–26. doi: 10.1111/j.1365-2656.2009.01639.x.

[53] Kees van Oers, Pieter Jan Drent, Niels Jeroen Dingemanse, and Bart Kempenaers. “Personality is associated with extrapair paternity in great tits, Parus major”. In: Animal Behaviour 76.3 (2008), pp. 555–563. doi: 10.1016/j.anbehav.2008.03.011.

[54] Shaohong Feng, Josefin Stiller, Yuan Deng, Joel Armstrong, Qi Fang, Andrew Hart Reeve, Duo Xie, Guangji Chen, Chunxue Guo, Brant C. Faircloth, et al. “Dense sampling of bird diversity increases power of comparative genomics”. In: Nature 587.7833 (2020), pp. 252–257. doi: 10.1038/s41586-020-2873-9.

[55] Joshua V. Peñalba, Yuan Deng, Qi Fang, Leo Joseph, Craig Moritz, and Andrew Cockburn. “Genome of an iconic Australian bird: High-quality assembly and linkage map of the superb fairy-wren (Malurus cyaneus)”. In: Molecular Ecology Resources 20.2 (2020), pp. 560–578. doi: 10.1111/1755-0998.13124.

[56] Linnéa Smeds, Anna Qvarnström, and Hans Ellegren. “Direct estimate of the rate of germline mutation in a bird”. In: Genome Research 26.9 (2016), pp. 1211–1218. doi: 10.1101/gr.204669.116.

[57] Richard J. Wang, Gregg W.C. Thomas, Muthuswamy Raveendran, R. Alan Harris, Harshavardhan Doddapaneni, Donna M. Muzny, John P. Capitanio, Predrag Radivojac, Jeffrey Rogers, and Matthew W. Hahn. “Paternal age in rhesus macaques is positively associated with germline mutation accumulation but not with measures of offspring sociability”. In: Genome Research 30.6 (2020), pp. 826–834. doi: 10.1101/gr.255174.119.

[58] Lucie A Bergeron, Søren Besenbacher, Jaco Bakker, Jiao Zheng, Panyi Li, George Pacheco, MikkelHolger S Sinding, Maria Kamilari, M Thomas P Gilbert, Mikkel H Schierup, et al. “The germline mutational process in rhesus macaque and its implications for phylogenetic dating”. In: GigaScience 10.5 (2021), giab029. doi: 10.1093/gigascience/giab029.

[59] Ellie E. Armstrong, Sarah B. Carey, Alex Harkess, Gabriele Zenato Lazzari, Katherine A. Solari, Jesuś E. Maldonado, Robert C. Fleischer, Neel Aziz, Patricia Walsh, Klaus-Peter Koepfli, et al. “Parameterizing Pantherinae: de novo mutation rate estimates from Panthera and Neofelis pedigrees”. In: Genome Biology and Evolution 17.4 (2025), evaf060. doi: 10.1093/gbe/evaf060.

[60] Yiguan Wang and Darren J. Obbard. “Experimental estimates of germline mutation rate in eukaryotes: a phylogenetic meta-analysis”. In: Evolution Letters 7.4 (2023), pp. 216–226. doi: 10.1093/evlett/qrad027.

[61] Bernard Y. Kim, Christian D. Huber, and Kirk E. Lohmueller. “Inference of the distribution of selection coefficients for new nonsynonymous mutations using large samples”. In: Genetics 206.1 (2017), pp. 345–361. doi: 10.1534/genetics.116.197145.

[62] Jeffrey Rogers, Michael C. Mahaney, Shelly M. Witte, Shalini Nair, Deborah Newman, Steven Wedel, Lawrence A. Rodriguez, Karen S. Rice, Susan H. Slifer, Andrey Perelygin, et al. “A Genetic Linkage Map of the Baboon (Papio hamadryas) Genome Based on Human Microsatellite Polymorphisms”. In: Genomics 67.3 (2000), pp. 237–247. doi: 10.1006/geno.2000.6245.

[63] Laura A. Cox, Michael C. Mahaney, John L. VandeBerg, and Jeffrey Rogers. “A second-generation genetic linkage map of the baboon (Papio hamadryas) genome”. In: Genomics 88.3 (2006), pp. 274–281. doi: 10.1016/j.ygeno.2006.03.020.

[64] Cyril J Versoza, Sarah Weiss, Ravneet Johal, Bruno La Rosa, Jeffrey D Jensen, and Susanne P Pfeifer. “Novel Insights into the Landscape of Crossover and Noncrossover Events in Rhesus Macaques (Macaca mulatta)”. In: Genome Biology and Evolution 16.1 (2024), evad223. doi: 10.1093/gbe/evad223.

[65] Veronika N. Laine, Toni I. Gossmann, Kyle M. Schachtschneider, Colin J. Garroway, Ole Madsen, Koen J. F. Verhoeven, Victor de Jager, Hendrik-Jan Megens, Wesley C. Warren, Patrick Minx, et al. “Evolutionary signals of selection on cognition from the great tit genome and methylome”. In: Nature Communications 7.1 (2016), p. 10474. doi: 10.1038/ncomms10474.

[66] B Hansson, M Ljungqvist, D A Dawson, J C Mueller, J Olano-Marin, H Ellegren, and J-Å Nilsson. “Avian genome evolution: insights from a linkage map of the blue tit (Cyanistes caeruleus)”. In: Heredity 104.1 (2010), pp. 67–78. doi: 10.1038/hdy.2009.107.

[67] Hui Zhen Tan, Phoebe Scherer, Katarina C. Stuart, Sarah Bailey, Kate D. Lee, Patricia Brekke, John G. Ewen, Annabel Whibley, and Anna W. Santure. “A high-density linkage map reveals broadand fine-scale sex differences in recombination in the hihi (stitchbird; Notiomystis cincta)”. In: Heredity 133.4 (2024), pp. 262–275. doi: 10.1038/s41437-024-00711-3.

[68] Russell B. Corbett-Detig, Daniel L. Hartl, and Timothy B. Sackton. “Natural Selection Constrains Neutral Diversity across A Wide Range of Species”. In: PLOS Biology 13.4 (2015), e1002112. doi: 10.1371/journal.pbio.1002112.

[69] Cheng Xue, Muthuswamy Raveendran, R. Alan Harris, Gloria L. Fawcett, Xiaoming Liu, Simon White, Mahmoud Dahdouli, David Rio Deiros, Jennifer E. Below, William Salerno, et al. “The population genomics of rhesus macaques (Macaca mulatta) based on whole-genome sequences”. In: Genome Research 26.12 (2016), pp. 1651–1662. doi: 10.1101/gr.204255.116.

[70] Ludovic Dutoit, Reto Burri, Alexander Nater, Carina F. Mugal, and Hans Ellegren. “Genomic distribution and estimation of nucleotide diversity in natural populations: perspectives from the collared flycatcher (Ficedula albicollis) genome”. In: Molecular Ecology Resources 17.4 (2017), pp. 586–597. doi: 10.1111/1755-0998.12602.

[71] Jun Chen, Sylvain Glémin, and Martin Lascoux. “Genetic Diversity and the Efficacy of Purifying Selection across Plant and Animal Species”. In: Molecular Biology and Evolution 34.6 (2017), pp. 1417–1428. doi: 10.1093/molbev/msx088.

[72] Henry J. Barton and Kai Zeng. “The impact of natural selection on short insertion and deletion variation in the great tit genome”. In: Genome Biology and Evolution 11.6 (2019), pp. 1514–1524. doi: 10.1093/gbe/evz068.

[73] Dongfeng Wang, Hosein Salehian-Dehkordi, Langda Suo, and Fenghua Lv. “Impacts of population size and domestication process on genetic diversity and genetic load in genus Ovis”. In: Genes 14.10 (2023), p. 1977. doi: 10.3390/genes14101977.

[74] Katherine Carbeck, Peter Arcese, Irby Lovette, Christin Pruett, Kevin Winker, and Jennifer Walsh. “Candidate genes under selection in song sparrows co-vary with climate and body mass in support of Bergmann’s rule”. In: Nature Communications 14.1 (2023), p. 6974. doi: 10.1038/s41467-023-42786-2.

[75] Nathan P. Myhrvold, Elita Baldridge, Benjamin Chan, Dhileep Sivam, Daniel L. Freeman, and S. K. Morgan Ernest. “An amniote life-history database to perform comparative analyses with birds, mammals, and reptiles”. In: Ecology 96.11 (2015), pp. 3109–3109. doi: 10.1890/15-0846r.1.

[76] John Damuth. “Interspecific allometry of population density in mammals and other animals: the independence of body mass and population energy-use”. In: Biological Journal of the Linnean Society 31.3 (1987), pp. 193–246. doi: 10.1111/j.1095-8312.1987.tb01990.x.

[77] Charles Perrier, Quentin Rougemont, and Anne Charmantier. “Demographic history and genomics of local adaptation in blue tit populations”. In: Evolutionary Applications 13.6 (2020), pp. 1145–1165. doi: 10.1111/eva.13035.

[78] Chenlu Di, Swetha Ramesh, Jason Ernst, and Kirk E. Lohmueller. “The landscape of fitness effects of putatively functional noncoding mutations in humans”. In: bioRxiv (2025). doi: 10.1101/2025.05.14.654124.

[79] John R. Pannell and Brian Charlesworth. “Effects of metapopulation processes on measures of genetic diversity”. In: Philosophical Transactions of the Royal Society of London. Series B: Biological Sciences 355.1404 (2000), pp. 1851–1864. doi: 10.1098/rstb.2000.0740.

[80] Michael C. Whitlock and Nicholas H. Barton. “The effective size of a subdivided population”. In: Genetics 146.1 (1997), pp. 427–441. doi: 10.1093/genetics/146.1.427.

[81] Graham McVicker, David Gordon, Colleen Davis, and Phil Green. “Widespread genomic signatures of natural selection in hominid evolution”. In: PLoS Genetics 5.5 (2009), e1000471. doi: 10.1371/journal.pgen.1000471.

[82] Xinyi Li and Jeremy J. Berg. “Background selection in recombining genomes and its consequences for the maintenance of variation in complex traits”. In: Proceedings of the National Academy of Sciences 123.14 (2026), e2513613123. doi: 10.1073/pnas.2513613123.

[83] Alexey S. Kondrashov and James F. Crow. “A molecular approach to estimating the human deleterious mutation rate”. In: Human Mutation 2.3 (1993), pp. 229–234. doi: 10.1002/humu.1380020312.

[84] Joseph Matheson, Ulises Hernández, Jason Bertram, and Joanna Masel. “Human deleterious mutation rate slows adaptation and implies high fitness variance”. In: bioRxiv (2023). doi: 10.1101/2023.09.01.555871.

[85] Ulises Hernández, Walid Mawass, Joseph Matheson, and Joanna Masel. “Rare variants drive high variance in human ancestral fitness at mutation-selection-drift balance”. In: bioRxiv (2026). doi: 10.64898/2026.07.29.741368.

[86] Michael B. Morrissey, Loeske E. B. Kruuk, and Alastair J. Wilson. “The danger of applying the breeder’s equation in observational studies of natural populations”. In: Journal of Evolutionary Biology 23.11 (2010), pp. 2277–2288. doi: 10.1111/j.1420-9101.2010.02084.x.

[87] Pierre de Villemereuil, Anne Charmantier, Debora Arlt, Pierre Bize, Patricia Brekke, Lyanne Brouwer, Andrew Cockburn, Steeve D. Côté, F. Stephen Dobson, Simon R. Evans, et al. “Fluctuating optimum and temporally variable selection on breeding date in birds and mammals”. In: Proceedings of the National Academy of Sciences 117.50 (2020), pp. 31969–31978. doi: 10.1073/pnas.2009003117.

[88] Reinhard Bürger and Michael Lynch. “Evolution and extinction in a changing environment: a quantitative-genetic analysis”. In: Evolution 49.1 (1995), pp. 151–163. doi: 10.1111/j.1558-5646.1995.tb05967.x.

[89] Russell Lande and Susan Shannon. “The role of genetic variation in adaptation and population persistence in a changing environment”. In: Evolution 50.1 (1996), pp. 434–437. doi: 10.1111/j.1558-5646.1996.tb04504.x.

[90] Jason Bertram and Zahra Shafiei. “Strong amplification of quantitative genetic variation under a balance between mutation and fluctuating stabilizing selection”. In: Genetics 233.1 (2026), iyag063. doi: 10.1093/genetics/iyag063.

[91] Thomas Lenormand. “Gene flow and the limits to natural selection”. In: Trends in Ecology & Evolution 17.4 (2002), pp. 183–189. doi: 10.1016/S0169-5347(02)02497-7.

[92] Anne Charmantier, Claire Doutrelant, Gabrielle Dubuc-Messier, Amélie Fargevieille, and Marta Szulkin. “Mediterranean blue tits as a case study of local adaptation”. In: Evolutionary Applications 9.1 (2016), pp. 135–152. doi: 10.1111/eva.12282.

[93] Austin Burt. “The evolution of fitness”. In: Evolution 49.1 (1995), pp. 1–8. doi: 10.1111/j.1558-5646.1995.tb05954.x.

[94] Andrew P. Hendry, Daniel J. Schoen, Matthew E. Wolak, and Jane M. Reid. “The contemporary evolution of fitness”. In: Annual Review of Ecology, Evolution, and Systematics 49.1 (2018), pp. 457–476. doi: 10.1146/annurev-ecolsys-110617-062358.

[95] Loeske E. B. Kruuk and Jarrod D. Hadfield. “How to separate genetic and environmental causes of similarity between relatives”. In: Journal of Evolutionary Biology 20.5 (2007), pp. 1890–1903. doi: 10.1111/j.1420-9101.2007.01377.x.

[96] Katie V. Stopher, Craig A. Walling, Alison Morris, Fiona E. Guinness, Tim H. Clutton-Brock, Josephine M. Pemberton, and Daniel H. Nussey. “Shared spatial effects on quantitative genetic parameters: accounting for spatial autocorrelation and home range overlap reduces estimates of heritability in wild red deer”. In: Evolution 66.8 (2012), pp. 2411–2426. doi: 10.1111/j.1558-5646.2012.01620.x.

[97] Amelie Baud, Sarah McPeek, Nancy Chen, and Kimberly A. Hughes. “Indirect genetic effects: a cross-disciplinary perspective on empirical studies”. In: Journal of Heredity 113.1 (2022), pp. 1–15. doi: 10.1093/jhered/esab059.

[98] Walid Mawass and Emmanuel Milot. “Assessing the impact of pedigree attributes on the validity of quantitative genetic parameter estimates”. In: Journal of Evolutionary Biology 38.4 (2025), pp. 439–456. doi: 10.1093/jeb/voaf010.

[99] Steinar Engen, Russell Lande, and Bernt-Erik Sæther. “Effective size of a fluctuating age-structured population”. In: Genetics 170.2 (2005), pp. 941–954. doi: 10.1534/genetics.104.028233.

[100] Bjarki Eldon and John Wakeley. “Coalescent processes when the distribution of offspring number among individuals is highly skewed”. In: Genetics 172.4 (2006), pp. 2621–2633. doi: 10.1534/genetics.105.052175.

[101] Richard A. Neher and Oskar Hallatschek. “Genealogies of rapidly adapting populations”. In: Proceedings of the National Academy of Sciences 110.2 (2013), pp. 437–442. doi: 10.1073/pnas.1213113110.

[102] Dimitrios Diamantidis, Wai-Tong Fan, Matthias Birkner, and John Wakeley. “Bursts of coalescence within population pedigrees whenever big families occur”. In: Genetics 227.1 (2024), iyae030. doi: 10.1093/genetics/iyae030.

[103] Meike J. Wittmann, Sylvain Mousset, and Joachim Hermisson. “Modeling the genetic footprint of fluctuating balancing selection: from the local to the genomic scale”. In: Genetics 223.4 (2023), iyad022. doi: 10.1093/genetics/iyad022.

[104] Olivia L. Johnson, Raymond Tobler, Joshua M. Schmidt, and Christian D. Huber. “Seasonal fluctuations in fitness result in severe reductions in effective population size”. In: G3: Genes, Genomes, Genetics 16.11 (2026), jkag252. doi: 10.1093/g3journal/jkag252.

[105] Manas Geeta Arun, Aidan Angus-Henry, Darren J. Obbard, and Jarrod D. Hadfield. “Estimating the additive genetic variance for relative fitness from changes in allele frequency”. In: Genetics 232.1 (2026), iyaf232. doi: 10.1093/genetics/iyaf232.

[106] Nancy Chen, Ivan Juric, Elissa J. Cosgrove, Reed Bowman, John W. Fitzpatrick, Stephan J. Schoech, Andrew G. Clark, and Graham Coop. “Allele frequency dynamics in a pedigreed natural population”. In: Proceedings of the National Academy of Sciences 116.6 (2019), pp. 2158–2164. doi: 10.1073/pnas.1813852116.

[107] Jarrod D. Hadfield. “MCMC Methods for Multi-Response Generalized Linear Mixed Models: The MCMCglmm R Package”. In: Journal of Statistical Software 33.2 (2010). doi: 10.18637/jss.v033.i02.

[108] Michael Lynch and William G. Hill. “Phenotypic evolution by neutral mutation”. In: Evolution 40.5 (1986), pp. 915–935. doi: 10.1111/j.1558-5646.1986.tb00561.x.

[109] Armando Caballero. “Developments in the prediction of effective population size”. In: Heredity 73.6 (1994), pp. 657–679. doi: 10.1038/hdy.1994.174.

[110] James F. Crow and Carter Denniston. “Inbreeding and variance effective population numbers”. In: Evolution 42.3 (1988), pp. 482–495. doi: 10.1111/j.1558-5646.1988.tb04154.x.

[111] Motoo Kimura. “On the probability of fixation of mutant genes in a population”. In: Genetics 47.6 (1962), pp. 713–719. doi: 10.1093/genetics/47.6.713.

[112] Jinliang Wang, Enrique Santiago, and Armando Caballero. “Prediction and estimation of effective population size”. In: Heredity 117.4 (2016), pp. 193–206. doi: 10.1038/hdy.2016.43.

[113] Nicholas H. Barton. “The effect of hitch-hiking on neutral genealogies”. In: Genetical Research 72.2 (1998), pp. 123–133. doi: 10.1017/S0016672398003462.

[114] Enrique Santiago and Armando Caballero. “Joint prediction of the effective population size and the rate of fixation of deleterious mutations”. In: Genetics 204.3 (2016), pp. 1267–1279. doi: 10.1534/genetics.116.188250.

[115] Nicholas H. Barton and Peter D. Keightley. “Understanding quantitative genetic variation”. In: Nature Reviews Genetics 3.1 (2002), pp. 11–21. doi: 10.1038/nrg700.

[116] Mark Kirkpatrick and Nick Barton. “Chromosome inversions, local adaptation and speciation”. In: Genetics 173.1 (2006), pp. 419–434. doi: 10.1534/genetics.105.047985.

[117] Sam Yeaman. “Genomic rearrangements and the evolution of clusters of locally adaptive loci”. In: Proceedings of the National Academy of Sciences 110.19 (2013), E1743–E1751. doi: 10.1073/pnas.1219381110.

[118] Anne Charmantier and Jacques Blondel. “A contrast in extra-pair paternity levels on mainland and island populations of Mediterranean blue tits”. In: Ethology 109.4 (2003), pp. 351–363. doi: 10.1046/j.1439-0310.2003.00880.x.

[119] Anne Charmantier, Jacques Blondel, Philippe Perret, and Marcel M. Lambrechts. “Do extra-pair paternities provide genetic benefits for female blue tits Parus caeruleus?” In: Journal of Avian Biology 35.6 (2004), pp. 524–532. doi: 10.1111/j.0908-8857.2004.03296.x.

[120] Takeshi Kawakami, Linnéa Smeds, Niclas Backström, Arild Husby, Anna Qvarnström, Carina F. Mugal, Pall Olason, and Hans Ellegren. “A high-density linkage map enables a second-generation collared flycatcher genome assembly and reveals the patterns of avian recombination rate variation and chromosomal evolution”. In: Molecular Ecology 23.16 (2014), pp. 4035–4058. doi: 10.1111/mec.12810.

[121] Ben C. Sheldon and Hans Ellegren. “Sexual selection resulting from extrapair paternity in collared flycatchers”. In: Animal Behaviour 57.2 (1999), pp. 285–298. doi: 10.1006/anbe.1998.0968.

[122] K. van Oers, A. W. Santure, I. De Cauwer, N. E. M. van Bers, R. P. M. A. Crooijmans, B. C. Sheldon, M. E. Visser, J. Slate, and M. A. M. Groenen. “Replicated high-density genetic maps of two great tit populations reveal fine-scale genomic departures from sex-equal recombination rates”. In: Heredity 112.3 (2014), pp. 307–316. doi: 10.1038/hdy.2013.107.

[123] Samantha C. Patrick, Joanne R. Chapman, Hannah L. Dugdale, John L. Quinn, and Ben C. Sheldon. “Promiscuity, paternity and personality in the great tit”. In: Proceedings of the Royal Society B: Biological Sciences 279.1734 (2012), pp. 1724–1730. doi: 10.1098/rspb.2011.1820.

[124] Nina Bircher, Kees van Oers, Camilla A. Hinde, and Marc Naguib. “Extraterritorial forays by great tits are associated with dawn song in unexpected ways”. In: Behavioral Ecology 31.4 (2020), pp. 873–883. doi: 10.1093/beheco/araa040.

[125] Pirmin Nietlisbach, Glauco Camenisch, Thomas Bucher, Jon Slate, Lukas F. Keller, and Erik Postma. “A microsatellite-based linkage map for song sparrows (Melospiza melodia)”. In: Molecular Ecology Resources 15.6 (2015), pp. 1486–1496. doi: 10.1111/1755-0998.12414.

[126] Rebecca J. Sardell, Lukas F. Keller, Peter Arcese, Thomas Bucher, and Jane M. Reid. “Comprehensive paternity assignment: genotype, spatial location and social status in song sparrows, Melospiza melodia”. In: Molecular Ecology 19.19 (2010), pp. 4352–4364. doi: 10.1111/j.1365-294X.2010.04805.x.

[127] Gabriela K. Hajduk, Andrew Cockburn, Nicolas Margraf, Helen L. Osmond, Craig A. Walling, and Loeske E. B. Kruuk. “Inbreeding, inbreeding depression, and infidelity in a cooperatively breeding bird”. In: Evolution 72.7 (2018), pp. 1500–1514. doi: 10.1111/evo.13496.

[128] Fay Morland, John G. Ewen, Anna W. Santure, Patricia Brekke, and Nicola Hemmings. “Demographic drivers of reproductive failure in a threatened bird: insights from a decade of data”. In: Proceedings of the National Academy of Sciences 121.36 (2024), e2319104121. doi: 10.1073/pnas.2319104121.

[129] Josephine Pemberton, Susan E. Johnston, T. John Fletcher, Darwin Tree of Life Barcoding collective, Wellcome Sanger Institute Tree of Life programme, Wellcome Sanger Institute Scientific Operations: DNA Pipelines collective, Tree of Life Core Informatics collective, and Darwin Tree of Life Consortium. “The genome sequence of the red deer, Cervus elaphus Linnaeus 1758”. In: Wellcome Open Research 6 (2021), p. 336. doi: 10.12688/wellcomeopenres.17493.1.

[130] Susan E Johnston, Jisca Huisman, Philip A Ellis, and Josephine M Pemberton. “A High-Density Link-age Map Reveals Sexual Dimorphism in Recombination Landscapes in Red Deer (Cervus elaphus)”. In: G3: Genes, Genomes, Genetics 7.8 (2017), pp. 2859–2870. doi: 10.1534/g3.117.044198.

[131] Susan E Johnston, Camillo Bérénos, Jon Slate, and Josephine M Pemberton. “Conserved Genetic Architecture Underlying Individual Recombination Rate Variation in a Wild Population of Soay Sheep (Ovis aries)”. In: Genetics 203.1 (2016), pp. 583–598. doi: 10.1534/genetics.115.185553.

[132] Jocelyn Poissant, John T Hogg, Corey S Davis, Joshua M Miller, Jillian F Maddox, and David W Coltman. “Genetic linkage map of a wild genome: genomic structure, recombination and sexual dimorphism in bighorn sheep”. In: BMC Genomics 11.1 (2010), p. 524. doi: 10.1186/1471-2164-11-524.

[133] Sanjit Singh Batra, Michal Levy-Sakin, Jacqueline Robinson, Joseph Guillory, Steffen Durinck, Tauras P Vilgalys, Pui-Yan Kwok, Laura A Cox, Somasekar Seshagiri, Yun S Song, et al. “Accurate assembly of the olive baboon (Papio anubis) genome using long-read and Hi-C data”. In: GigaScience 9.12 (2020), giaa134. doi: 10.1093/gigascience/giaa134.

[134] Felix L. Wu, Alva I. Strand, Laura A. Cox, Carole Ober, Jeffrey D. Wall, Priya Moorjani, and Molly Przeworski. “A comparison of humans and baboons suggests germline mutation rates do not track cell divisions”. In: PLOS Biology 18.8 (2020), e3000838. doi: 10.1371/journal.pbio.3000838.

[135] Chentao Yang, Fang Li, Zijun Xiong, Klaus-Peter Koepfli, Oliver Ryder, Polina Perelman, Qiye Li, and Guojie Zhang. “A draft genome assembly of spotted hyena, Crocuta crocuta”. In: Scientific Data 7.1 (2020), p. 126. doi: 10.1038/s41597-020-0468-9.

[136] Olga Dudchenko, Sanjit S. Batra, Arina D. Omer, Sarah K. Nyquist, Marie Hoeger, Neva C. Durand, Muhammad S. Shamim, Ido Machol, Eric S. Lander, Aviva Presser Aiden, et al. “De novo assembly of the Aedes aegypti genome using Hi-C yields chromosome-length scaffolds”. In: Science 356.6333 (2017), pp. 92–95. doi: 10.1126/science.aal3327.

[137] Annika Herdtle, Chris Duncan, Marta B. Manser, and Tim Clutton-Brock. “Patterns and drivers of female extra-pair mating in wild Kalahari meerkats”. In: Behavioral Ecology 36.3 (2025), araf016. doi: 10.1093/beheco/araf016.

[138] Franc Jaňzekovič, Boris Kryštufek, Wellcome Sanger Institute Tree of Life Management, Samples and Laboratory team, Wellcome Sanger Institute Scientific Operations: Sequencing Operations, Wellcome Sanger Institute Tree of Life Core Informatics team, and Tree of Life Core Informatics collective. “The genome sequence of the European Snow Vole, Chionomys nivalis (Martins, 1842)”. In: Wellcome Open Research 10 (2025), p. 156. doi: 10.12688/wellcomeopenres.23922.1.

